# Evaluating ARG-estimation methods in the context of estimating population-mean polygenic score histories

**DOI:** 10.1101/2024.05.24.595829

**Authors:** Dandan Peng, Obadiah J. Mulder, Michael D. Edge

## Abstract

Scalable methods for estimating marginal coalescent trees across the genome present new opportunities for studying evolution and have generated considerable excitement, with new methods extending scalability to thousands of samples. Benchmarking of the available methods has revealed general tradeoffs between accuracy and scalability, but performance in downstream applications has not always been easily predictable from general performance measures, suggesting that specific features of the ARG may be important for specific downstream applications of estimated ARGs. To exemplify this point, we benchmark ARG estimation methods with respect to a specific set of methods for estimating the historical time course of a population-mean polygenic score (PGS) using the marginal coalescent trees encoded by the ancestral recombination graph (ARG). Here we examine the performance in simulation of seven ARG estimation methods: `ARGweaver, RENT+, Relate, tsinfer+tsdate, ARG-Needle, ASMC-clust`, and `SINGER`, using their estimated coalescent trees and examining bias, mean squared error (MSE), confidence interval coverage, and Type I and II error rates of the downstream methods. Although it does not scale to the sample sizes attainable by other new methods, `SINGER` produced the most accurate estimated PGS histories in many instances, even when `Relate, tsinfer+tsdate, ARG-Needle` and `ASMC-clust` used samples ten or more times as large as those used by `SINGER`. In general, the best choice of method depends on the number of samples available and the historical time period of interest. In particular, the unprecedented sample sizes allowed by `Relate, tsinfer+tsdate, ARG-Needle`, and `ASMC-clust` are of greatest importance when the recent past is of interest—further back in time, most of the tree has coalesced, and differences in contemporary sample size are less salient.

## 1 Introduction

The ancestral recombination graph, or ARG (Griffiths & Marjoram, 1996), is a rich representation of the history of a sample of haplotypes, including all the mutation, recombination, and common-ancestry events that affect contemporary variation. Thus, the ARG encodes all historical information that can be extracted from a sample of contemporary genomes, much of it in gene genealogies or coalescent trees (R. Hudson, 1990; Wakeley, 2016) for every location in the genome, termed “local” or “marginal” trees. The true ARG is generally unknown, and estimation of the ARG is a very challenging problem. Nonetheless, the last ten years have witnessed major advances in ARG estimation, with new methods that provide estimated ARGs with unprecedented accuracy, scalability, or both (Deng et al., 2024; Kelleher et al., 2019; Mirzaei & Wu, 2017; Rasmussen et al., 2014; Speidel et al., 2019; Zhang et al., 2023). These advances have produced a great deal of excitement about the potential of estimated ARGs in evolutionary biology and beyond (Brandt et al., 2024; Harris, 2019, 2023; Lewanski et al., 2024; Nielsen et al., 2025; Wong et al., 2024).

The promise of estimated ARGs depends on their performance in downstream applications. Many of the available ARG estimation methods have been benchmarked in general terms (Brandt et al., 2022; Deng et al., 2024), revealing tradeoffs between accuracy and scalability. (We refer to all the methods we consider here as “ARG estimation” methods, regardless of whether they estimate the full ARG, including recombinations between marginal trees, or just a set of marginal trees.) However, early indications are that the actual performance of ARG estimators in downstream applications can vary in ways that are not necessarily predicted by a generic accuracy vs. scalability tradeoff (Fan et al., 2022, 2023). Thus, it seems as though the performance of estimated ARGs in downstream procedures may depend on specific features of the ARG and how well they are estimated. To explore this point in depth, we conducted thorough benchmarking of ARG estimators with respect to a set of methods for studying polygenic traits. These methods take estimated marginal trees as input. Polygenic traits—traits influenced by genetic variants from across the genome—are a promising area for applications of estimated ARGs (Christ et al., 2024; Edge & Coop, 2019; Gunnarsson et al., 2024; Link et al., 2023; Stern et al., 2021b; Wang et al., 2024; Zhang et al., 2023; Zhu et al., 2024). Understanding the evolution of polygenic traits is challenging in part because signals of selection are spread across many loci. Researchers can study the history of polygenic traits by examining fossil records (Kappelman, 1996) or ancient DNA (Mathieson et al., 2015) where available. However, many traits do not leave a fossil record, and ancient DNA is not always available. An alternative approach is to examine the genomes of contemporary individuals, perhaps in combination with the estimated effects of contemporary variants on target traits (Berg & Coop, 2014; Field et al., 2016; Racimo et al., 2018; M. R. Robinson et al., 2015; Uricchio et al., 2019), examining either allele-frequency differences among groups or traces of selection in patterns of within-population genetic diversity. If allele-frequency changes can be estimated from contemporary genetic data, then those allele-frequency changes, in combination with information about associations between alleles and traits of interest, can be used to study selection on traits. One reason estimated ARGs may be useful in population-genetic analysis of such subtle signals is that, to the extent the estimated ARG is correct, it naturally integrates information from flanking genomic regions in a way that reflects the history of recombination near the locus (Link et al., 2023). Relatedly, others have noted that even if estimated tree sequences are incorrect, the fact that they integrate information from nearby segments can prove useful in downstream inference (Whitehouse et al., 2024).

Edge & Coop (2019) proposed a set of methods to estimate the historical time course of a predicted population-mean trait level using local coalescent trees embedded in an ARG. The trait prediction is known as a polygenic score (PGS) or polygenic index. Although some of their methods are applicable to any trait prediction formed from genetic data, Edge & Coop focused on a population-mean PGS expressed as a weighted sum of population allele frequencies at unlinked loci, 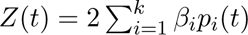, where the weight *β_i_* is the additive effect size on a trait of interest of an allele at locus *i*, and *p_i_*(*t*) is the frequency of the effect allele at locus *i* at time *t*. The estimators operate by estimating allele-frequency changes at the loci contributing to the PGS. Under neutrality, the proportion of lineages at time *t* in a coalescent tree subtending derived alleles in the contemporary sample is an unbiased estimator of the derived allele frequency at time *t*. Under selection, the ancestors of the sample are a biased sample from the ancestral population, but ideas from phylodynamics can be borrowed to form noisy estimates of the number of carriers of each allelic type in a specified time period.

There are other ARG-based methods for identifying selection. In particular, Relate (Speidel et al., 2019) computes a statistic designed to detect selection on a single tree. `CLUES` (Stern et al., 2019) uses a full-likelihood approach to identify loci under selection integrating over posterior samples of a local tree. `SIA` (Hejase et al., 2022) trains a neural network to identify trees that appear to show evidence of selection. All of these methods are for individual genetic loci. `PALM` (Stern et al., 2021a) considers evidence for selection on polygenic traits, identifying a selection gradient that maximizes the likelihood of local trees across sites that are associated with a polygenic trait. The estimators of Edge & Coop that we consider here work with polygenic traits, like `PALM`, but they aim to estimate the historical time course of a population-level polygenic score rather than a selection gradient for a trait.

To gain a deeper understanding of the performance of different ARG estimation approaches in estimating population-mean PGS histories, we applied the Edge & Coop framework to estimate local trees from seven methods for ARG estimation. For comparison with the work of Edge & Coop (2019), we evaluated `RENT+` (Mirzaei & Wu, 2017), which they used in their original paper. We also evaluated the performance of `ARGweaver` (Rasmussen et al., 2014), which predates `RENT+` and provides more accurate estimates on smaller samples, as well as Relate (Speidel et al., 2019), `tsinfer+tsdate` (Kelleher et al., 2019; Wohns et al., 2022), `ARG-Needle` and `ASMC-clust` (Zhang et al., 2023), and SINGER (Deng et al., 2024), all of which scale to larger samples. We also consider the effect of sample size on the resulting estimates.

## 2 Methods

We simulated derived-allele frequency trajectories at all unlinked loci that affect a trait. The true population-mean PGS trajectory was computed as the weighted sum 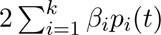 of these allele-frequency trajectories, where the weights *β_i_* are additive effect sizes and *p_i_*(*t*) is the frequency of the effect allele at locus *i* at time *t*. Taking these simulated allele-frequency trajectories as input, we generated corresponding coalescent trees and haplotypes using `mssel` (Berg & Coop, 2015). The haplotypes were fed to all of the tree estimation software packages considered here, and the estimated trees were saved. Upon obtaining both the true and estimated trees, we applied the allele-frequency estimators from Edge & Coop (2019) to them, and the estimated allele-frequency trajectories were used to compute estimated population-mean PGS trajectories using the additive effect sizes. The estimated population-mean PGS trajectory was then compared with the true trajectory using various metrics (Fig. 1).

**Figure 1:**
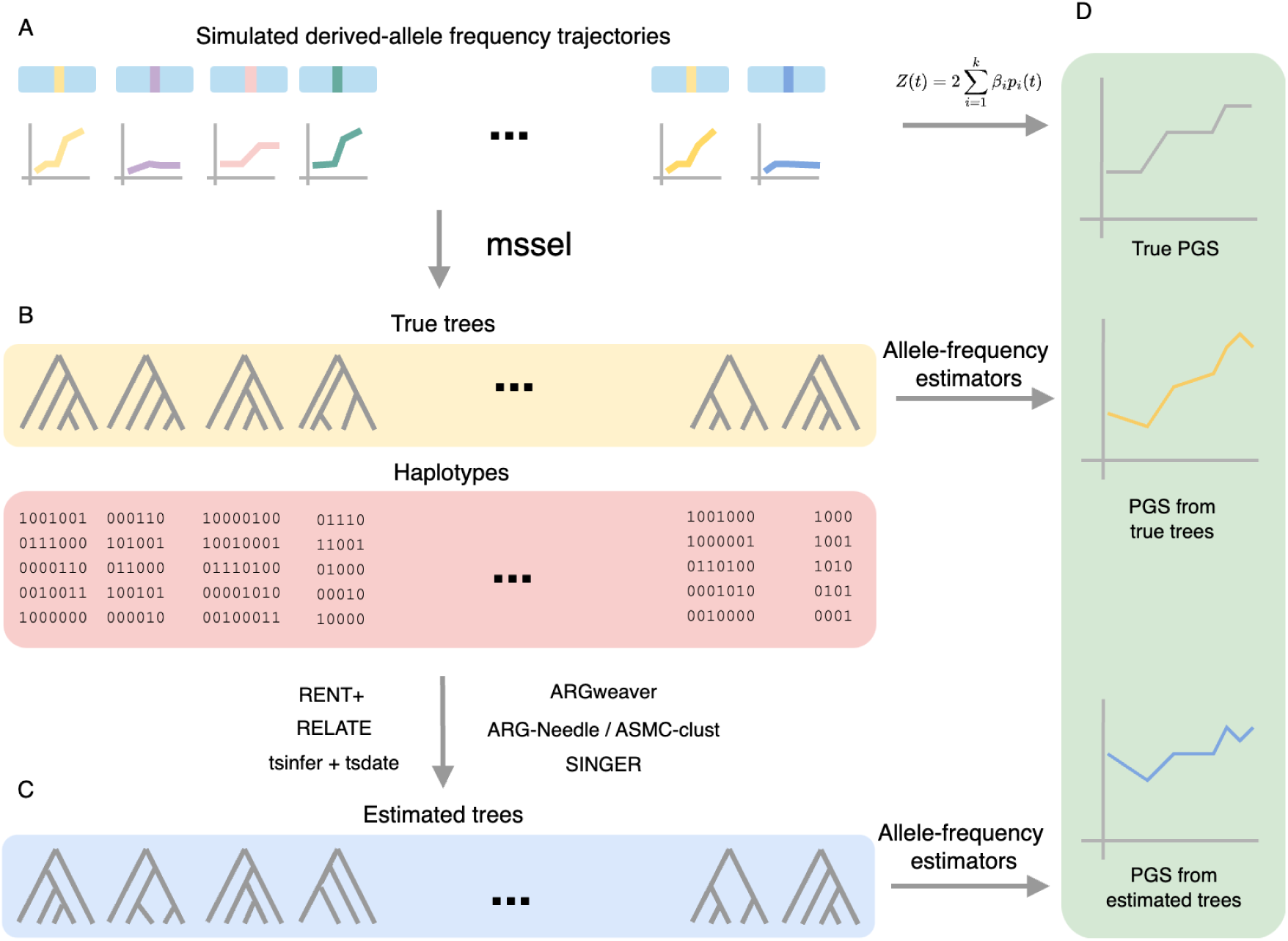
Method overview. A) Simulate derived allele frequency trajectories at unlinked loci. B) Generate true trees and haplotypes from those trajectories. C) Obtain estimated trees from tree estimation software. D) Apply allele-frequency estimators to true and estimated trees, then compare the true and estimated population-mean PGS trajectories.

### 2.1 Simulations

We simulated the population-mean PGS trajectories of traits additively determined by 100 unlinked loci under two scenarios: (i) neutral evolution, and (ii) trait-increasing directional selection occurring from 0.04 to 0.02 coalescent units ago, with neutrality at other time points. Assuming an effective population size of 10,000 and a generation time of 30 years, the period of 0.02-0.04 coalescent units in the past corresponds to 12,000-24,000 years in the past, or the most recent part of the Upper Paleolithic. Because the bulk of ancient DNA evidence is from samples more recent than this (Mathieson et al., 2015; Speidel et al., 2021; Stern et al., 2021b), indirect methods to detect selection in humans are perhaps especially of interest in this epoch, given that allele-frequency changes cannot as easily be examined directly (Le et al., 2022; Mathieson & Terhorst, 2022). The PGS is a weighted sum of the allele frequencies, with weights equal to the additive effect sizes. There are no environmental effects or interactions—in our simulations, there is no distinction between the PGS and the trait. Further, we treat all the effect sizes as known.

For each trait, we simulated the derived allele-frequency histories of 100 trait-associated loci. For each locus, an effect size for the derived allele was drawn from a normal distribution N(0, *h*^2^*σ*^2^*w/n*) in which the heritability *h*^2^ and the contemporary variance *σ*^2^ of the trait are set to 1, *w* is a modified version of Watterson’s constant calculated as 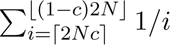 (*N* is the effective population size; *c* is the minimum minor allele frequency being drawn, we set *c* = 0.01), and *n* is the number of loci affecting the trait, set to 100. We consider a hypothetical trait with a heritability of 1 and for which “true” additive effect sizes are known. We do this because we are interested in considering estimation accuracy at the level of a population-mean polygenic score rather than the trait to which the polygenic score corresponds. Polygenic scores will differ from trait values due to biases and errors in GWAS estimation and SNP heritabilities less than one, among other factors. Further, back in time, systematic changes in the environment and difficulties in “porting” polygenic scores trained in modern samples onto ancient samples will affect accuracy. Thus, we limit our focus to accuracy in population-mean polygenic score estimation. However, we emphasize that the model for selection we use assumes that selection occurs on the polygenic score itself. Given that most polygenic scores are noisy predictors of the traits with which they are associated, the selection gradients we simulate might be thought of as corresponding to larger selection gradients on actual trait values.

Next, we simulated allele-frequency trajectories for each locus. For both neutral and selected traits, the majority of the history of each locus was simulated backward in time—for neutral traits, simulation was entirely backward, and for selected traits, simulation was backward-in-time prior and subsequent to the period in which selection occurred. The simulation is forward-in-time during the selection interval. We set the probability of obtaining a derived-allele frequency *k/*2*N* at the more recent end of the period simulated backward in time to be inversely proportional to *k*, where *k* ∈ {1, …, 2*N* − 1}. This sampled allele frequency serves as the present derived-allele frequency under neutrality and as the initial allele frequency at the onset of the selection in the recent-selection scenario.

Given a derived-allele frequency *p_i_*(*t*), the allele frequency at 1/(2*N*) coalescent time units in the past was drawn from N(*p_i_*(*t*)(2 − 1/2*N*), *p_i_*(*t*)(1 − *p_i_*(*t*))/2*N*) (Berg & Coop, 2015; Lee & Coop, 2017; Przeworski et al., 2005). During the selection period, we simulated the derived-allele frequency forward in time in steps of 1/(2*N*) coalescent units. The frequency *p_i_*(*t*+1) was drawn from N(*p_i_*(*t*)+*sp_i_*(*t*)(1−*p_i_*(*t*)), *p_i_*(*t*)(1−*p_i_*(*t*))/2*N*) conditional on the frequency *p_i_*(*t*), where *s* is the selection coefficient on the derived allele at time *t*. The value of *s* is *αβ*, where *α* is the selection gradient on the trait at time *t* and *β* is the effect size of the derived allele. (We set the value of *α* according to the expected trait variance at the onset of selection, which was 1.) This procedure is an approximation of allele-frequency dynamics under polygenic selection, but it is one that captures the overall patterns we seek to study (see supplementary text S1 and Supplementary Figure S1 for a comparison with truncation selection simulated forward in time).

After the allele-frequency trajectory was simulated, the polygenic-score trajectory calculated based on the allele-frequency trajectory was retained if the difference in population-mean PGS between the onset and end of the selection was within 5% of the expected change 2*NδtS*, where *δt* is the duration of selection in coalescent units and *S* is the selection differential on the PGS. We imposed this 5% cutoff in order to ensure that the degree of trait change during the period of selection was similar across iterations, and to ease comparisons with the results of Edge & Coop (2019), who used the same cutoff. It leads to selection of the 25-30% of simulated trait trajectories closest to the expectation in the parameter regime we used.

We used an unpublished modified version of `ms` (R. R. Hudson, 2002) called `mssel` (Berg & Coop, 2015) to produce simulated local coalescent trees and haplotypes across a region flanking the causal variant. In `mssel`, for most simulations, the sample size was set to 2,000 and the number of derived chromosomes was drawn as a binomial random variable with a size of 2,000 and success frequency equal to the contemporary allele frequency. We selected an effective population size *N* of 10,000, and a haplotype length of 200,000 base pairs (with the effect locus at position 100,000). The per-base-pair mutation rate was set as 2e-8, and the per-base-pair recombination rate was set as 2.5e-8. These values were transformed to population-scaled `mssel` inputs of `-r` 199.5 and `-t` 159.68. To explore additional scenarios, we also independently simulated larger sample sizes (5,000), haplotype of 500,000 base pairs, realistic human demography, genotyping error, and phasing error.

### 2.2 Software specifications

The coalescent trees produced by `mssel` for the selected sites are the true trees, and the haplotypes corresponding to the true trees were used as input to ARG estimation software to generate estimated trees. A brief description of each piece of estimation software can be found in the supplementary text. The ARG estimation programs vary with respect to the total sample size that can be used. We thus down-sampled the 2,000 simulated haplotypes generated in each simulation as necessary. We used 20 randomly drawn haplotypes as input to `ARGweaver`, and 20 and 200 haplotypes for `RENT+` and `SINGER`. For comparison with these methods, we also used the same subsamples of 20 and 200 haplotypes as input to `Relate` and `tsinfer+tsdate`. We did not use subsamples for `ARG-Needle` or `ASMC-clust`, which require sample sizes of at least 300 haplotypes.

#### ARGweaver

`ARGweaver` site input files were generated by a custom `Python` function. We ran the `arg-sample` program with the same values in the simulation (`-N` 10000, `-r` 2.5e-8, `-m 2e-8`) with the SMC’ model (`-smcprime`). The total number of sample iterations (`-n`) was set to 700 with the first 200 iterations as burn-in, and 50 estimated trees were extracted for one locus, one from every 10th iteration (`-sample-step 10`).

#### RENT+

Haplotypes generated by mssel were converted to `RENT+` format. We ran `RENT+` using -t to estimate branch lengths for local trees and `-l` to specify the proportional positions on the chromosomes.

#### Relate

We wrote a script to generate `Relate` haps, map, and sample files from simulated haplo-types. `Relate` was run with `-mode All, -m 2e-8` (mutation rate), and `-N 20000` (haploid effective population size). We used the add-on module `RelateExtract` to extract the tree corresponding to the selected site.

#### tsinfer+tsdate

The `tsinfer` input file was generated by calling the `add_site` function. Coalescent trees were estimated with `tsinfer` using default settings. The age of nodes in the tree was estimated using `tsdate` with the same parameter values as in the simulations (`Ne = 10000, mutation rate = 2e-8`). We converted the tree at the selected site to Newick format with the `as_newick` method in tskit.

#### ARG-Needle and ASMC-clust

The haps, map, and sample input file of `ARG-Needle` were generated by custom R functions. We set the `-normalize` parameter as the default value and used the constant 20K-sized (haploid population size) demography for `-normalize demography`. `ARG-Needle` requires a “decoding file” containing information about the demographic model, time discretization, and allele-frequency information. We created our own decoding file with the `prepare_decoding` function in `Python package asmc` – derived allele frequencies were calculated on the basis of the simulated haplotypes; the discretized time intervals were set as 14 intervals of 30 generations each starting in the present (0, 30, 60…, 420), followed by 16 more ancient intervals of 100 generations (520, 620, 720,…, 1920); the demography file was built with a constant population size (*N_e_* = 20, 000). The output tree was converted to Newick format with the `arg_to_newick` function from the arg needle lib package. Both choices for parameter `--mode` (‘array’ and ‘sequence’) were tested. To run `ASMC-clust`, we set the parameter `-asmc_clust` to 1. Other parameters were kept at the default values.

#### SINGER

The VCF file was generated from the simulated data by a custom R function. The mutation rate (`-m`) was set to 2e-8 and the ratio (`-ratio`) was set to 1.25. The population size (`-Ne`) was set to 10,000. `SINGER` was run for 700 iterations, with the first 200 iterations as burn-in. We took 50 samples from the rest, thinning every 10 iterations (`-step 10`). The estimated trees were stored as tree sequences. The marginal tree at the selected site was converted to Newick format with the `as_newick` method in `tskit`.

### 2.3 Branch length units

The ARG estimation tools we considered use different units for branch length. We standardized branch lengths to units of 2*N* generations for all trees. True trees from `mssel` are produced in units of 4*N* generations, so we multiplied the branch lengths by 2. In the manuscript describing `RENT+`, there is some ambiguity about the units, which are described as “standard coalescent units” (typically units of 2*N* generations), but in some calculations appear as if they are in units of 4*N* generations (Supplementary Table S1-S2). We applied the estimators on both the original branch lengths (assuming units of 2*N* generations) and a rescaled branch length (assuming the reported values are in units of 4*N* generations). We show the rescaled version in comparison with the other methods—although Edge & Coop (2019) treated the reported branch lengths as if they were in units of 2*N* generations, we believe that the makers of `RENT+` intended the branch lengths to be interpreted as in units of 4*N* generations. However, assuming 2*N* -generation units leads to slightly better performance on average (Supplementary Fig. S2-S3, Supplementary Table S1). The `ARGweaver`, `tsinfer+tsdate`, `ARG-Needle`, `ASMC-clust`, and `SINGER` trees are reported in units of generations, and we divided their original branch length by 2*N*. `Relate` trees use years as the unit and assume 28 years per generation when exported in Newick format, so we divided the original branch length by 28 ∗ 2*N*. Average times to most recent common ancestor (tMRCA) from the original branch length and the scaled branch length for each method are listed in Supplementary Tables S1 and S2.

### 2.4 Applying population-mean PGS trajectory estimators to the estimated trees

We applied the three estimators proposed in Edge & Coop (2019): the “proportion-of-lineages” estimator, the “waiting-time” estimator, and the “lineages-remaining” estimator (further description of the estimators in Supplementary text). All three estimators were applied to the estimated trees to obtain the estimated allele-frequency trajectories for each locus. For `ARGweaver` and `SINGER` trees, since there is more than one estimated tree for each locus, we applied the estimators to each sampled tree and took the average result from 50 trees as the estimated frequency trajectory for that locus. As discussed in Edge & Coop (2019), the proportion-of-lineages estimator is expected to perform well under neutral evolution but to be biased during periods of directional selection. The other two estimators are expected to be approximately unbiased given the true trees but to be much more variable. The waiting-time estimator, as written in Edge & Coop (2019) is not applicable in cases in which there are polytomies in a coalescent tree. We devised a scheme to address this limitation (see Supplementary text).

As a general note about the estimators of Edge & Coop (2019), we emphasize that the relationship between changes in the population-mean PGS and trait changes is not straight-forward. Even if PGS histories are estimated perfectly on an accurate PGS, phenotypes may be influenced by environmental changes and gene-by-gene or gene-by-environment interactions. Additionally, the linkage disequilibrium (LD) between markers and causal variants will change over time, causing the relationship between markers and the phenotype to change with it (Martin et al., 2017). Finally, some variants affecting the trait in the past will have been lost in the present (Carlson et al., 2022) or may no longer be detectable in GWAS. Nonetheless, Edge & Coop’s methods can be used to identify and roughly date patterns of allele-frequency change that are not consistent with neutral evolution.

### 2.5 Benchmarking metrics

We evaluated the performance of the ARG-estimation software as inputs to various approaches to PGS-history estimation, comparing them in terms of bias, mean squared error (MSE), 95% confidence interval coverage, and type I error rates and power (at the 0.05 level) of the *T_X_* statistic for testing directional selection. The *T_X_* statistic is computed as a sum of squared, standardized changes in an estimated population-mean PGS between pre-specified time points. That is, 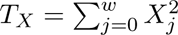, with

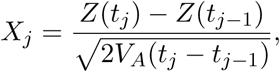

where *Z*(*t_j_*) is the population-mean PGS at time *t_j_*, and *V_A_* is the additive genetic variance of the PGS 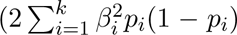). Under neutral evolution, *X_j_* approximately follows a standard normal distribution, and the allele-frequency changes are independent in distinct time intervals. Thus *T_X_*, the sum of the 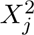, should approximately follow the *χ*^2^(*w*) distribution. However, under directional selection, the population-mean PGS changes more quickly than predicted under neutrality, leading to *T_X_* values that are large compared with the expected *χ*^2^(*w*) distribution.

The efficacy of Edge & Coop’s estimators depends on the accuracy of branch lengths in the estimated trees. To elucidate the differences in software performance, we extracted the pairwise coalescent times from true trees and estimated trees. Then we compared the point estimate of coalescent times between the two, and the overall distribution of pair-wise coalescent times from estimated trees against the expected exponential distribution, similar to the benchmarking recently performed by Brandt et al. (2022). To obtain the samples of pairwise coalescent times, we gathered trees containing the causal allele from 10 randomly selected simulated traits, resulting in a total of 1,000 trees. Specifically, for `Relate`, `tsinfer`, `ARG-Needle`, and `ASMC-clust`, we collected pairwise coalescence times from their 2,000-sample trees. For `RENT+` and `SINGER`, we used 200-sample subtrees, and for `ARGweaver`, we used 20-sample subtrees. In the case of `ARGweaver` and `SINGER`, we averaged pairwise coalescence time across 50 sampled trees per locus. The averaged times correspond to the same pairs of tips in each sampled tree. Due to memory constraints, we could not plot all pairwise coalescence times from one-thousand 200-sample and 2,000-sample trees. Instead, we randomly sampled 190,000 pairwise coalescence times, which is 1,000 times the total number of pairs of tips on a 20-tip tree (i.e. 1,000 times 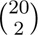).

We also examined the impact of a selection event on the distribution of pairwise coalescence times by setting the selection coefficient to ∼0.004, causing the derived-allele frequency to increase from approximately 0.3 to 0.7 during the selection period. Then 100 estimated trees with 300 samples were generated on the basis of this allele-frequency trajectory. We compared the MSE of the estimates from true trees with different sample sizes (20, 200, and 2,000 haplotypes) to assess the effect of the increased sample size.

To explore the relationship between measures of tree-topology accuracy and performance in estimation of population-mean PGS history, we examined the Robinson–Foulds (D. F. Robinson & Foulds, 1981) and Kendall–Colijn distance (Kendall & Colijn, 2016) between true trees and estimated trees and the proportion of estimated trees with mono-phyletic derived tips.

## 3 Results

### 3.1 Runtime comparison

The ARG-estimation tools we consider vary substantially in their runtime. One important difference is between tools that incorporate MCMC sampling (`ARGweaver` and `SINGER`) and those that do not. Both `ARGweaver` and `SINGER` sample trees from the posterior distribution, and the reported runtime represent 700 MCMC samples (200 burn-in). `SINGER` is, as expected, substantially faster than `ARGweaver`. Additionally, `tsinfer` has a clear advantage in speed with larger sample sizes. Average runtimes for every tool are listed in Table 1.

**Table 1:**
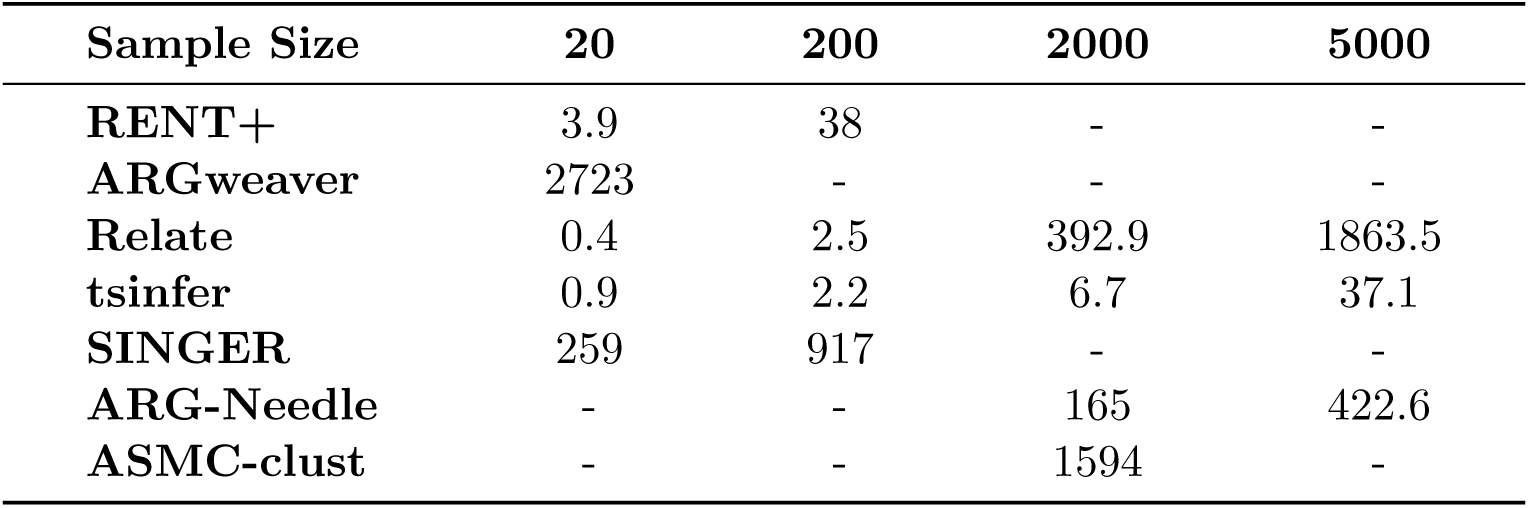
Average runtime for estimating ARGs for one 200kb segment of 20, 200, 2,000 and 5,000 samples from the ARG-estimation tools. Runtimes are measured in seconds.

### 3.2 Bias, mean squared error, and confidence interval coverage under neutrality

When traits are simulated under neutral evolution and the true trees are known, the proportion-of-lineages procedure can be viewed as a maximum-likelihood estimator for the allele-frequency history (Edge & Coop, 2019). The waiting-time and lineages-remaining estimators have previously been observed to be more variable than the proportion-of-lineages estimator with either the true trees or `RENT+` trees (Edge & Coop, 2019).

With respect to bias, we find that, in line with previous results, none of the estimators display much bias when estimated with the true trees or `RENT+` under neutrality. In addition, almost all newly tested tools/algorithms (`Relate`, `tsinfer+tsdate`, `ARGweaver`, `ASMC-clust`, and `SINGER`) follow similar patterns. Differences in bias resulting from trees estimated with these different pieces of software, especially during the recent past, and especially with the proportion-of-lineages estimator, are relatively minor (Fig. 2A-C, Supplementary Fig. S4-S6). Results for `ARG-Needle` are reported in the supplement and show more bias than other estimated trees (Supplementary Fig. S6). The likely reason for this is that `ARG-Needle`’s algorithm does not guarantee that mutations map to particular branches in the ARG. Instead, carriers of the derived allele may form polyphyletic groups on the local marginal tree. The frequency of non-monophyly among tips carrying the derived allele is much higher in `ARG-Needle` trees than in trees estimated by any other method (Supplementary Table S3). Although such polyphyly may be acceptable for many purposes, it can cause major problems for the estimators of Edge & Coop (2019), particularly in the ancient past. However, if `ARG-Needle` trees are manually modified to force monophyly among derived-allele-carrying tips, then the bias of `ARG-Needle` is similar to other methods (Supplementary Fig. S7). We also found that when the “–mode” parameter in `ARG-Needle` is set to “sequence”, the results improve over the default value of “array” (Supplementary Fig. S7), which is sensible given that the simulated data we provide to `ARG-Needle` includes all variants flanking the focal site.

**Figure 2:**
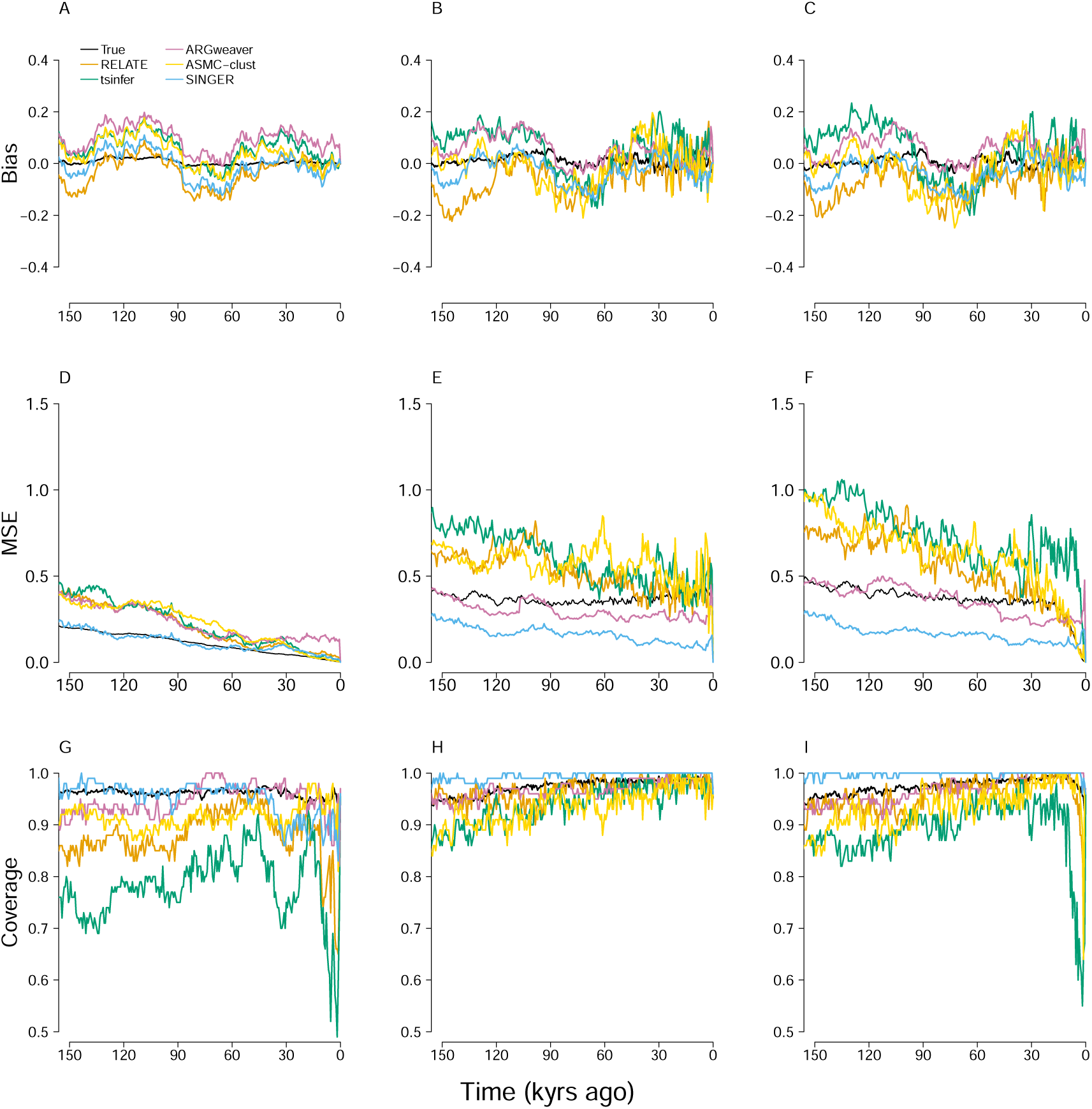
The bias (A-C), MSE (D-F), and confidence-interval coverage (G-I) of the proportion-of-lineages (left column), waiting-time (middle column), and lineages-remaining estimators (right column), with the true trees and estimated trees from each ARG-estimation method as input. For the estimates computed from true coalescent trees (black lines), 1000 simulations were performed with a sample of 2,000 chromosomes. In each simulation, the PGS was formed from 100 loci and evolved neutrally. `Relate`, `tsinfer`, and `ASMC-clust` were used to reconstruct trees with 2,000 chromosomes. `SINGER` was run with 200 chromosomes, and `ARGweaver` was run with 20 chromosomes. Lines represent means from 100 simulations. Times are displayed assuming diploids with *N_e_* = 10, 000 and a generation time of 30 years, i.e. one coalescent unit corresponds to 600,000 years.

The advantage of the proportion-of-lineages estimator over the waiting-time and lineages-remaining estimators under neutrality is more apparent when examining estimated MSE. In the three MSE plots (Fig. 2D-F), MSE tends to increase from the recent past into the distant past for all estimators and all software packages. This is unsurprising. Under neutrality, the variance of the proportion-of-lineages estimator is inversely related to the number of lineages ancestral to the sample (Edge & Coop, 2019). Thus, the fact that most lineages coalesce in the recent past implies that the estimator is more variable in the distant past. The proportion-of-lineages estimator (Fig. 2D) exhibits a much lower MSE than the other two estimators. The true trees, again unsurprisingly, provide the best MSE. However, among the estimated ARGs, the ones produced by software that scales to large samples do not necessarily show markedly better performance. Across much of the range examined, `SINGER`, which uses samples of only 200 haplotypes, has the lowest MSE, and in the more distant past, `ARGweaver`, which uses samples of only 20 haplotypes, is comparable with `ASMC-clust`, which uses samples of 2,000 haplotypes. In the MSE plots of the waiting-time and lineages-of-remaining estimators, the estimates derived from `ARGweaver` and `SINGER` trees consistently show relatively low MSE values (Fig. 2E, F).

We assessed credible interval coverage with `SINGER` and `ARGweaver` and confidence interval coverage with the true trees and all other methods (Fig 2G-I). The true trees produce acceptable coverage with all estimators. Credible intervals from `SINGER`-estimated trees also show consistently high coverage in most cases, though they drop below 95% coverage in the recent past with the proportion-of-lineages estimator. Other estimated trees, however, all produced somewhat lower coverage, with a tendency for coverage to decline into the more distant past.

`ARGweaver` and `SINGER` sometimes outperform the true trees in terms of MSE or credible interval coverage when the lineages-remaining and waiting-time estimators are used, a counterintuitive result. However, individual `ARGweaver` and `SINGER` trees do not outperform the true trees (Supplementary Fig. S8 and S9); it is only when results are averaged across the many trees produced by these methods that performance exceeds the true trees. The lineages-remaining and waiting-time estimators are noisy estimators that are sensitive to the timing of individual coalescent events, and this result suggests that their variability can be reduced by averaging results across many trees compatible with the data.

The bias, MSE, and coverage under neutrality with larger flanking regions, simulated genotyping error, and simulated phasing error can be found in Supplementary Fig. S10 - S13.

### 3.3 Bias, mean squared error, and confidence interval coverage under recent directional selection

Under selection, the proportion-of-lineages estimator is known to be biased, as the ancestors of the sample are not representative of the ancestral population from which they are drawn (Edge & Coop, 2019). The waiting-time and lineages-remaining estimators were developed to avoid this bias.

When there is a burst of directional selection in the recent past, the performance differences among ARG-estimation software packages are more pronounced. In contrast to the neutral scenario, and as expected, the proportion-of-lineages estimator is more strongly biased, has a higher MSE, and has lower confidence-interval coverage than the other estimators (Fig. 3A-I, Supplementary Fig. S14-S16), particularly during the period of selection and more anciently, as expected. Looking backward in time, the bias of the proportion-of-lineages estimator between the present and the end of the period in which selection occurred is low. This basic pattern also appears in the MSE and confidence-interval coverage plots, with good performance between the present and the end of directional selection, declining into the past during the period in which selection occurred, and then recovering during the neutral period that preceded selection.

**Figure 3:**
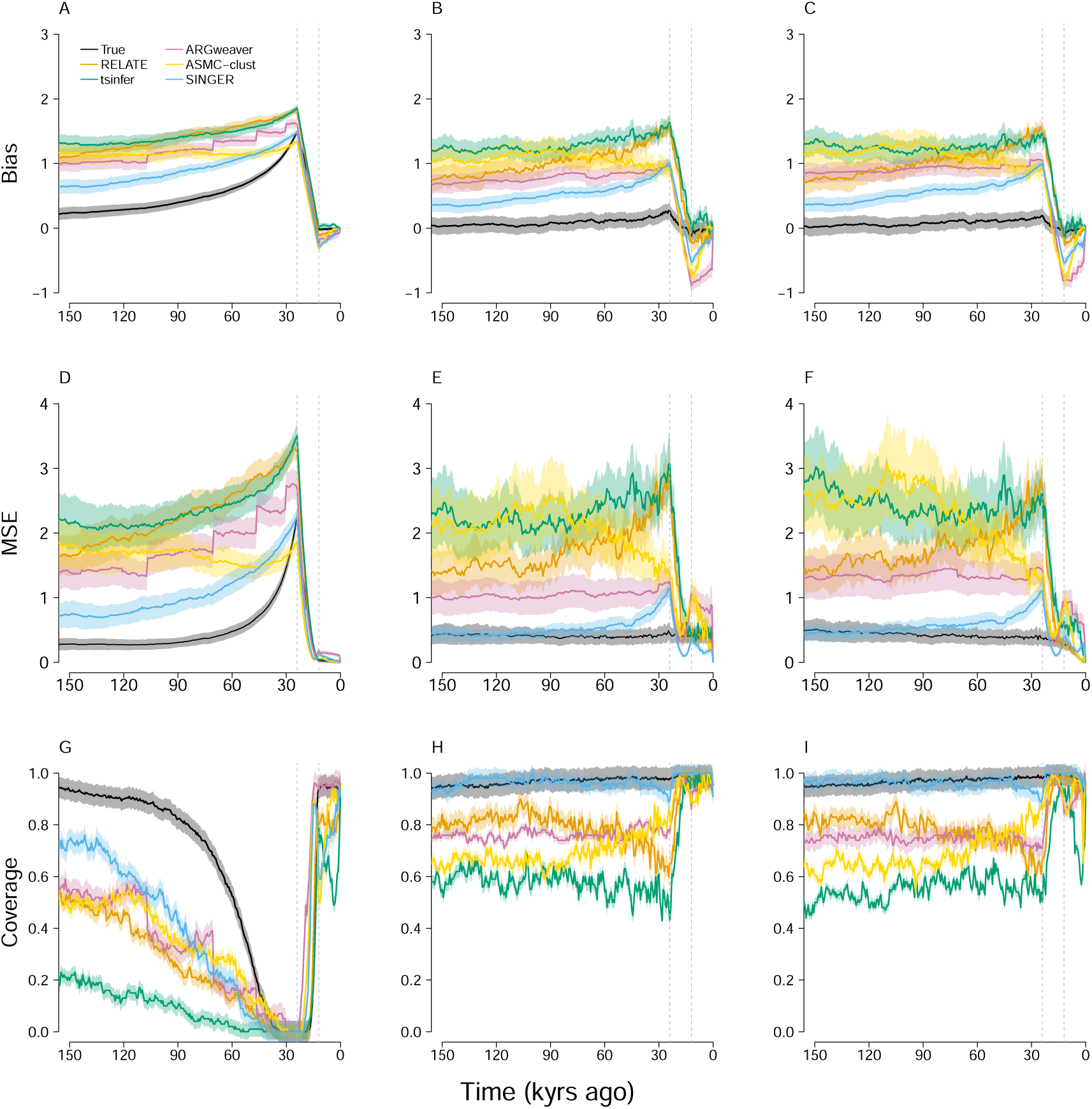
The bias (A-C), MSE (D-F), and confidence-interval coverage (G-I) of the proportion-of-lineages (left column), waiting-time (middle column), and lineages-remaining estimators (right column), with the true trees and estimated trees from each ARG-estimation method as input. The PGS is influenced by a period of directional selection that occurred from 0.04 to 0.02 coalescent units ago as represented by the vertical dotted lines. The parameters used to run all software are otherwise identical to those applied under neutrality (Fig. 2).

The waiting-time and the lineages-remaining estimators show substantially lower bias (Fig. 3B-C). The true trees generally show acceptable performance throughout the time period examined, with only slight bias during the period of selection and fairly uniform performance across time on other desiderata. However, all estimated trees produce noticeably worse performance on all criteria (Fig. 3B-I). Overall, among the estimated trees, the `SINGER` trees produce the lowest bias and MSE and interval coverage closest to the nominal level. `ARGweaver` ranks second across most of the investigated time. `ASMC-clust` and `Relate` are competitive in the recent and distant past respectively.

As in the neutral case, we also compared performance between individual trees and averages across many posterior `ARGweaver` and `SINGER` trees (Supplementary Figs. S17 and S18). We also compared the performance of original and modified `ARG-Needle` trees (Supplementary Figs. S19); and also for scenarios with larger flanking regions, simulated genotyping error, and simulated phasing error (Supplementary Figs. S20 - S23).

### 3.4 Power of *T_X_*

Next, we examined the type I error rate and power of tests of neutrality using *T_X_*, a test statistic sensitive to changes in a population-mean PGS that are larger than expected under neutrality. *T_X_*can be understood as a version of the *Q_X_* statistic (Berg & Coop, 2014) applied to a population-mean PGS from one population through time, rather than multiple populations sampled at the present (Edge & Coop, 2019). To use *T_X_*, one picks a set of time points to calculate the statistic 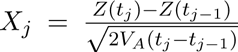,where *Z*(*t_j_*) is the estimated population-mean PGS at time *t_j_* and *V_A_* is the additive genetic variance of the PGS. With the true allele-frequency trajectories, under neutrality, the sum across time points 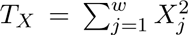 computed from *w* distinct intervals is approximately *χ*^2^(*w*) distributed (Edge & Coop, 2019).

In line with previous results, *T_X_* has an acceptable type I error rate when compared against the the *χ*^2^ distribution only when the proportion-of-lineages estimator is used to form the statistic (Table 2, Supplementary Table S4). With the proportion-of-lineages estimator, the observed type I error rates for the true trees, `tsinfer+tsdate` trees, `ASMC-clust` trees, and `SINGER` trees do not differ significantly from the nominal rate (`RENT+` and `ARG-Needle` trees in Supplementary Table S5 and S6). For `Relate`, the type I error rate is slightly higher than nominal. However, with all estimators and tree estimation software, calibrated type I error rates can be recovered by using a permutation distribution instead of the theoretical *χ*^2^ distribution, as expected.

**Table 2:** Type-I error/Power: Proportion-of-lineages. For the type I error simulations, asterisks indicate whether the observed type I error rate differs significantly from the nominal rate of 0.05: *p < 0.05; ** p < 0.01; ***p < 0.001.

| Input |  | Proportion-of-lineages |  |
| --- | --- | --- | --- |
| | | $\chi^2$ distribution | Permutation distribution |
| Neutral | True trees | 0.069 | 0.054 |
|  | **Relate** | 0.1* | 0.07 |
|  | **tsinfer** | 0.07 | 0.04 |
|  | **ASMC-clust** | 0.04 | 0.02 |
|  | **SINGER** | 0.09 | 0.05 |
| Selection | True trees | 0.982 | 0.981 |
|  | **Relate** | 0.14 | 0.21 |
|  | **tsinfer** | 0.19 | 0.07 |
|  | **ASMC-clust** | 0.69 | 0.57 |
|  | **SINGER** | 0.9 | 0.85 |

We also tested *T_X_*’s power in simulations that included a period of directional selection between 0.02 and 0.04 coalescent units in the past. In line with the previous results, *T_X_*calculated from the proportion-of-lineages-estimated allele frequencies is much more powerful than when the other two estimators of allele frequency are used. Considering proportion-of-lineages *T_X_*, power was approximately 98% with the true trees. Except for `SINGER`, most estimated trees lead to a substantial loss in power. Comparing against the permutation distribution, `SINGER` trees led to an observed power of 85%, whereas `ASMC-clust` produced *T_X_* statistics with a power of 57%, and `Relate` and `tsinfer+tsdate` trees produced power estimates of 21% and 7% respectively. Power obtained with `RENT+` trees with different branch length units can be found in Supplementary Table S5.

### 3.5 Comparison between true and simulated pairwise coalescence time

A major contributor to differences in performance among the tree-estimation procedures is the accuracy of the estimated coalescence times. We compared the pairwise coalescence times from estimated trees with their true values. As measured by the Spearman correlation between true and estimated pairwise coalescence times, `SINGER` outperforms other software packages, with `ARGweaver` and `ASMC-clust` close behind. It is also possible to see finer-grained patterns in the log-scaled coalescence times displayed in Figure 4. Overall, all three of `SINGER`, `ARGweaver`, and `ASMC-clust` show a tendency to overestimate coalescence times. But for short coalescence times, `ASMC-clust` tends toward underestimation (Fig. 4A,D). Both `Relate` and `tsinfer+tsdate` estimates appear biased for moderate-length coalescence times (between 0.1 and 1); `Relate` tends to underestimate these values, whereas `tsinfer+tsdate` tends to overestimate them (Fig. 4B-C). Additionally, `Relate` produces more variable estimates of short coalescence times than `tsinfer+tsdate`, as previously observed by Fan et al. (Fan et al., 2023). Finally, the time discretizations used by `ARGweaver` and `ASMC-clust` are visible as horizontal bands. These patterns are qualitatively similar when examined in simulations performed under neutrality (Supplementary Fig. S24, see also Brandt et al. 2022) and in general distributions of pairwise coalescence times (Supplementary Fig. S25-S26).

**Figure 4:**
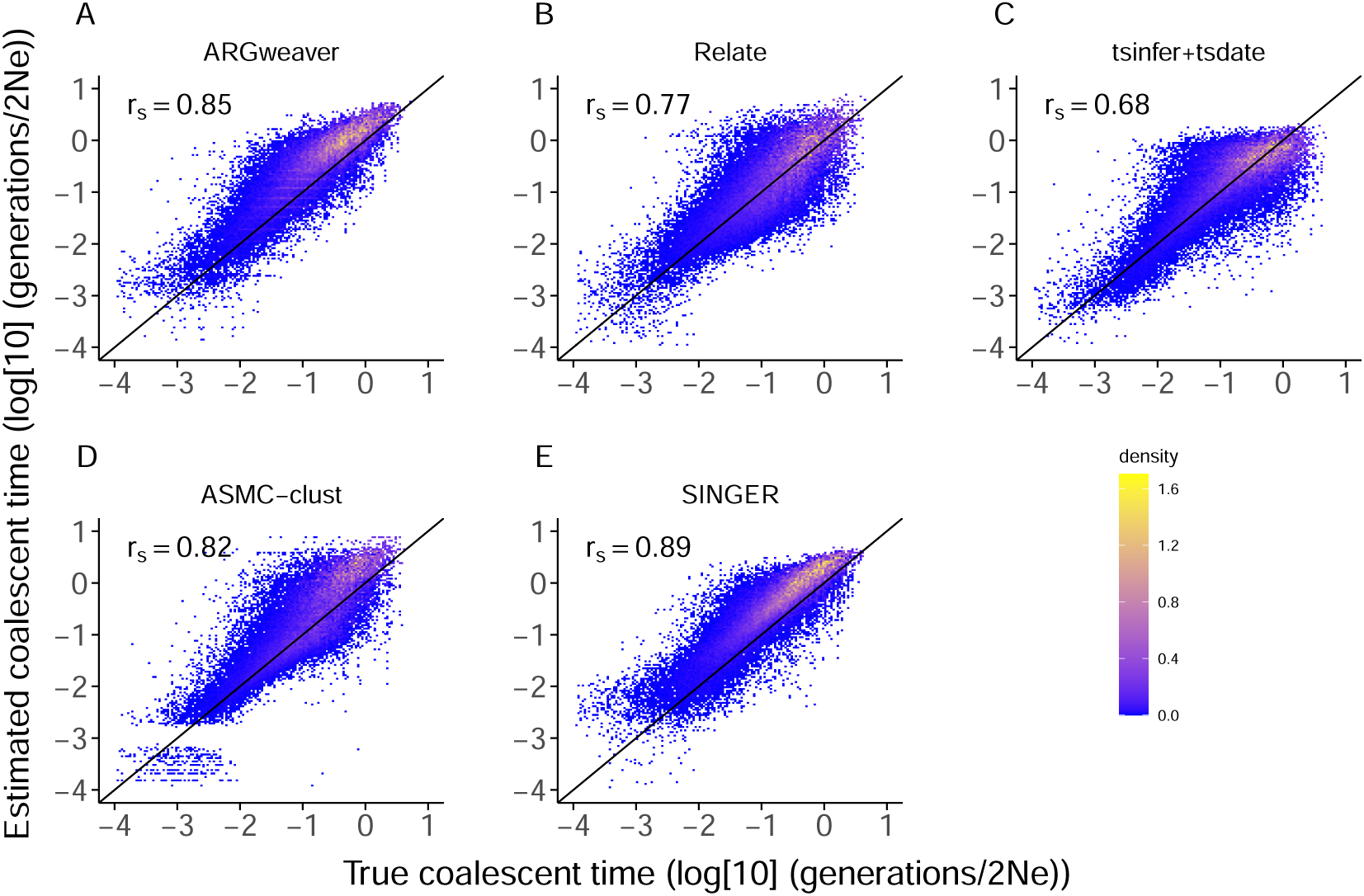
The comparison of pairwise coalescence times from true trees and estimated trees under directional selection. Each dot represents the true and estimated (log) coalescence time between a pair of samples. The diagonal line shows *x* = *y*, i.e. true times equal to estimated times. The values in the top-left corner show Spearman correlation coefficients.

To examine the effect of selection on coalescence time estimation, we checked the distribution of pairwise coalescence time from 100 trees with 300 tips, each subject to a selection event that increases the allele frequency from 0.3 to 0.7 between 0.04 and 0.02 coalescent units in the past. This selection event results in a high density of coalescences in the recent past within the derived-allele subtree, leading to a higher frequency of short coalescent times compared with the neutral simulations. This creates a distinct dip in the histogram of pairwise coalescence times (Fig. 5A). This pattern can also be observed, to varying degrees, in the distributions of pairwise coalescence times from estimated trees (Fig. 5B-E). Nevertheless, the dip in the histogram is wider and deeper when examining the true trees than when examining times from estimated trees. The density of recent coalescence times is also too low in estimated trees.

**Figure 5:**
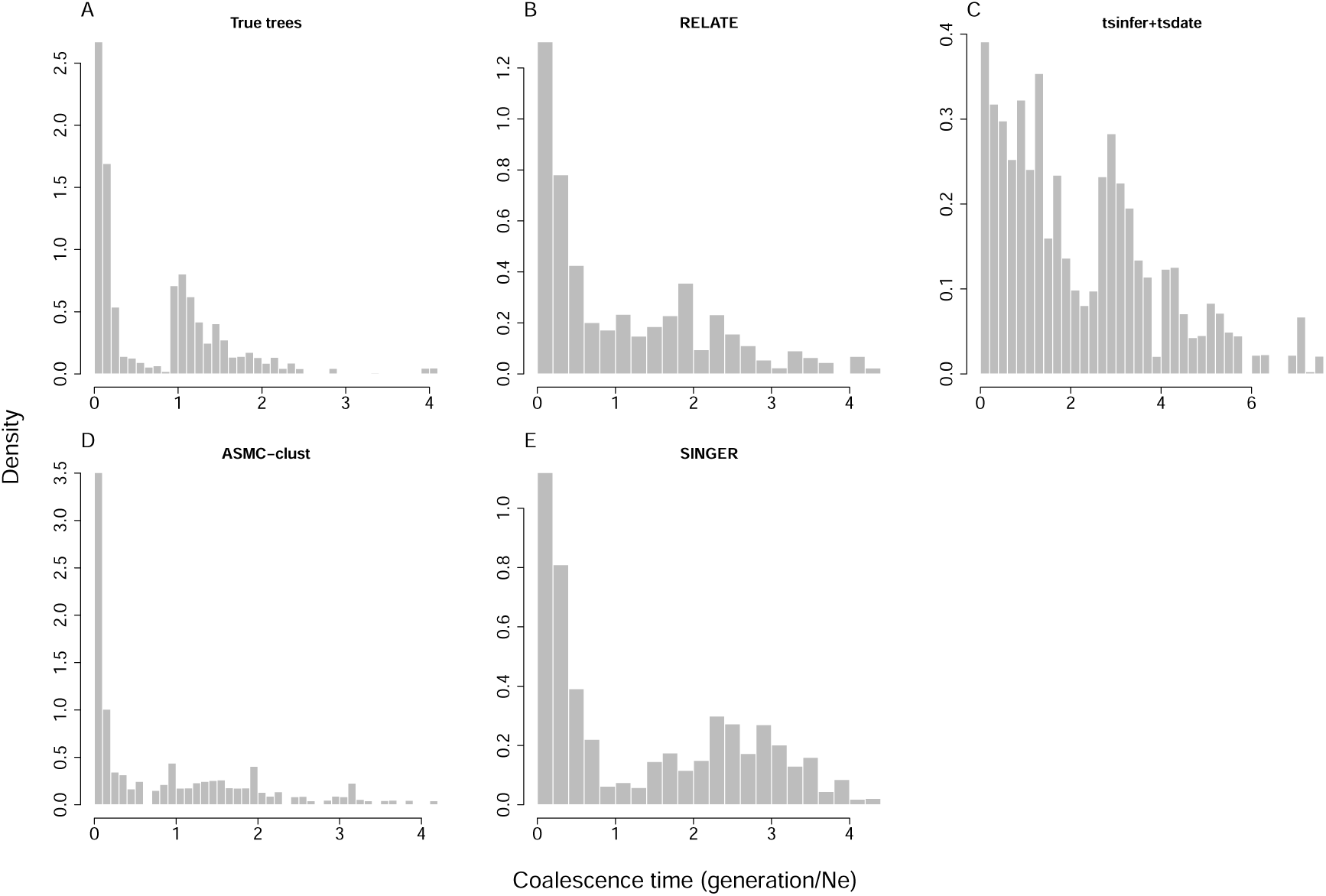
Distribution of pairwise coalescence times from true and estimated trees at a locus undergoing strong selection. Histograms show times from 100 trees with 300 tips each. Each tree (and flanking regions) was simulated assuming strong selection that increases the minor allele frequency from 0.3 to 0.7 between .04 and .02 coalescent units in the past.

Supplementary Table S7 shows the correlation between the true and estimated time to most recent common ancestor (tMRCA) for each software package and sample size under neutrality.

### 3.6 Topological aspects of ARG estimation accuracy

In addition to the correlation between true and estimated pairwise coalescence times, we considered other aspects of ARG estimation accuracy. Specifically, with respect to topology, we considered the rate of polyphyly among derived tips (and related measures), the Robinson–Foulds distance, and the Kendall–Colijn distance (with *λ* = 0, i.e. excluding branch length information). Results are shown in Supplementary Tables S3 and S8.

Each of these measures appears to track performance with respect to the estimators we consider here to some degree, but there are observations in which software that produces more accurate trees via each topological metric produces less accurate population-mean PGS histories. For example, with 200 tips, `RENT+` produces trees with lower Robinson–Foulds distances than `SINGER`, despite `SINGER`’s excellent performance on our benchmarks. With respect to the Kendall–Colijn distance, with 200 tips, `tsinfer` trees perform better than both `Relate` and `RENT+`, despite the latter’s generally better performance on our benchmarks.

### 3.7 Impact of sample size on estimation results

Another potential contributor to differences in performance among estimates derived from distinct ARG estimation procedures is sample size—ARG estimators differ in the sample sizes they can accommodate, and this variation is reflected in our simulations. Because large samples of haplotypes coalesce quickly in the recent past, we expect increased sample size to benefit allele-frequency trajectory estimation primarily in the recent past. We compared estimated PGS histories formed from the true trees obtained from samples of different sizes. The MSE of the proportion-of-lineages estimates decreases substantially when the sample size increases from 20 to 200. Notably, the MSE is not much larger for samples of size 200 than for samples of 2,000 lineages under neutrality, when this estimator is expected to perform well (Fig. 6A). Even in the recent past, the difference is slight. This pattern extends to the other two estimators when applied under selection (Fig. 6E-F). But under neutrality, the waiting-time and lineages-remaining estimators perform approximately equally regardless of sample size (Fig. 6B-C). The pattern observed for true trees also applies to estimated trees (Supplementary Fig. S4-S6, S14-S16).

**Figure 6:**
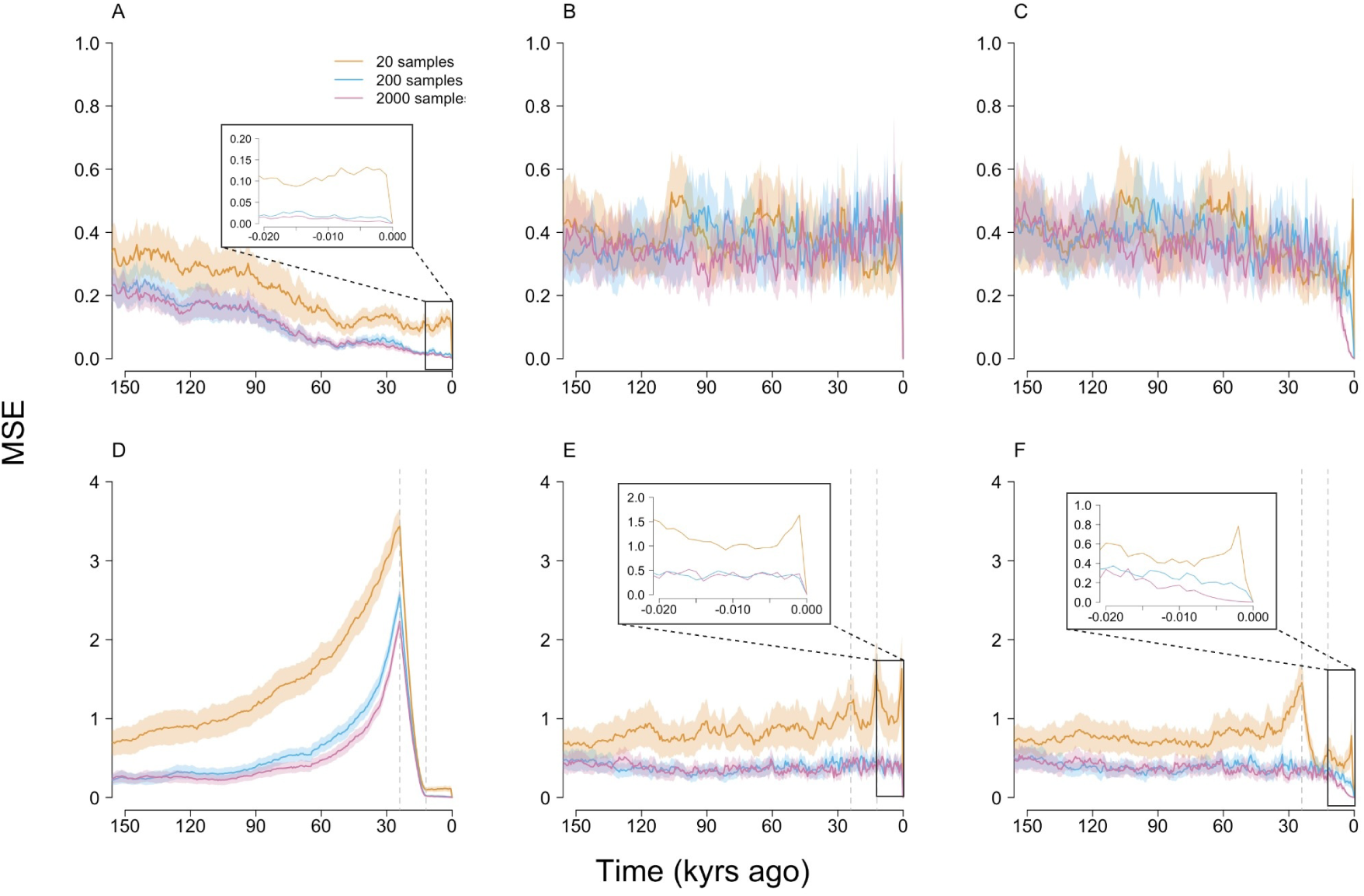
The MSE of the proportion-of-lineages (left column), waiting-time (middle column), and lineages-remaining estimators (right column) estimates from true trees with different sample sizes under neutrality (A-C) and directional selection (D-F).

Although we see little improvement in MSE when using true trees with samples of 2,000 relative to samples of 200 haplotypes, it is possible that large samples aid in marginal tree estimation, such that large samples produce better results with methods that can handle their size, not because the additional tips are helpful *per se*, but rather because the trees they produce are more nearly accurate. We do not find clear evidence of this possibility when increasing to 2,000 or 5,000 samples (Fig.S27, S28), but `tsinfer+tsdate` and `ARG-Needle` can accommodate samples much larger than this, which we do not explore.

### 3.8 Empirical data analysis

Edge & Coop (2019) analyzed data from the GBR (British) subsample of the 1000 Genomes project with respect to polygenic predictions of height formed from effect sizes from either GIANT (Wood et al., 2014) or the UK Biobank (Neale Lab, 2017). They found that GIANT effect sizes produced an impression of an increase in height over the last 60-90ky in ancestors of the GBR individuals, but UK Biobank effect sizes produced no such effect, consistent with recent (at the time) evidence that GIANT effect sizes were subject to biases from population stratification, small at the level of individual loci but large when combined into a polygenic score (Berg et al., 2019; Sohail et al., 2019). We repeated the analyses of Edge & Coop using trees estimated by `Relate`, `tsinfer+tsdate`, and `SINGER` (supplementary text and Fig. S29 - S31). The results are qualitatively similar to those of Edge & Coop (2019) in each case.

## 4 Discussion

We studied the performance of a set of methods for estimating the history of population-mean PGS using estimated ARGs, with a particular interest in the relative performance of different ARG estimation procedures. We used a broad range of methods appearing over the past decade—`ARGweaver` (Rasmussen et al., 2014), `RENT+` (Mirzaei & Wu, 2017), `Relate` (Speidel et al., 2019), `tsinfer+tsdate` (Kelleher et al., 2019; Wohns et al., 2022), `ARG-Needle`/`ASMC-clust` (Zhang et al., 2023), and `SINGER` (Deng et al., 2024)—that vary in their approaches, runtimes, and scalability. Previous work on these methods considered only estimated marginal trees from `RENT+` (Edge & Coop, 2019). We also benchmarked the estimated coalescence times emerging from these methods in general terms under neutrality and selection, providing an update on the work of Brandt and colleagues (Brandt et al., 2022), including several ARG estimation procedures that are newly released since their work.

Many of the basic patterns we observed were consistent across all ARG or marginal-tree estimation procedures, as well as being consistent with previous work. For example, regardless of the tree estimation procedure used, the resulting PGS history estimates are more accurate under neutral evolution than under selection, and PGS estimates designed to work either under neutrality (“proportion-of-lineages”) or under selection (“waiting-time” or “lineages-remaining”) had the expected pattern of relative performance.

At the same time, there were considerable differences in the accuracy of estimated population-mean PGS histories depending on the method used for marginal tree estimation. This is expected given the rapid development of ARG and marginal-tree estimation methods in the past ten years, which have resulted in increased scalability of several orders of magnitude (Brandt et al., 2022; Lewanski et al., 2024). However, bigger samples are not always better. In fact, the best performance overall from any set of estimated trees— whether measured in terms of mean squared error or confidence/credible interval coverage of the estimated PGS histories, or in terms of power in tests of natural selection—came from `SINGER` trees estimated with 200 haplotypes, rather than any of the tree estimation procedures that were capable of scaling to 2,000 haplotypes. There were several other cases in which smaller samples fit by either `RENT+` or `ARGweaver` outperformed trees estimated from much larger samples.

The fact that trees estimated with much larger samples of haplotypes do not always outperform trees estimated with smaller samples may be counterintuitive. One part of the explanation is the trade-off between accuracy and scalability in ARG estimation. ARG estimation is a difficult problem, and scalability can be achieved with simplifications that can reduce accuracy (Brandt et al., 2022, 2024; Deng et al., 2021, 2024). In line with this, `ARGweaver` and `SINGER` trees tend to feature more accurate branch lengths than methods that scale to larger samples (Brandt et al., 2022; Deng et al., 2024; Lewanski et al., 2024). The other important part of the explanation with respect to the estimators we explore here is that in large samples, coalescence initially happens very fast. Because the coalescence rate is proportional to 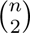, with *n* the number of lineages that have not yet coalesced, large samples imply large amounts of coalescence in the very recent past. As a result, even very large samples will be represented by a small number of lineages in the recent past (H. Chen & Chen, 2013; Frost & Volz, 2010; Griffiths, 1984; Jewett & Rosenberg, 2014; Maruvka et al., 2011; Slatkin & Rannala, 1997; Tavaŕe, 1984; Volz et al., 2009), and therefore increasing the sample size beyond a few hundred haplotypes produces more precise allele-frequency estimation primarily in the very recent past. This is reflected in Fig. 6, which shows that, even with the true trees, increasing the sample size from 200 to 2,000 does not clearly reduce the MSE of the Edge and Coop’s (2019) estimators in the scenarios in which they are each predicted to work well.

The ARG estimation methods we have examined have all been benchmarked previously. However, most of this benchmarking has been done with respect to general indicators of performance, such as the overall accuracy of local tree topologies (Kelleher et al., 2019; Rasmussen et al., 2014); a generalized Robinson–Foulds distance for ARGs (Zhang et al., 2023), the distribution of distances between topologically distinct local trees (Deng et al., 2021), and the distribution of pairwise coalescent times (Brandt et al., 2022; Deng et al., 2021). Although these general features are very important, it is not always straightforward to predict from them which methods will perform best when applied to a specific downstream task. For the estimators we explore here, at a given locus, all of them can be computed from the sets of coalescence times on the derived-allele and ancestral-allele subtrees. Thus, for these estimators, it is important that branch lengths are estimated accurately, as the branch lengths determine the coalescence times. Topology does not matter *per se*, but it does matter in practice, in that certain kinds of topological errors make it unlikely that the estimated coalescence times on each background will be nearly accurate. In line with this, we observe that `SINGER` excels in both our benchmarks and in preserving monophyly among tips carrying the derived allele, as expected in an infinite-sites mutational model.

For other tasks, other specific features may assume outsize importance. For example, in estimating the expected genetic relatedness matrix (eGRM), the estimation of long coalescence times matters a great deal, since many pairs of lineages, and thus many entries of the eGRM, are related by these long times, leading `Relate` to outperform `tsinfer+tsdate` for this task (Fan et al., 2022). In contrast, for some demographic inference problems, more recent coalescence times are more important, leading `tsinfer+tsdate` to outperform `Relate` (Fan et al., 2023). In addition to the specific features of the ARG that are important for a given downstream task, performance may vary according to the evolutionary scenario, e.g. the selection regime. In other words, there will not necessarily be an overall “best” method for ARG estimation—when choosing a method for ARG inference in empirical work, the specific strengths and weaknesses of particular methods may be more important than general considerations about overall accuracy or scalability.

This study adds to others suggesting that estimated ARGs and the marginal trees they encode are useful tools for the study of complex traits (M. Chen & Chiang, 2021; Edge & Coop, 2019; Link et al., 2023; Speidel et al., 2021; Stern et al., 2021b; Zhang et al., 2023). As ARG estimation methods continue to improve in both accuracy and scalability, they will open new opportunities for mapping genetic variants contributing to phenotypes, revealing polygenic adaptation, and exploring the relationship between natural selection and population history.

## 5 Code availability

Code and the compiled version of `mssel` used in this paper can be found at https://github.com/dandanpeng/ARG_Benchmarking.

## Acknowledgments

We thank members of the Edge, Mooney, and Pennell labs for helpful conversations and the anonymous peer reviewers for helpful comments on the manuscript. This work was supported by NIH grant R35GM137758 to MDE.

