## Supplementary text and figures for "Evaluating ARG-estimation methods in the context of estimating population-mean polygenic score histories"

Compiled on December 20, 2024

### S1 Simulation details: allele-frequency trajectories

The software we use for simulating true trees, `mssel`, is a modification of ms (Hudson 2002). `mssel` was written by Richard Hudson but never published by him, though it has been used in a number of other papers (Berg and Coop 2015; Lee and Coop 2017; Edge and Coop 2019; Lee and Coop 2019). `mssel` uses the structured-coalescent approach (Hudson and Kaplan 1988) to simulate coalescent histories for haplotypes linked to a selected variant, given a user-supplied allele-frequency trajectory for the selected variant. In words, `mssel` models coalescence on the background of each allele at the focal site as if haplotypes carrying those alleles formed variable-size populations back in time, with the changes in size corresponding to allele-frequency change. We provide a compiled version of `mssel` in our github repository.

We simulate polygenic selection in unlinked genomic regions containing one causal site each by using classic results about directional selection. Under directional selection on a single trait, the selection coefficient on an allele that has additive effect $\beta$ on the trait is approximately $\beta i/\sigma$, where $i$ is the selection intensity and $\sigma$ is the standard deviation of the trait (Falconer 1996; B. Charlesworth and D. Charlesworth 2010; Caballero 2020). We generate allele-frequency histories using this selection coefficient and then use them as input to `mssel`, which uses the approach of Hudson & Kaplan (Hudson and Kaplan 1988) to generate linked variation.

This scheme for simulating polygenic selection allows us to use backward-in-time simulations, saving time and computational resources, by considering the distribution of allele-frequency changes at each site marginally. It also allows us to condition on a set of causal variants common in the present. In real genomes, causal variants may be linked, leading to non-independence of allele-frequency change. Further, even in the absence of linkage, selection on a trait causes signed LD among causal loci, known as the Bulmer effect. This LD reduces additive variance for the first few generations, until an equilibrium is reached

(Falconer 1996; Caballero 2020). Ignoring these correlations can lead to differences in the total amount of trait change induced by selection, depending on genetic architecture (Walsh and Lynch 2018).

To explore these differences, we compared the simulation results obtained from a single-locus approximation with the previously described method and an explicit polygenic truncation selection simulation. Under the single-locus scheme, we simulate 0.02 coalescent units of selection on a quantitative trait using the scheme described in section 2.1 in main text, again using normally distributed effect sizes. We use a selection gradient corresponding to truncation selection eliminating the lowest 0.15% of the population, close to the value used in the main text. In the explicit polygenic truncation selection simulations, we compute individual-level trait values in each generation, eliminating the lowest 15/10,000 individuals, then generating the next generation of 10,000 diploids by picking parents to contribute haplotypes from the remaining individuals at random, allowing free recombination, and repeating this procedure for 400 generations, corresponding to 0.02 coalescent units in a diploid Wright–Fisher population of 10,000 individuals.

Simulation results from 100 iterations are shown in Fig. S1. Generally, the single-locus approximation produces a response fairly similar to truncation selection, both at the level of overall change in the trait and in terms of individual-site changes. The changes in the mean level of the trait induced by truncation selection are slightly smaller than the one-locus-at-a-time approximation. (1.84$\pm$ a standard error of .02 standard deviations on average, vs. $1.91 \pm .02$ standard deviations for the approximation. In contrast, 400 times the Breeder's equation expectation for the first generation gives a 1.96-standard-deviation change in the mean value.) Across all loci and simulation runs, both procedures led to a similar correlation of allele frequency and effect size after selection ($r = 0.18$ for the approximation and $r = 0.17$ for truncation selection.) In each simulation run, we included a site that started at minor allele frequency 0.1 and that initially contributed 0.01 to the additive variance (1/100 of the total), with an effect size of $\sqrt{1/18}$. On average, the frequency of this allele was somewhat higher at the end of selection in the approximation ($.16 \pm .006$) than in the truncation selection simulations ($.125 \pm .006$), but the overall distribution of the post-selection allele frequencies were qualitatively similar, with considerable variation among simulation runs.

### S2 Brief Descriptions of ARG Estimation Software

For completeness, we provide brief descriptions of the strategies for local tree/ARG estimation taken by the software packages we tested. More details are available in the referenced publications.

#### `ARGweaver`

`ARGWeaver` (Rasmussen et al. 2014) samples a data-compatible ARG generated by an approximation of the Sequentially Markov Coalescent (SMC, G. A. T. McVean and Cardin 2005) model that is discretized in time, called the Discretized Sequentially Markov Coalescent (DSMC). The key operation of `ARGweaver` is "threading," in which an ARG of $n$ sequences is sampled conditional on an ARG of $n-1$ sequences. The DSMC can be viewed as a hidden Markov model (HMM) with a state space defined by the possible local trees. Through the iterative process of removing and rethreading either sequences or subtrees, `ARGweaver` conducts inference by Markov-Chain Monte Carlo (MCMC).

#### `RENT+`

`RENT+` (Mirzaei and Wu 2017) is built on a previous heuristic genealogy inference approach called `RENT` (Wu 2011). The high-level idea underlying both `RENT` and `RENT+` is starting with a split (a bi-partition of rows) implied by the haplotypes at each site and then refining the local trees at specified sites. However, `RENT`'s approach is focused entirely on topology and ignores singletons because they are compatible with all possible tree topologies. `RENT+`, instead, starts from a set of "guide trees" formed by a version of UPGMA (Sokal and Michener 1958) at each site, modified to ensure that the resulting tree topology is compatible with an infinite-sites history for the immediate region. Because these guide trees are formed from local genetic distances, they incorporate singletons and thereby capture more information than the original version of `RENT`. Guide trees are used as the basis for forming local trees, in combination with a set of rules for "split propagation," ensuring that nearby pairs of local trees share topological information. The branch lengths and tree heights of the local trees are estimated from local haplotype Hamming distance matrices.

#### `Relate`

`Relate` (Speidel et al. 2019) constructs a distance matrix with rows estimating the relative order of coalescence events between a particular sequence and the remaining observed sequences using a modified Li & Stephens (N. Li and Stephens 2003) model, where the modification incorporates considerations of derived vs. ancestral allele status into the Li & Stephens approach. Then, the distance matrix is used to construct a rooted binary tree by a custom hierarchical clustering algorithm similar in spirit to UPGMA. When a mutation appears that cannot be mapped to the tree, a new tree is estimated. The branch lengths of the trees are estimated by mapping mutations onto each branch and applying an MCMC algorithm under a coalescent prior. A population-size history can be estimated alongside the branch lengths. Since an update (Stern et al. 2021), branch lengths can be sampled conditional on the estimated topology, but we do not consider samples from the branch length distribution here.

#### `tsinfer+tsdate0.2`

`tsinfer` (Kelleher, Wong, et al. 2019) proceeds by first constructing proxy ancestral haplotypes on the basis of shared derived alleles in a phased sample of contemporary haplotypes. The frequency of the derived allele is used as a proxy for the age of an ancestral haplotype, which also influences the length of the ancestral haplotype (since older haplotypes are likely to have been broken up by recombination). `tsinfer` then infers the most likely copying paths from the ancestral haplotypes to result in the contemporary sample under a modified Li & Stephens (N. Li and Stephens 2003) model. (Here, the modification to Li & Stephens is to add another state indicating non-ancestral material that cannot be copied to a contemporary sample). The copying paths determine a set of local trees, output as a `tskit` tree sequence (Kelleher, Etheridge, and G. McVean 2016). Then the ages of the ancestral haplotypes in the tree are inferred by an approximate Bayesian approach called `tsdate`, which relies on a generalization of the forward-backward algorithm the authors call the "inside-outside" algorithm (Wohns et al. 2022).

#### `ARG-Needle`

ARG-Needle (Zhang et al. 2023) leverages the output from the Ascertained Sequentially Markovian Coalescent (ASMC) (Palamara et al. 2018) algorithm. When given a new sample, ARG-Needle performs genotype hashing to detect a subset of candidate closest relatives rapidly and then uses the ASMC algorithm to estimate the pairwise coalescence time between the new sample and each of the candidate relatives. This output is then utilized by ARG-Needle to 'thread' the new sample to the ARG. After all samples have been threaded, ARG-Needle applies ARG normalization to refine the estimated node times, causing the estimated coalescence times to match more closely the times expected under the input demographic model.

#### `ASMC-clust`

Like `ARG-Needle`, `ASMC-clust` also utilizes the output from ASMC. `ASMC-clust` runs ASMC on all pairs of samples to obtain estimates of pairwise coalescence times. Then it performs the UPGMA clustering algorithm on the tMRCA matrix at each site to obtain a local tree. Its runtime scales quadratically with sample size.

#### `SINGER`

`SINGER` (Deng, Nielsen, and Song 2024) estimates coalescent trees by sampling from the posterior distribution, like `ARGweaver`, but it is heavily optimized in order to scale to larger data sets. A central innovation of `SINGER` is its two-step threading algorithm, which samples the joining branch in marginal trees for a new branch being threaded into the ARG and then samples the joining time for each of these joining branches. This operation makes the estimation algorithm faster by reducing the space of hidden states. A second key innovation is Sub-Graph Pruning and Re-grafting (SGPR), and extension of the subtree prune and regraph operation often used in phylogenetics. It prunes a sub-graph by introducing a cut to a random branch and extending the cut leftwards and rightwards to cover compatible trees, and re-grafts the subtrees by threading. SGPR is used for MCMC sampling.

### S3 Details of the allele-frequency time course estimators

Edge & Coop's (2019) estimators can be divided into two categories. The first estimator, the proportion-of-lineage estimator, treats the lineages in the marginal tree at time $t$ as representative of the ancient population at time $t$. It estimates $p_i(t)$, the allele frequency in the population at locus $i$ at time $t$, as the ratio of the counts of lineages at time $t$ that lead to contemporary haplotypes carrying the derived allele to the total number of lineages in the tree at time $t$,

$$\widehat{p_i(t)} = \frac{j_i(t)}{r_i(t)},$$

where $j_i(t)$ is the counts of lineages that carry the derived allele at locus $i$ at time $t$ and $r_i(t)$ is the total number of lineages ancestral to all contemporary samples at time $t$.

The other two estimators are called the "waiting-time estimator" and the "lineages-remaining estimator." Both of them consider the chromosomes in the sample population as if they were drawn from two separate "populations" defined by the allele they carry in the present. They attempt to estimate the historical sizes of these two "populations" using standard demographic inference techniques and to compare the number of estimated ancient carriers of the derived and ancestral alleles. The waiting-time approach estimates the "population" sizes according to the time interval between fixed numbers of coalescent events (Pybus, Rambaut, and Harvey 2000). The lineages-remaining approach estimates the same quantity by counting the number of coalescent events occurring between pre-specified time points.

The waiting-time estimator is written as

$$\widehat{p_i(t)} = \frac{\widehat{N_i(t)}}{\widehat{N_i(t)} + \widehat{M_i(t)}},$$

where $\widehat{N_i(t)} = \frac{Y}{2[1/(n_i-l)-1/n_i]}$. $N_i(t)$ and $M_i(t)$ are the size of the two separate subpopulations (i.e. the chromosomes ancestral to the two alleles) at locus $i$ at time $t$. Their values are estimated by a generalized skyline approach (Ho and Shapiro 2011; Pybus, Rambaut, and Harvey 2000). $Y$, the numerator of the $N_i(t)$ estimator, is the time elapsed while waiting for $l$ coalescent events, and $n_i$ is the number of lineages ancestral to lineages carrying the derived allele at the more recent end of the interval (i.e. more recently than the l coalescent events). Replacing these terms with the values from the ancestral allele's background gives us the estimated value of $M_i(t)$. Both $N_i(t)$ and $M_i(t)$ are assumed constant between coalescent events. However, this estimator is not applicable in cases when there are polytomies in a coalescent tree. We devised an approach to address this limitation (see below).

In the lineage-remaining approach, the estimator for $N_i(t)$ is

$$\widehat{N_i(t)} = \frac{\Delta t}{2\{\log[\frac{n_i^{(t)}}{n_i^{(t)}-1}] - \log[\frac{n_i^{(0)}}{n_i^{(0)}-1}]\}}.$$

$\Delta t$ is a prespecified time interval, and $n_i^{(0)}$ and $n_i^{(t)}$ correspond to the number of lineages that remain at the more ancient end and more recent end of the interval, respectively. Similar to the waiting-time estimator, during the $\Delta t$ interval, $N_i(t)$ is assumed to be constant. To estimate $M_i(t)$, the number of carriers of the ancestral allele, the same function is used, replacing $n_i^{(t)}$ and $n_i^{(0)}$ analogous quantities for lineages subtending the ancestral alleles, $m_i^{(t)}$ and $m_i^{(0)}$.

### S4 Update of the waiting-time estimator

The original waiting-time estimator does not allow for polytomies in a coalescent tree. If a polytomy exists, the resulting waiting time of zero will disrupt the allele frequency estimation. For those coalescent events involved in a polytomy, we assign each of them a new waiting time. Specifically, let $n_1$ be the number of lineages not yet coalesced before the polytomy, $n_2$ be the number of lineages remaining after the polytomy (so the polytomy involves $n_1 - n_2 + 1 = k + 1$ lineages coalescing to 1 lineage). Further, let $t_1$ be the time of the most ancient coalescence that is more recent than the polytomy, and let $t_2$ be the time of the polytomy (so the waiting time for the polytomy is $t_2 - t_1 = w$). We then assign $k-1$ new coalescence times occurring between $t_1$ and $t_2$, retaining the coalescence time $t_2$. Denoting these times $\tau_1, ..., \tau_{k-1}$, they have the values

$$\tau_i = t_1 + w\frac{\sum_{j=k+1}^{k+2-i}\frac{1}{j(j-1)}}{\sum_{j=k+1}^{2}\frac{1}{j(j-1)}},$$

The idea behind these approach is to assign coalescence times within the polytomy of $k+1$ lineages proportional to the expected coalescence times in a neutral coalescent tree of $k+1$ lineages.

### S5 Fixing the lineages-remaining estimator for edge cases

If there is no coalescence among ancestral lineages or derived lineages, which occurs if there is only one ancestral or derived lineage, then we replace the estimated allele frequency with the proportion-of-lineages estimate.

### S6 Modifying `ARG-Needle` trees

`ARG-Needle` trees do not ensure that mutations map to unique branches. As a result, the subtree formed by tips carrying the derived allele in the present is often polyphyletic with respect to the full tree. This introduces systematic errors into the allele-frequency history estimators of Edge and Coop (2019). To produce modified `ARG-Needle` trees, we traversed all internal nodes, evaluating each as to whether (1) all the tips descending from it correspond to haplotypes carrying the derived allele and (2) the node immediately above has tips descending from it carrying the ancestral allele. Any internal node meeting these criteria was recorded as the MRCA of a derived clade. Upon collecting the MRCA of all derived clades, we designated the most ancient MRCA as the new root for the entire derived subtree. We implemented this by updating the vector of coalescence times in `R`, adding the time for the new root and removing all older times. Accordingly, the number of lineages of each type of allele at different time points was also updated.

### S7 Calculating bias, MSE, and CI coverage

In all figures in which they are shown, the bias, MSE and the 95% interval coverage are estimated on the basis of 100 simulations. For each trait, the bias was estimated as the average difference between the estimated and true population-mean PGS across the 100 simulations evaluated every .001 coalescent units. The MSE was estimated as the average of the squared differences between the estimated and true population-mean PGS. The 95% interval coverage was estimated as the proportion of simulations in which the true population-mean PGS was within the confidence (or credible) interval computed from the estimated population-mean PGS. Where they are shown, confidence bands represent $\pm 1.96$ standard errors (estimated from the 100 simulations) above and below each of these measurements. For tools that generate multiple estimated trees at each focal locus (ARGweaver and SINGER), we first estimated the allele-frequency trajectory from each individual tree. These trajectories were then averaged across all trees, and the resulting average trajectory was used to calculate the estimated population-mean PGS.

### S8 Simulations of realistic human demography and data errors

#### Simulations with CEU demography

To explore the effect of a more realistic human demography, we provided a demographic model based on CEU to `mssel` in generating haplotypes and true trees. The demographic data was inferred by SMC++ (Terhorst, Kamm, and Song 2017) and rescaled to assume a mutation rate of 2e-8. We obtained the demographic information from the github repository (https://github.com/popgenmethods/pyrho/tree/master) for pyrho (Spence and Song 2019). In generating allele-frequency trajectories, we still assume that at the most recent time point prior to the onset of selection (or at the present in simulations without selection), causal sites have minor allele frequencies that follow the expectation for a standard neutral SFS, subject to the constraint of minor allele frequency greater than 1%. Compared with the CEU demography, this procedure generates more minor alleles with higher frequency than would be expected. We note that the process of GWAS ascertainment, which we do not model explicitly, would also be expected to lead to a bias for larger minor allele frequencies.

#### Simulation of genotyping error

To simulate genotyping error, we introduced errors in the haplotypes output by mssel, switching either 0 alleles to 1 or vice versa with an error rate of 0.1%. We chose this error rate on the basis of a study combining  2000 datasets from seven different sequencing instruments (Stoler and Nekrutenko 2021).

#### Simulation of phasing error

To simulate phasing error, we ran state-of-the-art phasing software on the genotypes produced by mssel. Specifically, we ran Beagle5.4 on mssel output haplotype with default phasing parameters (Browning et al. 2021) and used the output haplotypes as input to ARG-estimation software that requires phased data.

### S9 Analysis of empirical data on human height

Edge & Coop (2019) conducted an empirical analysis of estimated population-mean polygenic score histories for human height in the ancestors of the GBR subset of the 1000 Genomes Project panel. We repeated their analysis using ARG-estimation tools that were not available at the time of their work. We ran Relate, tsinfer+tsdate, and SINGER with height-associated SNPs used by Edge & Coop to get the estimated population-level polygenic score trajectory. Edge & Coop obtained SNP information including SNP ID, chromosome, and position from Racimo et al. (2018). They selected the SNP with the lowest p-value for an association test with height in each of 1700 nearly-independent linkage disequilibrium blocks identified by Berisa & Pickrell (2016). Effect sizes and p-values for the selected SNPs were taken from Neale Lab (2017) for UK Biobank and Wood et al. (Wood et al. 2014) for the 2014 GIANT consortium effect sizes. The sequence information is from 91 genomes in GBR subsample of 1000 Genome Project Consortium (2015). Taking each selected SNP as the focal site, we extended 100,000 bases on each side with tabix(H.

Li 2011). The resulting .vcf file was transferred to other formats than can be read by the various ARG estimation tools with `vcftools` or customized `R` functions.

As in Edge & Coop's analysis using `RENT+`, the $T_X$ statistic does not show strong evidence of selection on height over the past 60,000 years using either GIANT or UK Biobank effect size estimates. Specifically, we ran $T_X$ using a grid of times $0, 0.01, 0.02, ..., 0.1$ in coalescent units. With `Relate`, GIANT effect sizes produced $T_X = 22.3$, $p = 0.27$, and UK Biobank effect sizes produced $T_X = 12.0$, $p = 0.91$. With `tsdate+tsinfer`, GIANT effect sizes produced $T_X = 10.0$, $p = 0.12$, and UK Biobank effect sizes produced $T_X = 11.56$, $p = 0.06$. With `SINGER`, GIANT effect sizes produced $T_X = 25.8$, $p = 0.25$, and UK Biobank effect sizes produced $T_X = 7.6$, $p = 0.58$.

### S10 Supplementary Tables

Table S1 Average tMRCA of 100 trees in original units.

| **Sample size** | `mssel` (true trees) | `ARGweaver` | `RENT+` | `Relate` |
|---|---|---|---|---|
| **20** | 1.128 | 43970.640 | 1.096 | 993513 |
| **200** | 1.136 | - | 1.146 | 991855.1 |
| **2,000** | 1.215 | - | - | 982576.9 |

| **Sample size** | `tsinfer+tsdate` | `ARG-Needle` | `ASMC-clust` | `SINGER` |
|---|---|---|---|---|
| **20** | 36349.8 | - | - | 46178.76 |
| **200** | 35321.18 | - | - | 47080.96 |
| **2,000** | 34187.2 | 51243.16 | 44926.38 | - |

Table S2 Average tMRCA of 100 trees after branch-length rescaling.

| Sample size | mssel (true trees) | ARGweaver | RENT+ | Relate |
|---|---|---|---|---|
| **20** | 2.257 | 2.198 | 2.192 | 1.774 |
| **200** | 2.273 | - | 2.293 | 1.771 |
| **2,000** | 2.430 | - | - | 1.754 |

| Sample size | tsinfer+tsdate | ARG-Needle | ASMC-clust | SINGER |
|---|---|---|---|---|
| **20** | 1.817 | - | - | 2.308 |
| **200** | 1.766 | - | - | 2.354 |
| **2,000** | 1.709 | 2.562 | 2.246 | - |

Table S3 Proportion of trees with monophyletic tips

| Software | Sample size | monophyletic derived tips | monophyletic ancestral tips | monophyletic anc tips & non-monophyletic derived tips |
|---|---|---|---|---|
| **mssel** | 20 | 1 | 0.37 | 0 |
| | 200 | 1 | 0.33 | 0 |
| | 2000 | 1 | 0.320 | 0 |
| **ARGweaver** | 20 | 0.96 | 0.39 | 0.03 |
| **RENT+** | 20 | 0.93 | 0.34 | 0.08 |
| | 200 | 0.92 | 0.33 | 0.07 |
| **RELATE** | 20 | 0.86 | 0.24 | 0.004 |
| | 200 | 0.80 | 0.18 | 0 |
| | 2000 | 0.70 | 0.15 | 0 |
| **tsinfer+ tsdate** | 20 | 0.81 | 0.23 | 0.03 |
| | 200 | 0.73 | 0.17 | 0.015 |
| | 2000 | 0.56 | 0.14 | 0.019 |
| **SINGER** | 20 | 0.99 | 0.42 | 0.01 |
| | 200 | 0.99 | 0.34 | 0.0004 |
| **ARG-Needle** (array) | 2000 | 0.40 | 0.35 | 0.17 |
| **ARG-Needle** (sequence) | 2000 | 0.10 | 0.26 | 0.23 |
| **ASMC-clust** | 2000 | 0.76 | 0.34 | 0.02 |

Table S4 Type-I error/Power: Waiting-time and Lineages-remaining

| Input | | Waiting-time | | Lineages-remaining | |
|---|---|---|---|---|---|
| | | $\chi^2$ distribution | Permutation distribution | $\chi^2$ distribution | Permutation distribution |
| Neutral | True trees | 0.078*** | 0.061 | 0.045 | 0.045 |
| | Relate | 0.23*** | 0.05 | 0.04 | 0.05 |
| | tsinfer | 0.12** | 0.08 | 0.01 | 0.07 |
| | ASMC-clust | 0.02 | 0.03 | 0.03 | 0.08 |
| | SINGER | 0.35*** | 0.04 | 0.13 | 0.09 |
| Selection | True trees | 0.09 | 0.077 | 0.062 | 0.084 |
| | Relate | 0.28 | 0.05 | 0.08 | 0.1 |
| | tsinfer | 0.29 | 0.06 | 0 | 0.01 |
| | ASMC-clust | 0.01 | 0.05 | 0.04 | 0.06 |
| | SINGER | 0.56 | 0.14 | 0.3 | 0.15 |

For the type I error simulations, asterisks indicate whether the observed type I error rate differs significantly from the nominal rate of 0.05: $*p < 0.05; **p < 0.01; ***p < 0.001$. Relate, tsinfer, and ASMC-clust are run with 2,000 samples; SINGER is run with 200 samples.

Table S5 Power/type-I error of the RENT+ tree (200 samples) implementation of the $T_X$ statistic.

| | | **Proportion-of-lineages** | | **Waiting-time** | | **Lineages-remaining** | |
|---|---|---|---|---|---|---|---|
| | | $\chi^2$ distrib. | Permutation distrib. | $\chi^2$ distrib. | Permutation distrib. | $\chi^2$ distrib. | Permutation distrib. |
| **Neutral** | RENT+ ($2N$) | 0.11 | 0.05 | 0.33 | 0.04 | 0.05 | 0.05 |
| | RENT+ ($4N$) | 0.07 | 0.03 | 0.01 | 0.02 | 0.05 | 0.03 |
| **Selection** | RENT+ ($2N$) | 0.59 | 0.42 | 0.05 | 0.07 | 0.02 | 0.04 |
| | RENT+ ($4N$) | 0.28 | 0.23 | 0.05 | 0.07 | 0.05 | 0.06 |

The number in parentheses is the assumed units in which RENT+ branch lengths are.

Table S6 Power/type-I error of the `ARG-Needle` trees (2,000 samples) implementation of the $T_X$ statistic.

| | | Proportion-of-lineages | | Waiting-time | | Lineages-remaining | |
|---|---|---|---|---|---|---|---|
| | | $\chi^2$ distrib. | permutation distrib. | $\chi^2$ distrib. | permutation distrib. | $\chi^2$ distrib. | permutation distrib. |
| **Neutral** | array | 0.23 | 0.03 | 0.01 | 0.04 | 0.05 | 0.08 |
| | modified | 0.14 | 0.04 | 0.02 | 0.06 | 0.04 | 0.06 |
| | sequence | 0.29 | 0.05 | 0 | 0.07 | 0.01 | 0.04 |
| **Selection** | array | 0.67 | 0.12 | 0 | 0.03 | 0 | 0.04 |
| | modified | 0.56 | 0.21 | 0 | 0.04 | 0.04 | 0.05 |
| | sequence | 0.63 | 0.39 | 0 | 0.03 | 0.09 | 0.1 |

"array" and "sequence" refers to ARG-Needle trees estimated by setting "–mode" as those values, "modified" means the manually modified ARG-Needle trees from "array" mode

Table S7 Pearson correlation between true and estimated TMRCA for different software and sample sizes.

| Software | Sample size | Correlation coefficient |
|---|---|---|
| ARGweaver | 20 | 0.63 |
| RENT+ | 20 | 0.56 |
| | 200 | 0.47 |
| RELATE | 20 | 0.51 |
| | 200 | 0.49 |
| | 2000 | 0.50 |
| tsinfer+tsdate | 20 | 0.39 |
| | 200 | 0.32 |
| | 2000 | 0.38 |
| SINGER | 20 | 0.64 |
| | 200 | 0.62 |
| ARG-Needle (array) | 2000 | 0.14 |
| ARG-Needle (sequence) | 2000 | 0.49 |
| ASMC-clust | 2000 | 0.45 |

Table S8 Mean and standard deviation of topological distances between 1,000 estimated trees and true trees.

| **Software** | **Sample size** | **Robinson–Foulds** | **Kendall–Colijn** |
|---|---|---|---|
| `ARGweaver` | 20 | 0.43 ± 0.14 | 17.76 ± 5.98 |
| `RENT+` | 20 | 0.37 ± 0.14 | 16.57 ± 6.36 |
| | 200 | 0.46 ± 0.04 | 249.05 ± 86.50 |
| `RELATE` | 20 | 0.47 | 16.82 ± 5.29 |
| | 200 | 0.47 ± 0.04 | 261.47 ± 105.93 |
| | 2000 | 0.71 ± 0.01 | 4431.88 ± 1985.97 |
| `tsinfer+tsdate` | 20 | 0.47 ± 0.15 | 17.61 ± 6.50 |
| | 200 | 0.55 ± 0.046 | 240.47 ± 74.73 |
| | 2000 | 0.77 ± 0.01 | 3194.99 ± 1048.07 |
| `SINGER` | 20 | 0.40 ± 0.04 | 15.22 ± 5.11 |
| | 200 | 0.50 ± 0.039 | 206.35 ± 59.64 |
| `ARG-Needle (array)` | 2000 | 0.91 ± 0.025 | 5452.52 ± 1302.77 |
| `ARG-Needle (sequence)` | 2000 | 0.73 ± 0.012 | 7784.04 ± 3990.52 |
| `ASMC-clust` | 2000 | 0.72 ± 0.014 | 2814.64 ± 538.08 |

The Robinson–Foulds distance is normalized by the number of splits.

### S11 Supplementary Figures

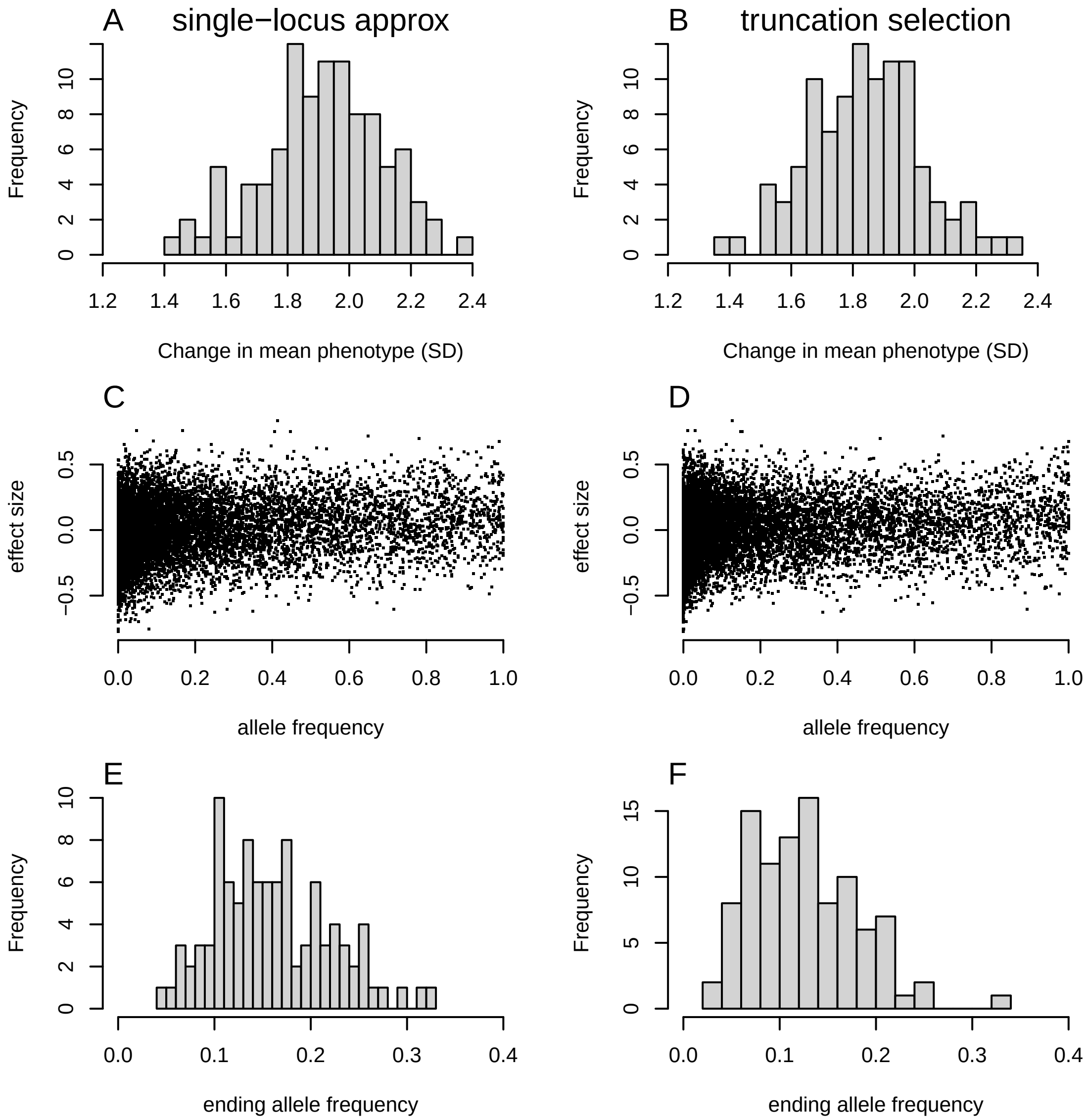


Figure S1: Comparison between the simulation results of single-locus approximation scheme and polygenic truncation scheme described in section S1. The left column shows results for the backward in time, one-locus-at-a-time approximation used in the main text; the right column shows results from forward-time simulations of truncation selection. A-B show the distribution of the change in the population-mean phenotype across 100 simulations. C-D show the relationship of effect size on the trait and allele frequency after selection. E-F show the distribution of ending frequencies for a locus starting at minor allele frequency 0.1 and with effect size $\sqrt{1/18}$ on the phenotype at the end of selection.

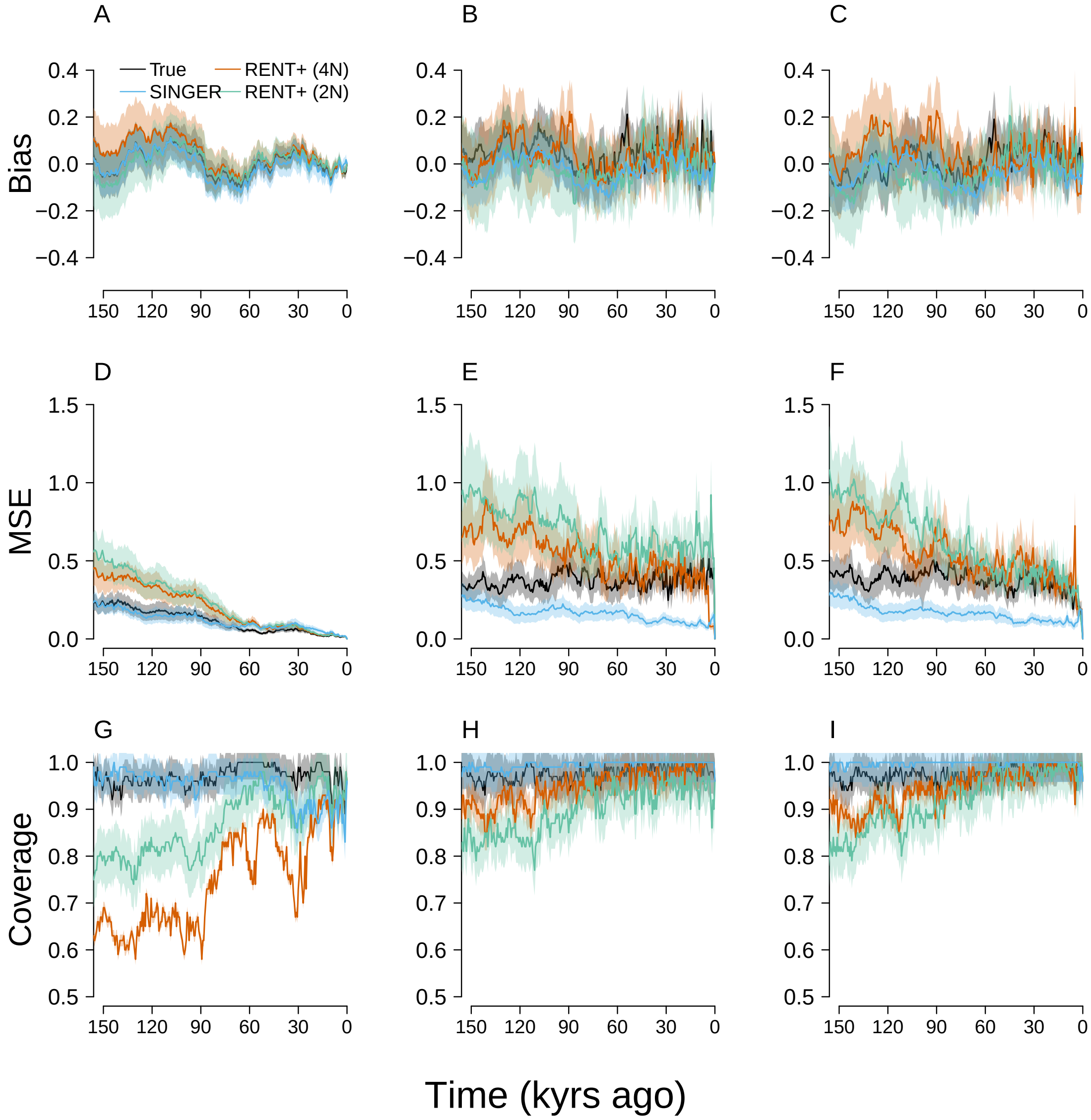


Figure S2: Investigation of branch-length scaling in `RENT+`. Here, we compare bias (A-C), MSE (D-F), and confidence-interval coverage (G-I) of the proportion-of-lineages (left column), waiting-time (middle column), and lineages-remaining estimators (right column), with estimates from `RENT+` trees assuming either $2N$ or $4N$ generations as units of branch length. Results for true trees and `SINGER` trees are shown for reference. In each simulation, the PGS was formed from 100 loci and evolved neutrally. 100 simulations were performed with 200 chromosomes. The reason we tested alternative scalings is that there is some ambiguity about the units of branch length in `RENT+` manuscript. The tMRCAs of `RENT+` trees (Supplementary Tables S1-S2) suggest that `RENT+` trees are reported in units of $4N$ generations. This contrasts with the assumption of Edge and Coop (2019), who used units of $2N$ generations. However, we see that assuming units of $2N$ generations does not lead to markedly worse population-mean PGS history estimates.

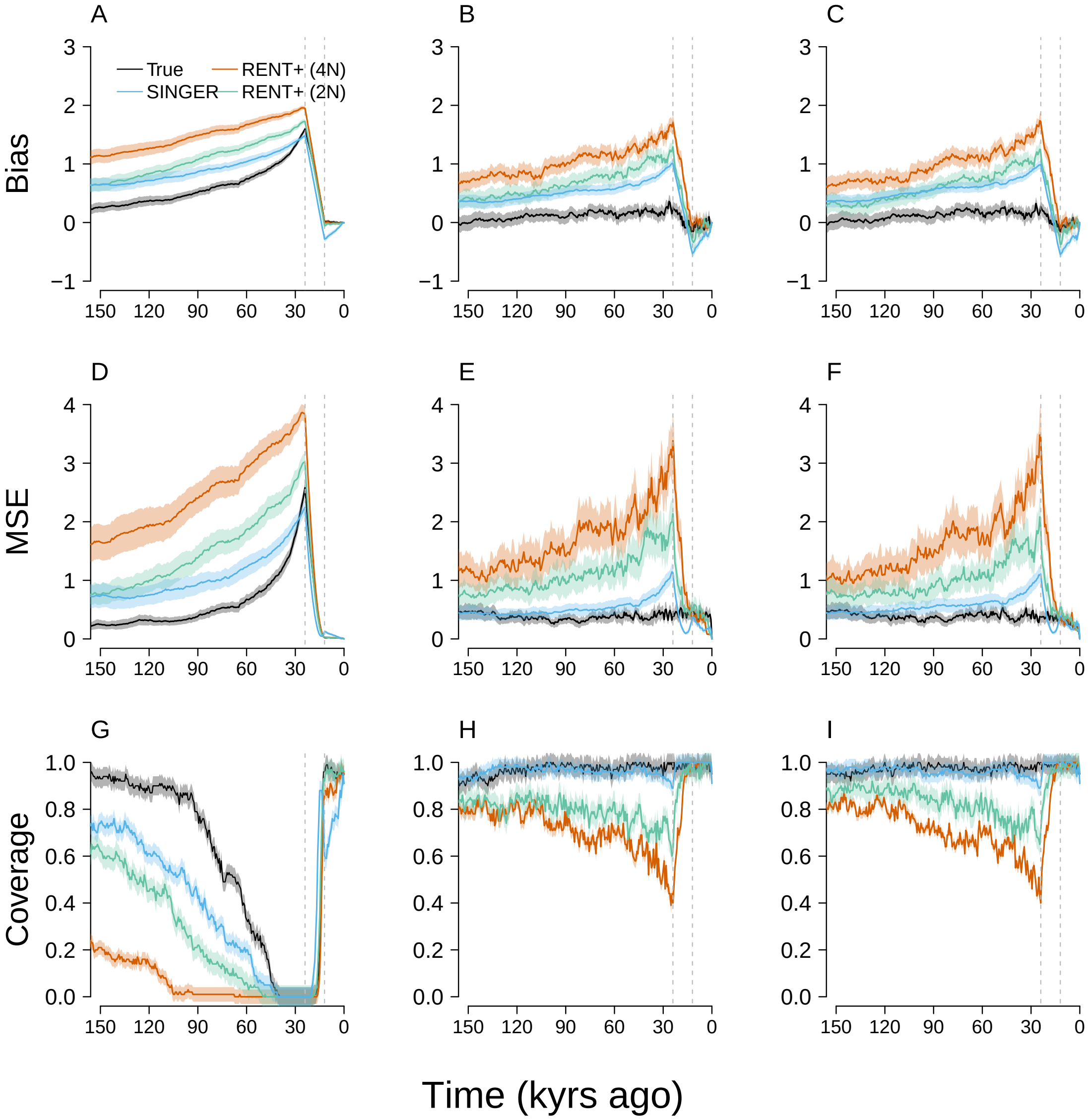


Figure S3: Performance of `RENT+` under selection. Here, we compare bias (A-C), MSE (D-F), and confidence-interval coverage (G-I) of the proportion-of-lineages (left column), waiting-time (middle column), and lineages-remaining estimators (right column), with estimates from `RENT+` trees using either $2N$ or $4N$ generations as branch length unit. In contrast to Figure S2, these simulations incorporate selection. In each simulation, the PGS was formed from 100 loci and evolved under directional selection that occurred from 0.04 to 0.02 coalescent units (units of $2N$ generations) ago. One hundred simulations were performed with 200 chromosomes.

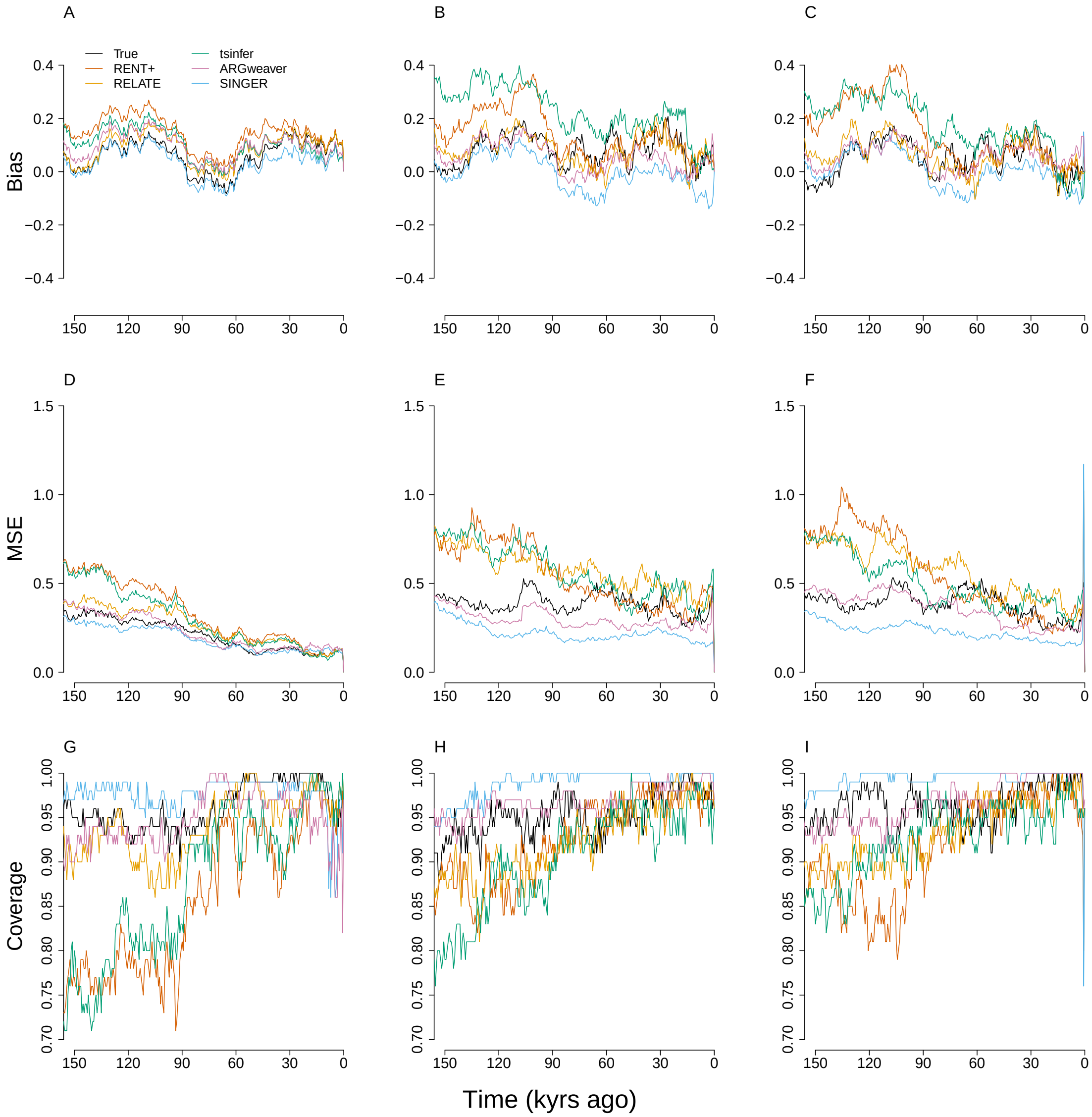


Figure S4: Performance of the methods under neutrality with samples of 20 chromosomes. Analogous to Figure 3 in the main text, we show the bias (A-C), MSE (D-F), and confidence-interval coverage (G-I) of the proportion-of-lineages (left column), waiting-time (middle column), and lineages-remaining estimators (right column), with the true trees and estimated trees from each ARG-estimation method as input under neutrality. Here, all methods are run with samples of 20 chromosomes, regardless of scalability. We also include `RENT+`, which was suppressed in the main text to simplify the presentation. Most estimated trees (including the true trees) show larger MSE and lower confidence-interval coverage when the sample size decreases to 20. `ARGweaver` and `SINGER` trees show lower MSE than true trees during some periods.

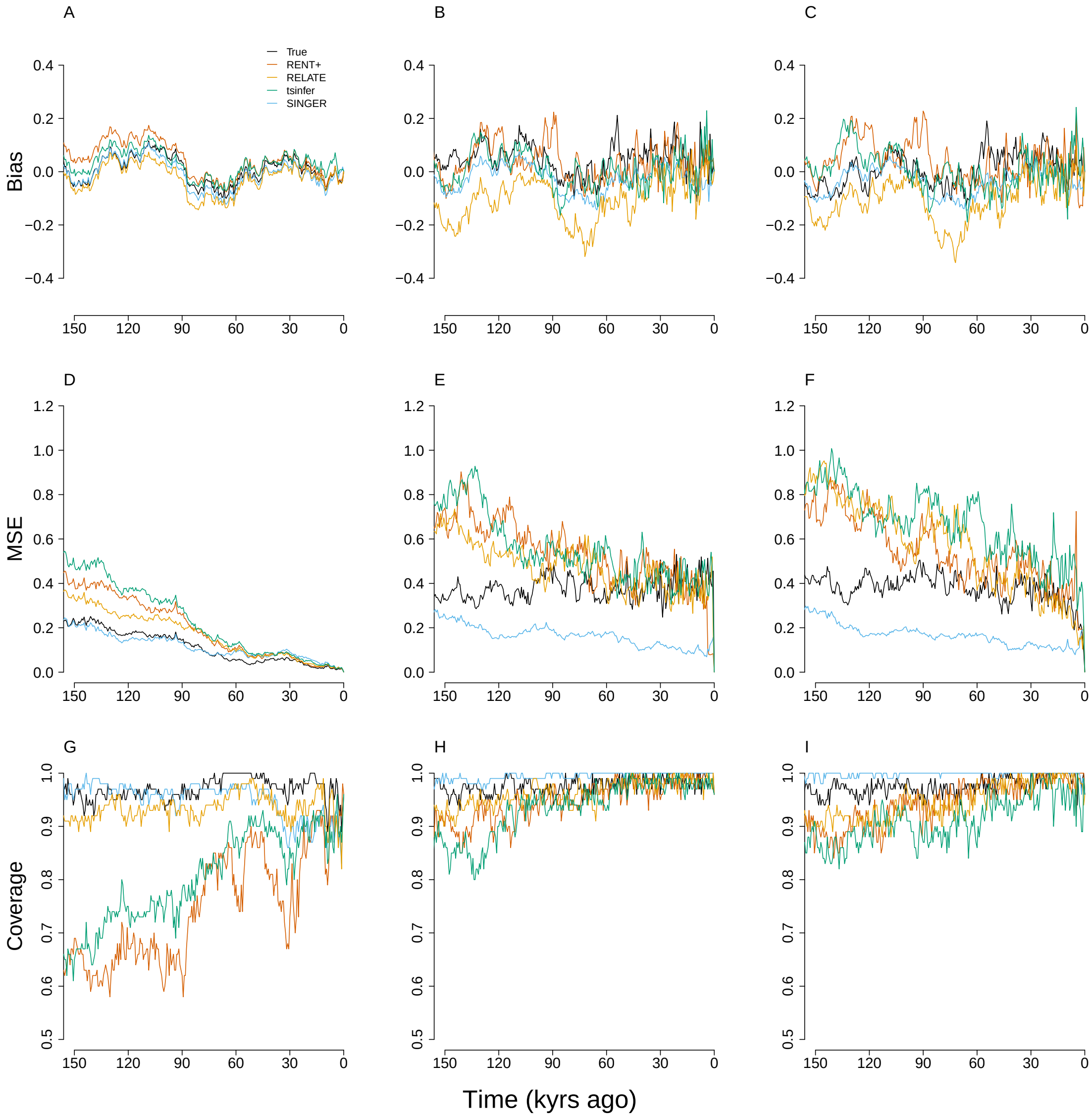


Figure S5: Performance of the methods under neutrality with samples of 200 chromosomes. Analogous to Figure 3 in the main text, we show the bias (A-C), MSE (D-F), and confidence-interval coverage (G-I) of the proportion-of-lineages (left column), waiting-time (middle column), and lineages-remaining estimators (right column), with the true trees and estimated trees from each ARG-estimation method as input under neutral evolution. Here, all methods are run with samples of 200 chromosomes, regardless of scalability. In each simulation, the PGS was formed from 100 loci and evolved neutrally. For each method, 100 simulations were performed.

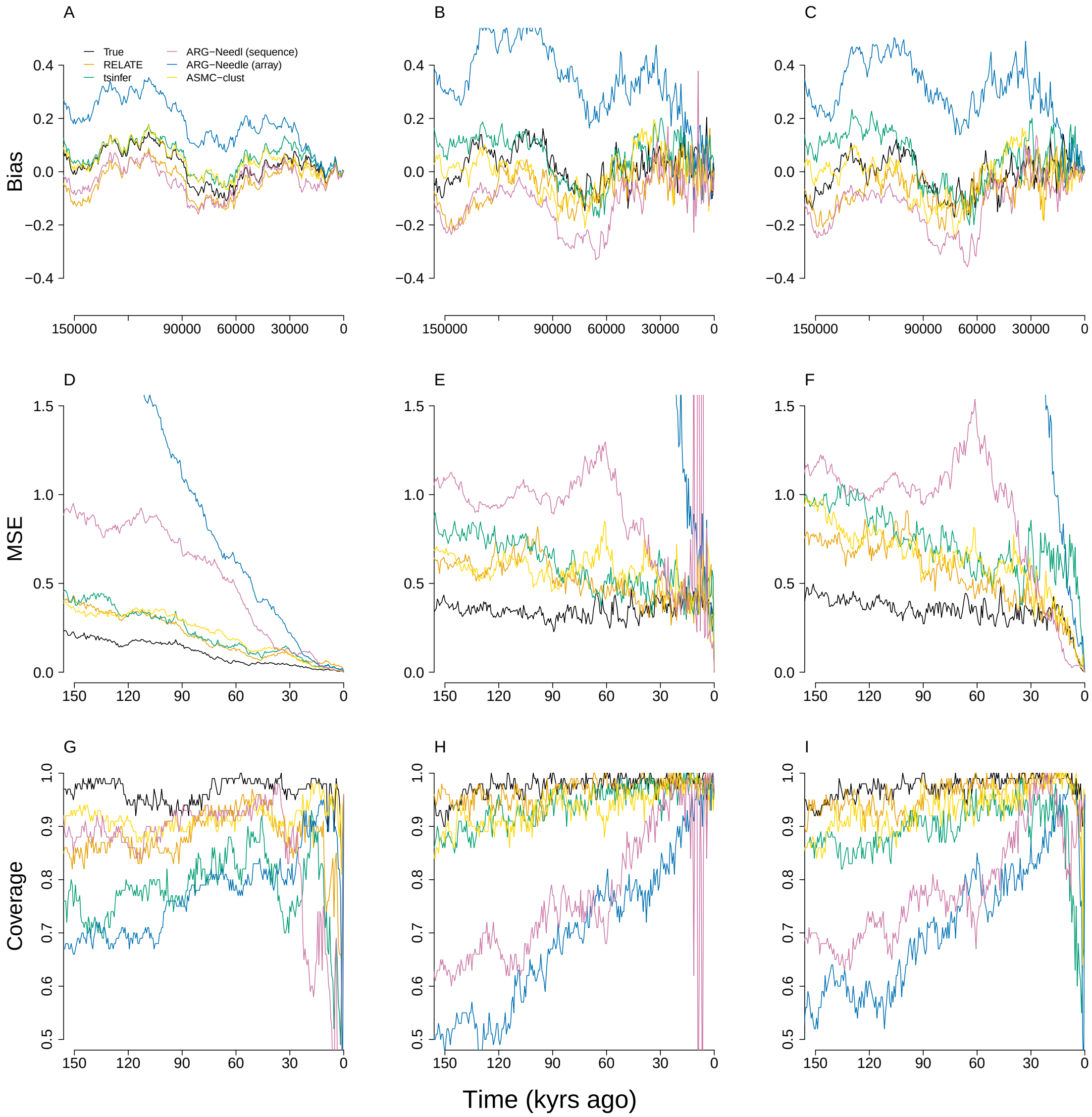


Figure S6: Performance of the methods under neutrality with samples of 2,000 chromosomes. Analogous to Figure 2 in the main text, we show bias (A-C), MSE (D-F), and confidence-interval coverage (G-I) of the proportion-of-lineages (left column), waiting-time (middle column), and lineages-remaining estimators (right column), with the true trees and estimated trees from each ARG-estimation method as input under neutral evolution. Here, all methods are run with samples of 2,000 contemporary chromosomes. In each simulation, the PGS was formed from 100 loci and evolved neutrally. For each method, 100 simulations were performed.

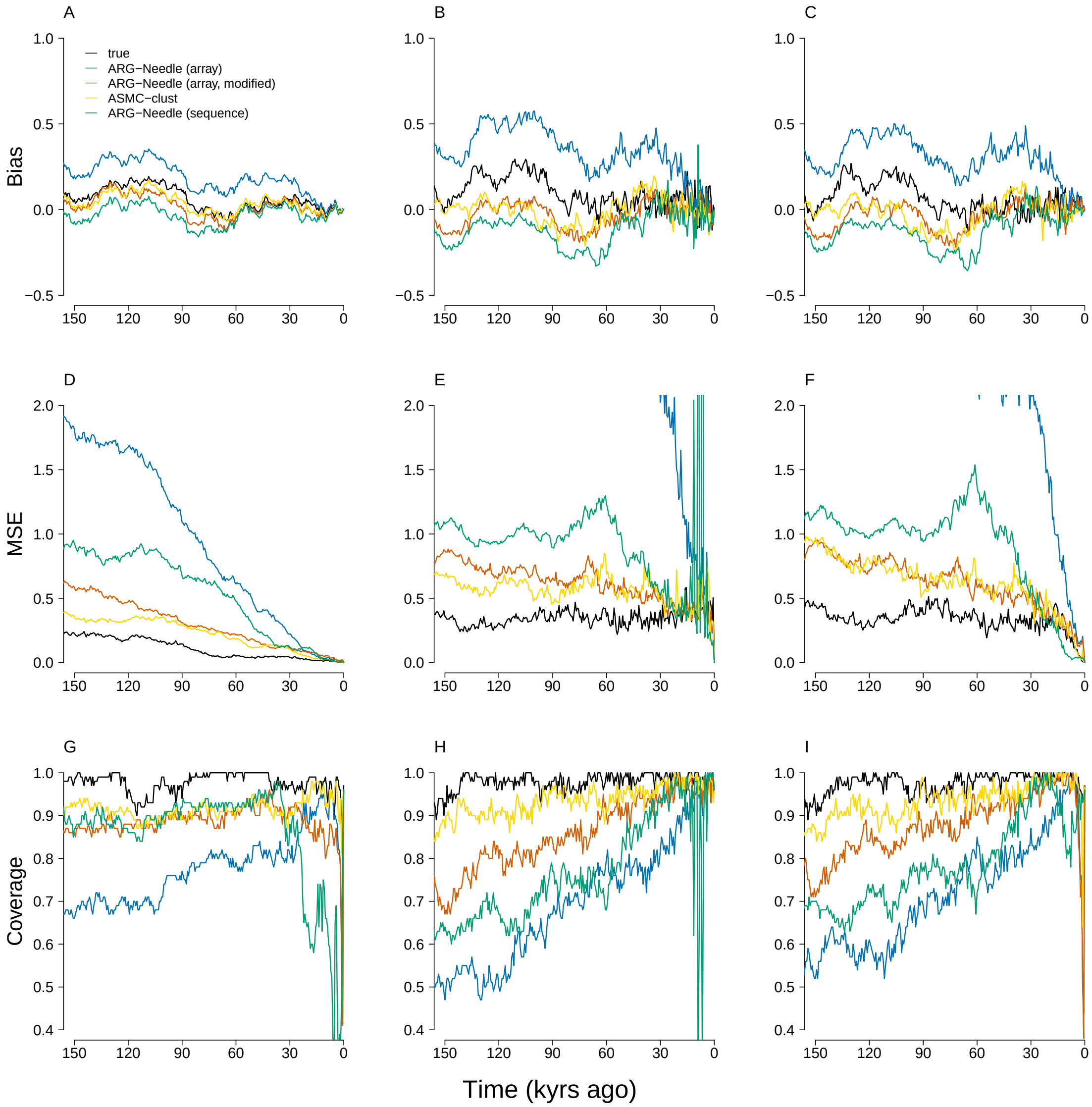


Figure S7: Performance of the methods under neutrality with a modified version of `ARG-Needle`. Here, we compare bias (A-C), MSE (D-F), and confidence-interval coverage (G-I) of the proportion-of-lineages (left column), waiting-time (middle column), and lineages-remaining estimators (right column), with the estimates from original and modified `ARG-Needle` trees. Results for true trees and `ASMC-clust` trees are shown for reference. In each simulation, the PGS is formed from 100 loci and evolved neutrally. 100 simulations were performed with 2,000 chromosomes. As described above, `ARG-Needle` was modified in order to induce the derived subtree to coalesce concurrently with the most anciently coalescing subtree of entirely derived tips.

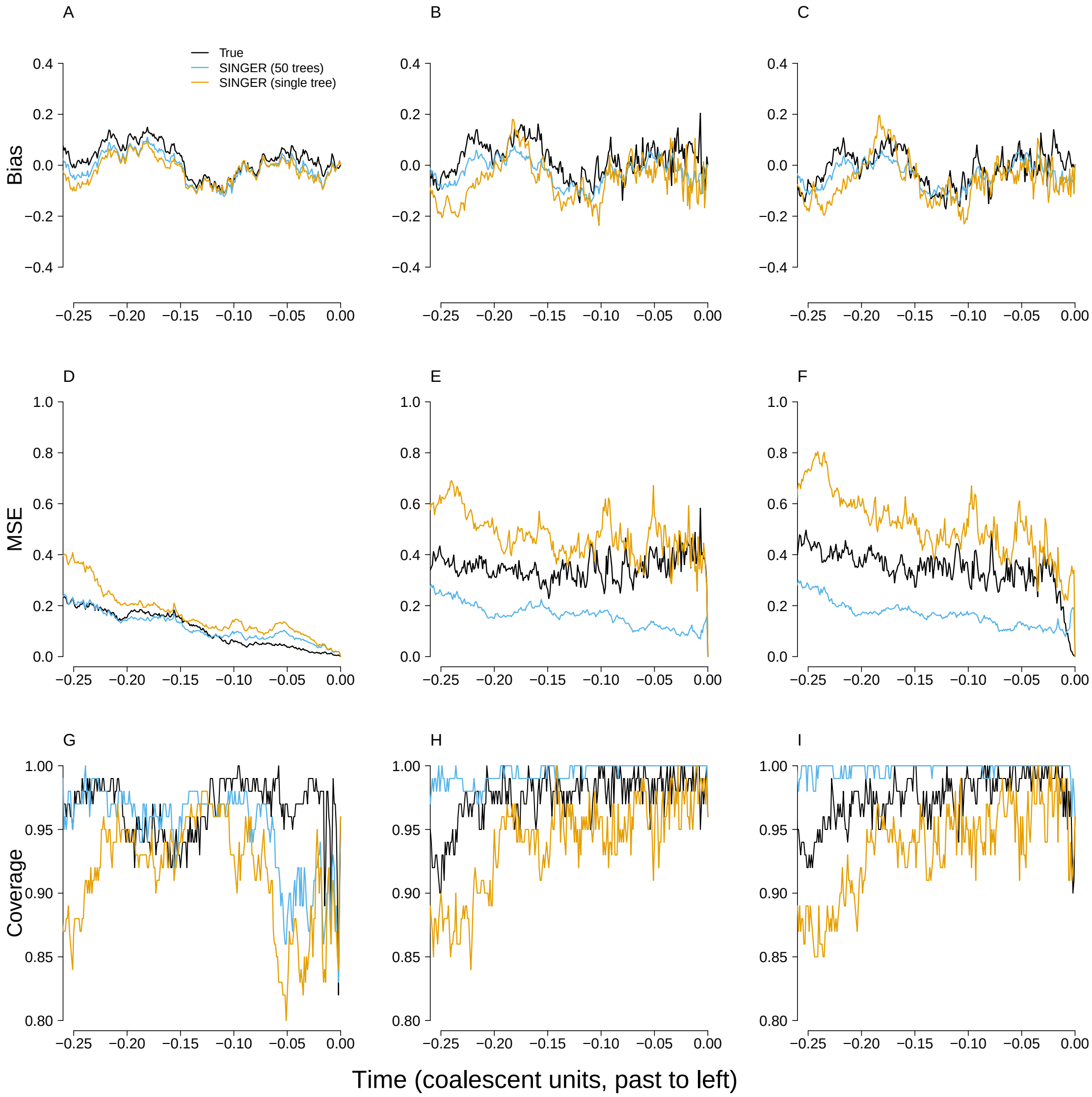


Figure S8: Performance of the methods under neutrality with a single posterior tree from SINGER, as opposed to 50 trees as used in the main text. Here, we compare bias (A-C), MSE (D-F), and confidence-interval coverage (G-I) of the proportion-of-lineages (left column), waiting-time (middle column), and lineages-remaining estimators (right column), with the estimates from one and the average of 50 SINGER trees. In each simulation, the PGS is formed from 100 loci and evolved neutrally. One hundred simulations were performed with 200 chromosomes.

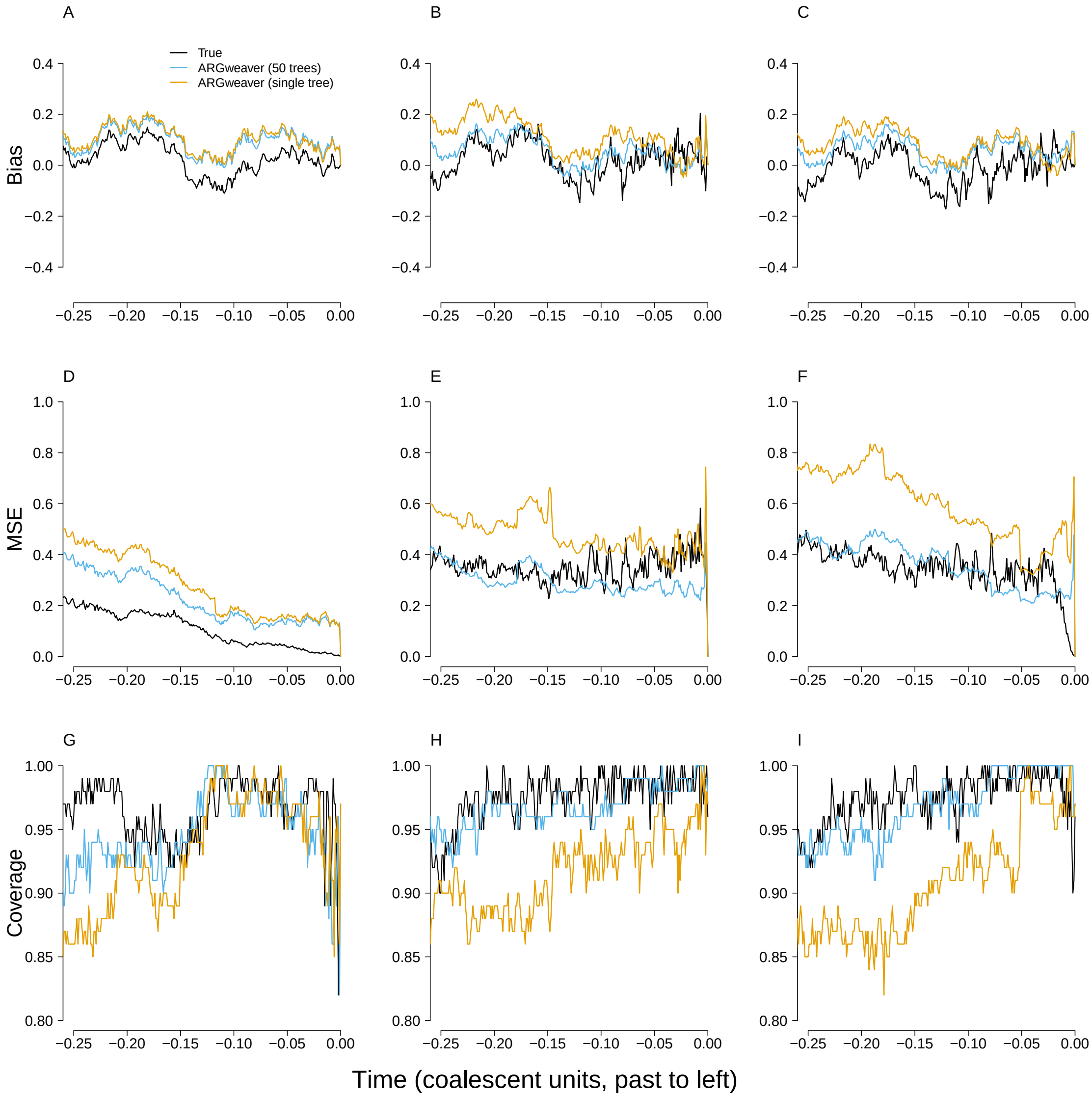


Figure S9: Performance of the methods under neutrality with a single posterior tree from `ARGweaver`, as opposed to 50 trees as used in the main text. Here, we compare bias (A-C), MSE (D-F), and confidence-interval coverage (G-I) of the proportion-of-lineages (left column), waiting-time (middle column), and lineages-remaining estimators (right column), with the estimates from one and the average of 50 `ARGweaver` trees. In each simulation, the PGS is formed from 100 loci and evolved neutrally. One hundred simulations were performed with 200 chromosomes.

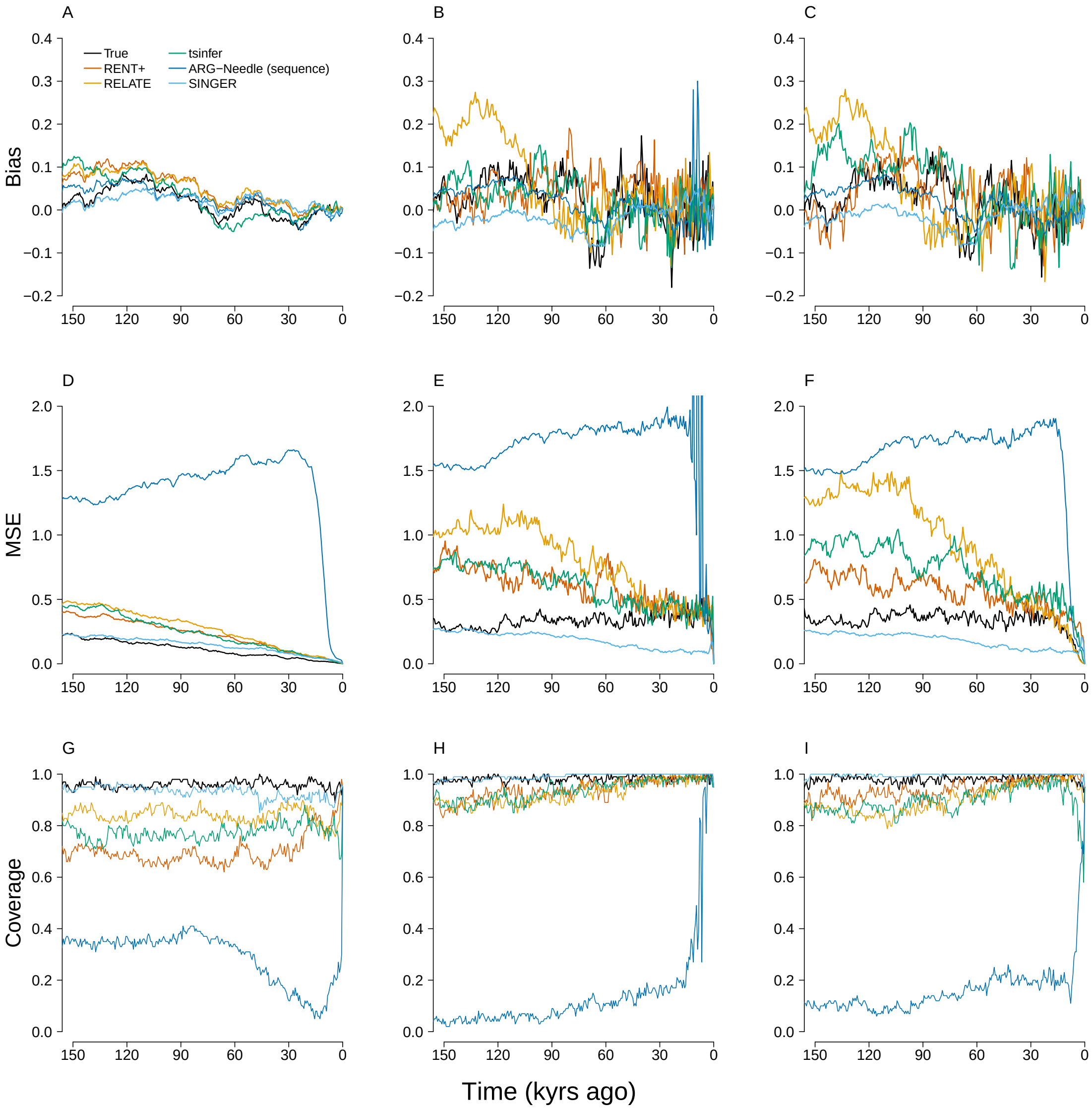


Figure S10: Results for a larger flanking region used for ARG estimation. The bias (A-C), MSE (D-F), and confidence-interval coverage (G-I) of the proportion-of-lineages (left column), waiting-time (middle column), and lineages-remaining estimators (right column), with the true trees and estimated trees from each ARG-estimation method as input. For the estimates computed from true coalescent trees (black lines), 100 simulations were performed with a sample of 2,000 chromosomes. In each simulation, the PGS was formed from 100 loci and evolved neutrally. In the main text, all ARG-estimation methods are run on the basis of a 200kb region centered on the causal locus, whereas here, the flanking region is 500kb. `Relate`, `tsinfer`, and `ASMC-clust` were used to reconstruct trees with 2,000 chromosomes. `SINGER` and `RENT+` were run with 200 chromosomes.

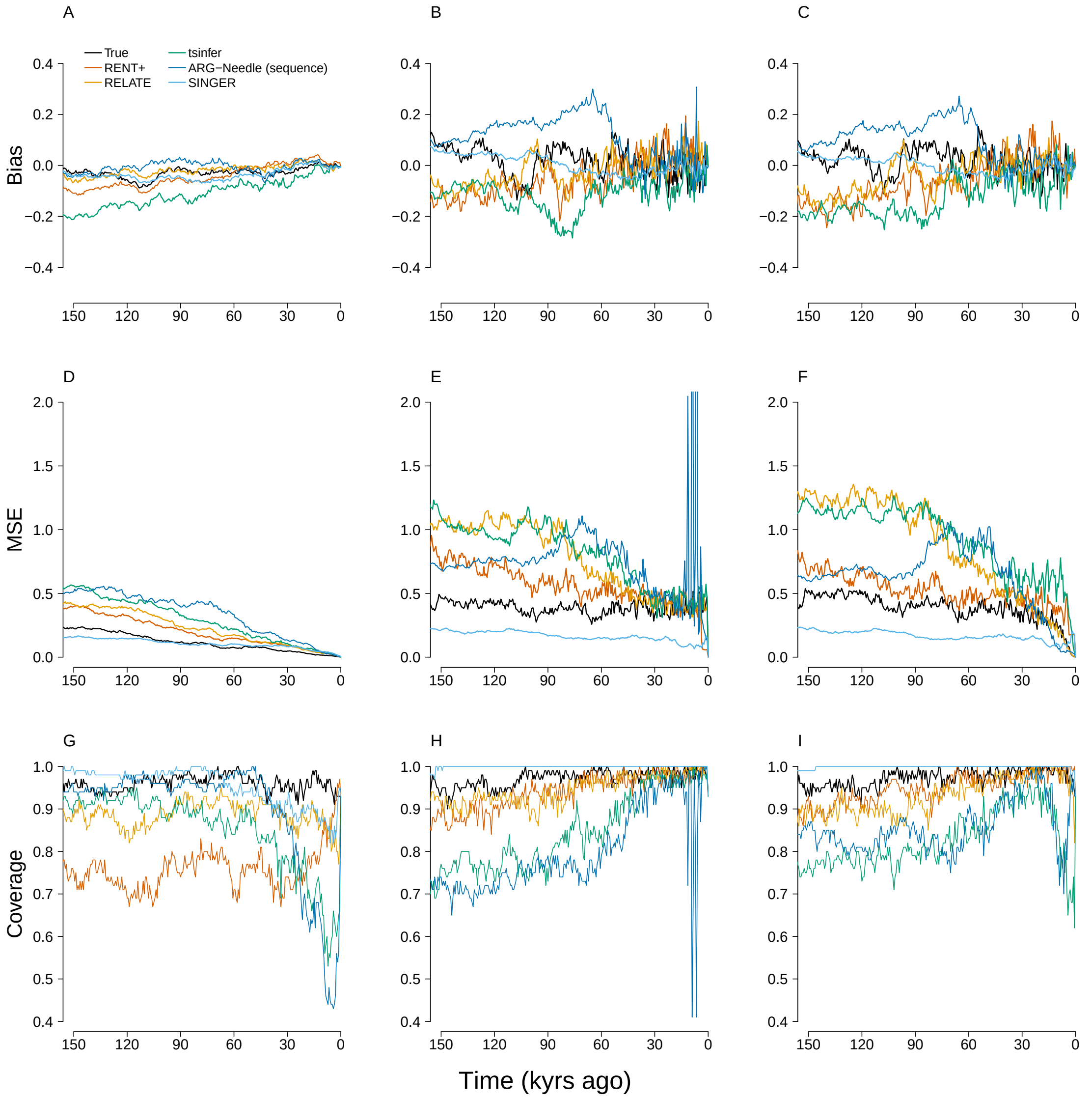


Figure S11: Simulation results including a CEU demography under neutrality. The bias (A-C), MSE (D-F), and confidence-interval coverage (G-I) of the proportion-of-lineages (left column), waiting-time (middle column), and lineages-remaining estimators (right column), with the true trees and estimated trees from each ARG-estimation method as input. For the estimates computed from true coalescent trees (black lines), 100 simulations were performed with a sample of 2,000 chromosomes and CEU demographic data. In each simulation, the PGS was formed from 100 loci and evolved neutrally. `Relate`, `tsinfer`, and `ASMC-clust` were used to reconstruct trees with 2,000 chromosomes. `SINGER` and `RENT+` were run with 200 chromosomes. Lines from estimated trees are based on 100 simulations.

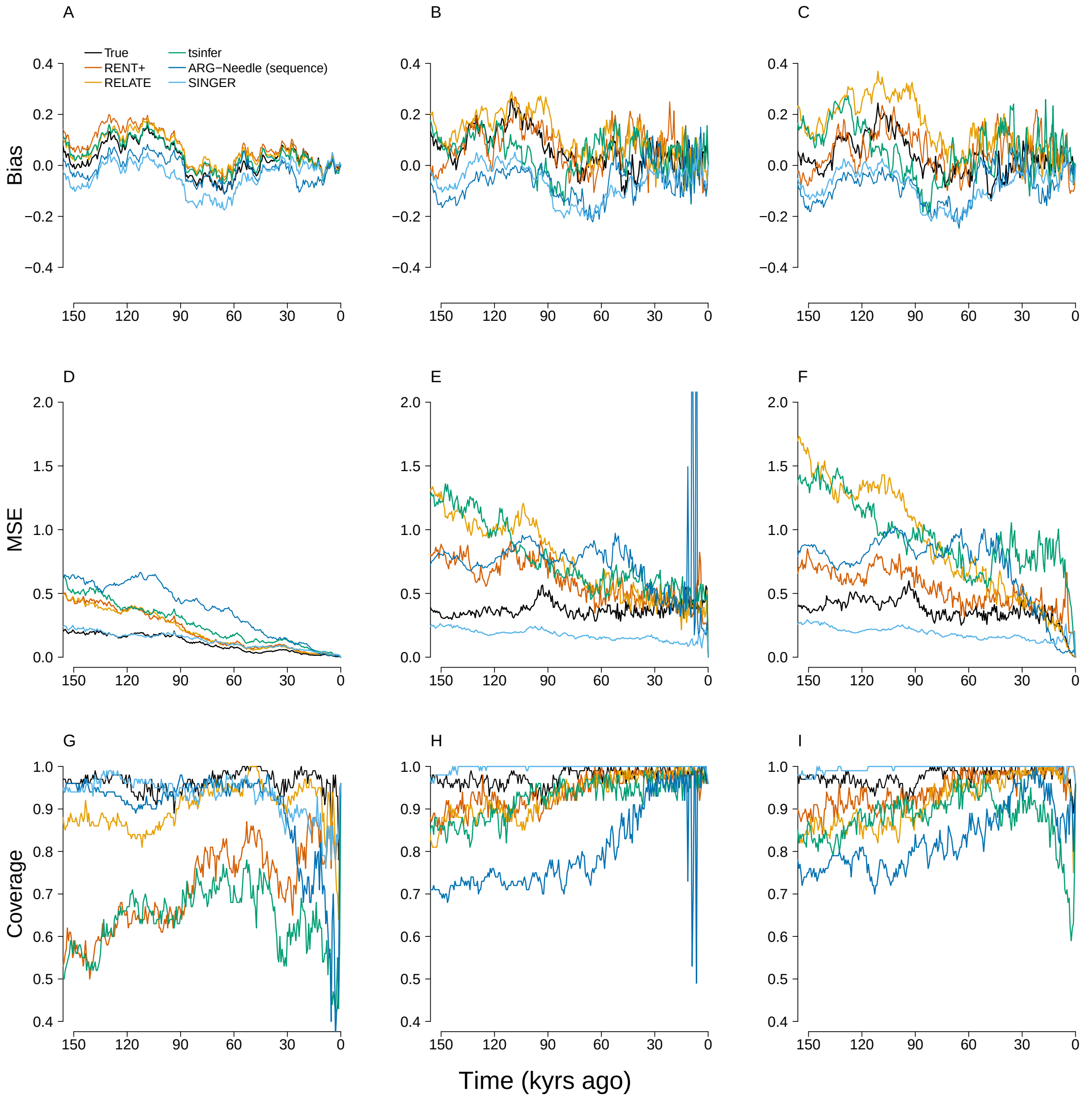


Figure S12: Simulation results including genotyping error with 0.1% rate under neutrality. The bias (A-C), MSE (D-F), and confidence-interval coverage (G-I) of the proportion-of-lineages (left column), waiting-time (middle column), and lineages-remaining estimators (right column), with the true trees and estimated trees from each ARG-estimation method as input. For the estimates computed from true coalescent trees (black lines), 100 simulations were performed with a sample of 2,000 chromosomes. In each simulation, the PGS was formed from 100 loci and evolved neutrally. The haplotypes input to ARG estimated tools were simulated with genotyping error. `Relate`, `tsinfer`, and `ASMC-clust` were used to reconstruct trees with 2,000 chromosomes. `SINGER` and `RENT+` were run with 200 chromosomes. Lines from estimated trees are based on 100 simulations.

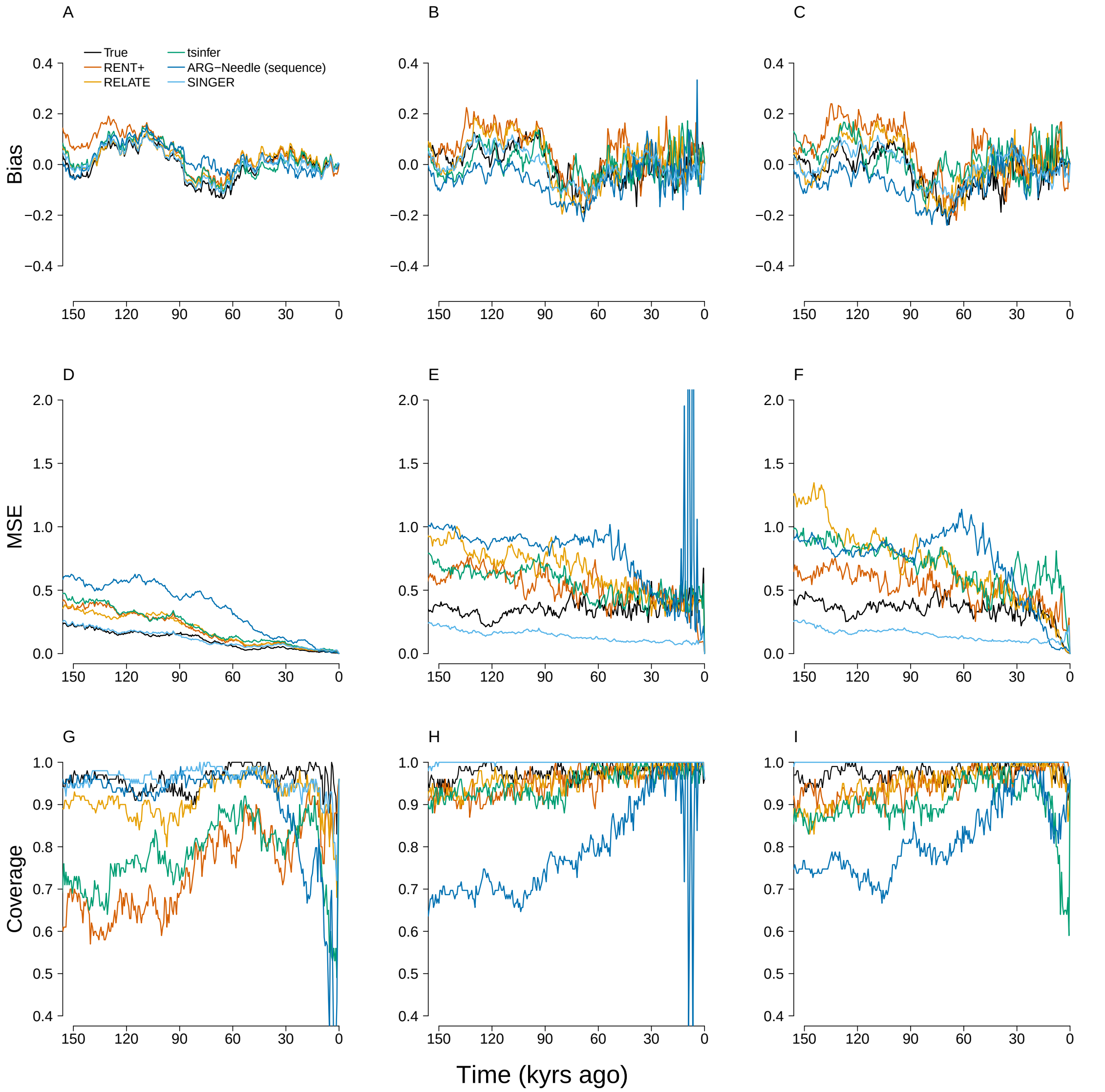


Figure S13: Simulation results including phasing error under neutrality. The bias (A-C), MSE (D-F), and confidence-interval coverage (G-I) of the proportion-of-lineages (left column), waiting-time (middle column), and lineages-remaining estimators (right column), with the true trees and estimated trees from each ARG-estimation method as input. For the estimates computed from true coalescent trees (black lines), 100 simulations were performed with a sample of 2,000 chromosomes. In each simulation, the PGS was formed from 100 loci and evolved neutrally. The haplotypes input to ARG estimated tools were simulated with phasing error. `Relate`, `tsinfer`, and `ASMC-clust` were used to reconstruct trees with 2,000 chromosomes. `SINGER` and `RENT+` were run with 200 chromosomes. Lines from estimated trees are based on 100 simulations.

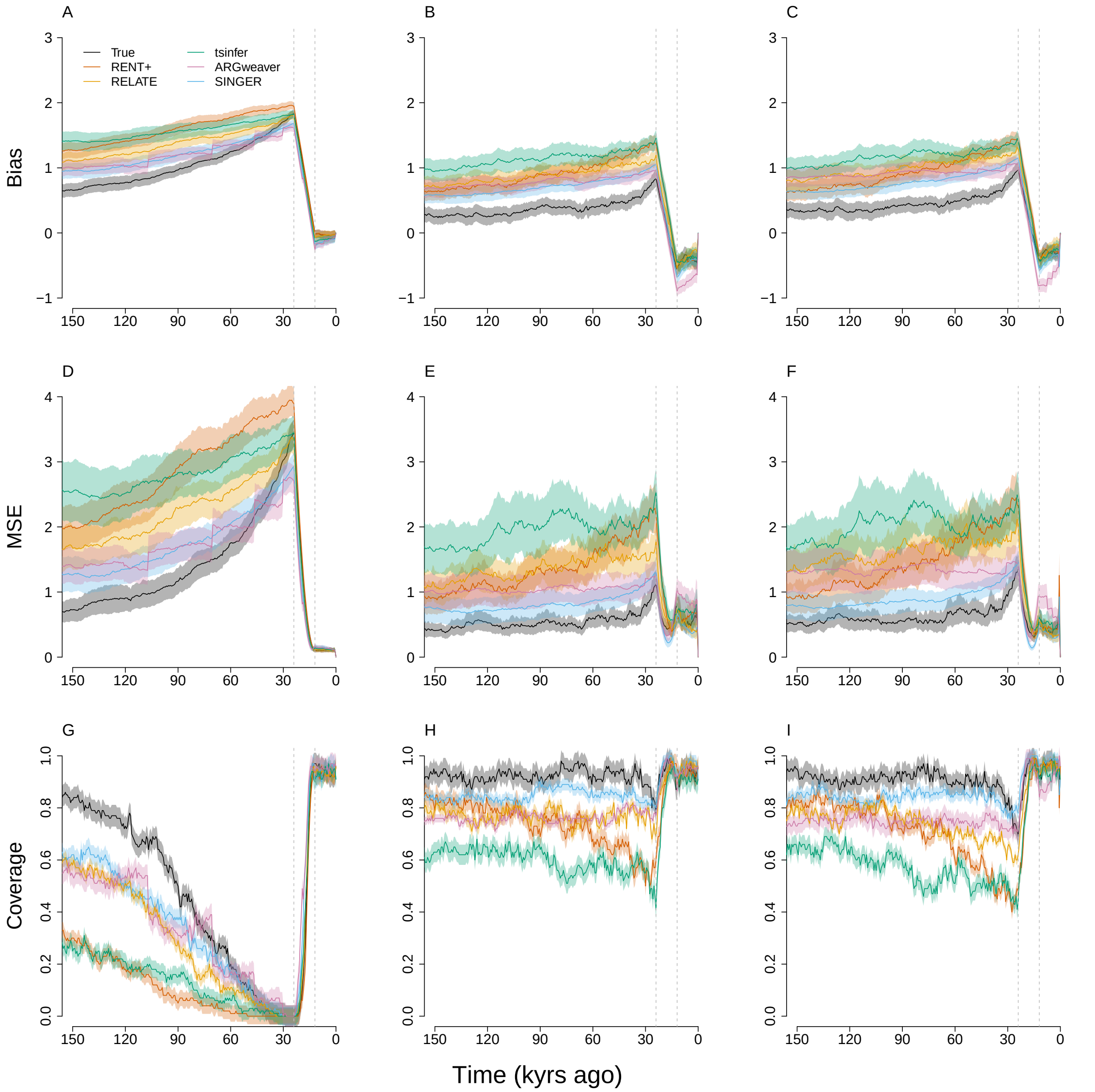


Figure S14: Performance of the methods under selection with samples of 20 chromosomes. Analogous to Figure 4 in the main text, we show the bias (A-C), MSE (D-F), and confidence-interval coverage (G-I) of the proportion-of-lineages (left column), waiting-time (middle column), and lineages-remaining estimators (right column), with the true trees and estimated trees from each ARG-estimation method as input under directional selection. In each simulation, the PGS was formed from 100 loci and evolved neutrally except between 0.02 and 0.04 coalescent units ago (units of $2N$ generations), during which time it was under directional selection. For each method, 100 simulations were performed with 20 chromosomes.

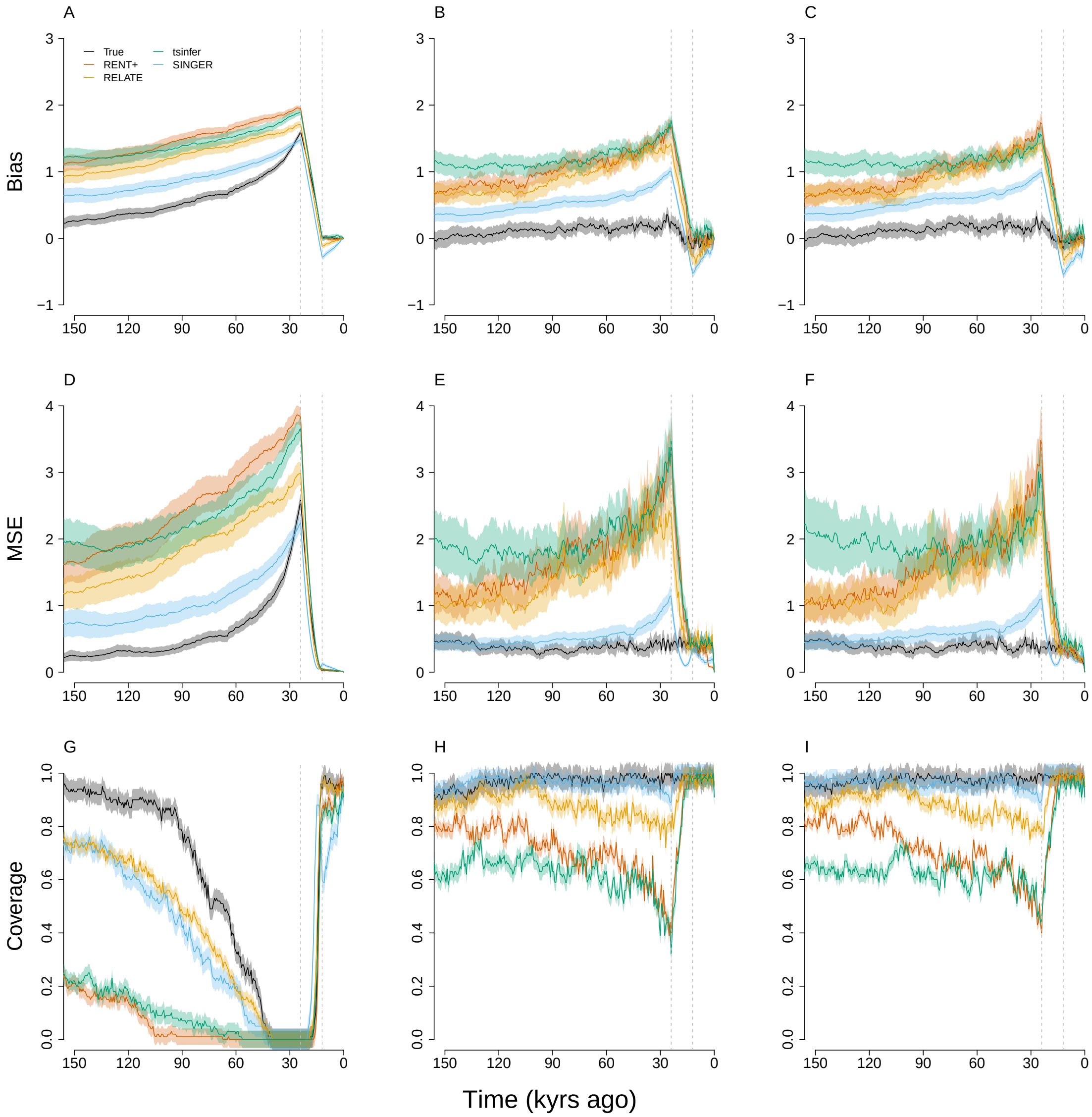


Figure S15: Performance of the methods under selection with samples of 200 chromosomes. The bias (A-C), MSE (D-F), and confidence-interval coverage (G-I) of the proportion-of-lineages (left column), waiting-time (middle column), and lineages-remaining estimators (right column), with the true trees and estimated trees from each ARG-estimation method as input under directional selection. In each simulation, the PGS was formed from 100 loci and evolved under directional selection between 0.02 and 0.04 coalescent units ago. For each method, 100 simulations were performed with 200 chromosomes.

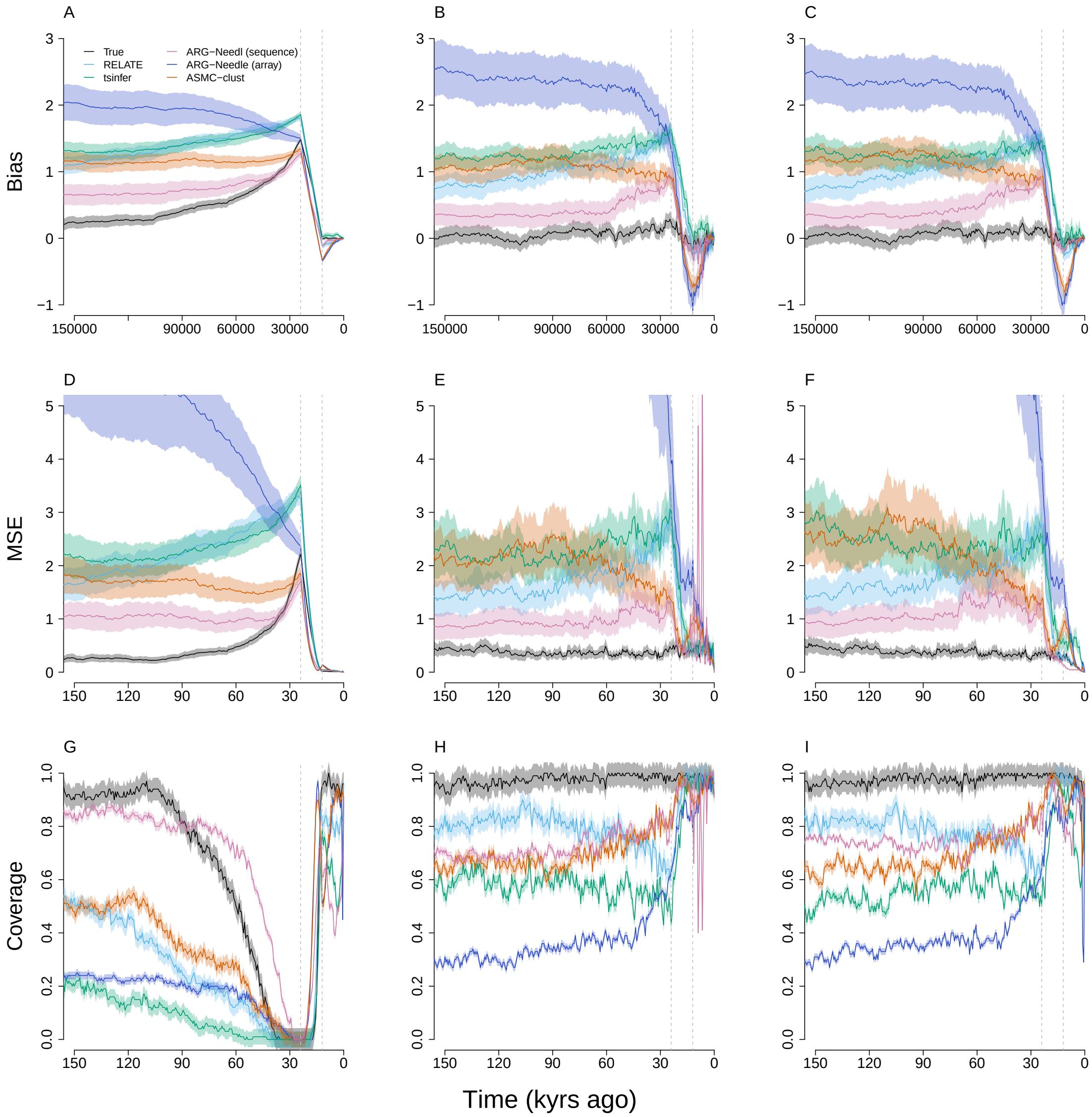


Figure S16: Performance of the methods under selection with samples of 2,000 chromosomes. The bias (A-C), MSE (D-F), and confidence-interval coverage (G-I) of the proportion-of-lineages (left column), waiting-time (middle column), and lineages-remaining estimators (right column), with the true trees and estimated trees from each ARG-estimation method as input under directional selection. In each simulation, the PGS was formed from 100 loci and evolved under directional selection between 0.02 and 0.04 coalescent units ago. For each method, 100 simulations were performed with 2,000 chromosomes.

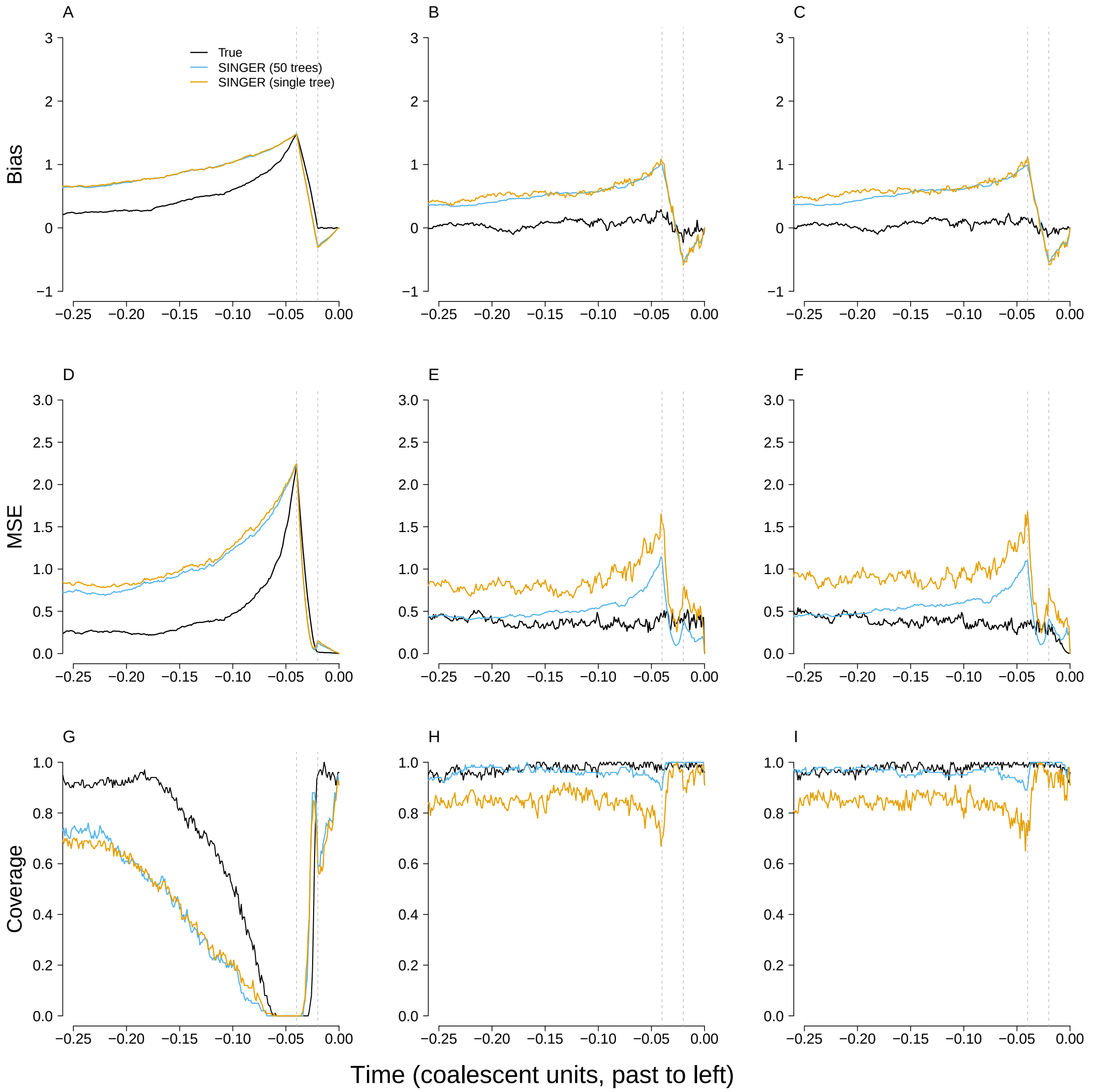


Figure S17: Performance of the methods under directional selection with a single posterior tree from SINGER, as opposed to 50 trees as used in the main text. Here, we compare bias (A-C), MSE (D-F), and confidence-interval coverage (G-I) of the proportion-of-lineages (left column), waiting-time (middle column), and lineages-remaining estimators (right column), with the estimates from one and 50 SINGER trees. In each simulation, the PGS is formed from 100 loci and evolved under directional selection between 0.02 and 0.04 coalescent units ago. One hundred simulations were performed with 200 chromosomes each.

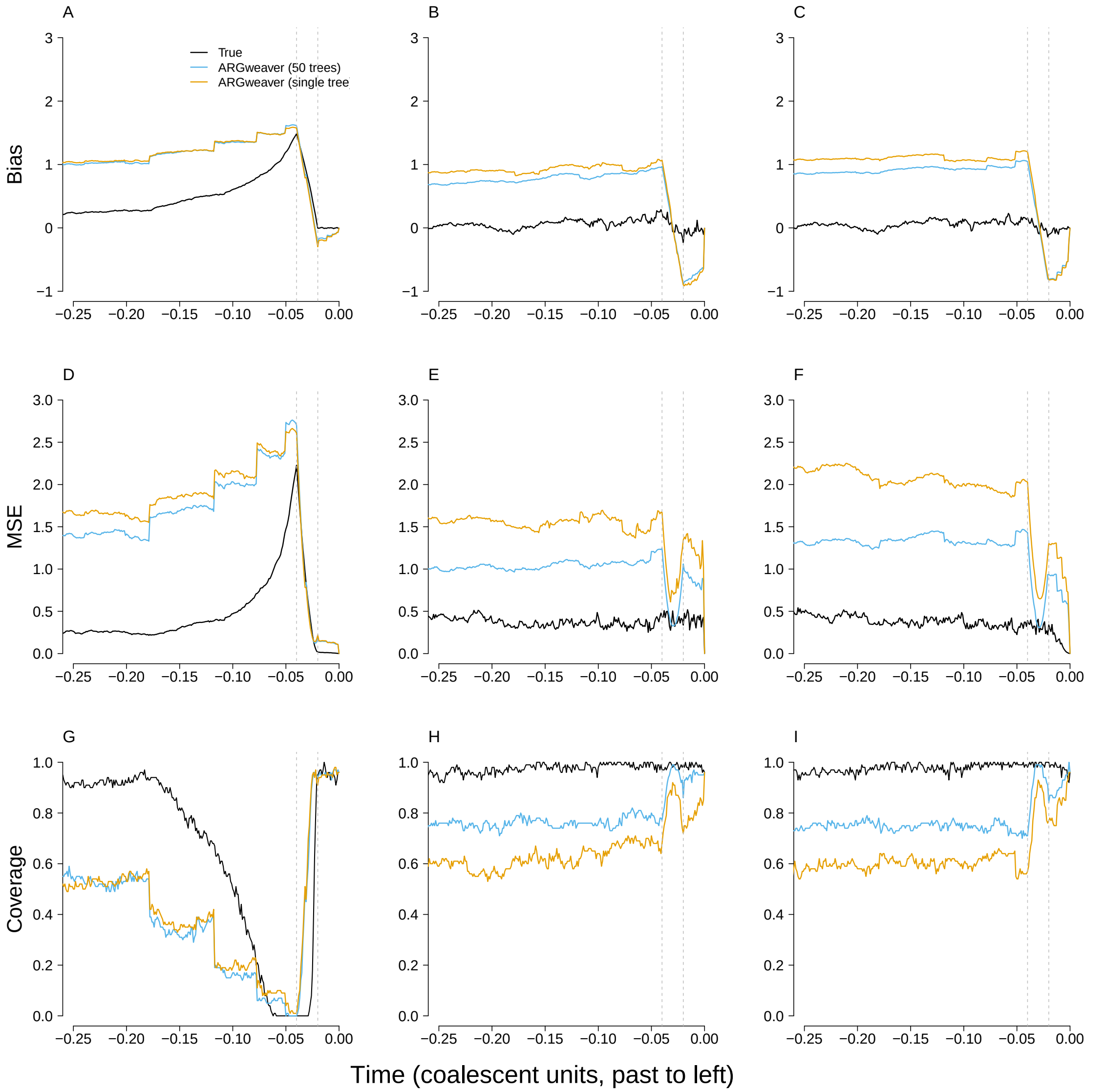


Figure S18: Performance of the methods under directional selection with a single posterior tree from `ARGweaver`, as opposed to 50 trees as used in the main text. Here, we compare bias (A-C), MSE (D-F), and confidence-interval coverage (G-I) of the proportion-of-lineages (left column), waiting-time (middle column), and lineages-remaining estimators (right column), with the estimates from one and 50 `ARGweaver` trees. In each simulation, the PGS is formed from 100 loci and evolved under directional selection between 0.02 and 0.04 coalescent units ago. 100 simulations were performed with 20 chromosomes.

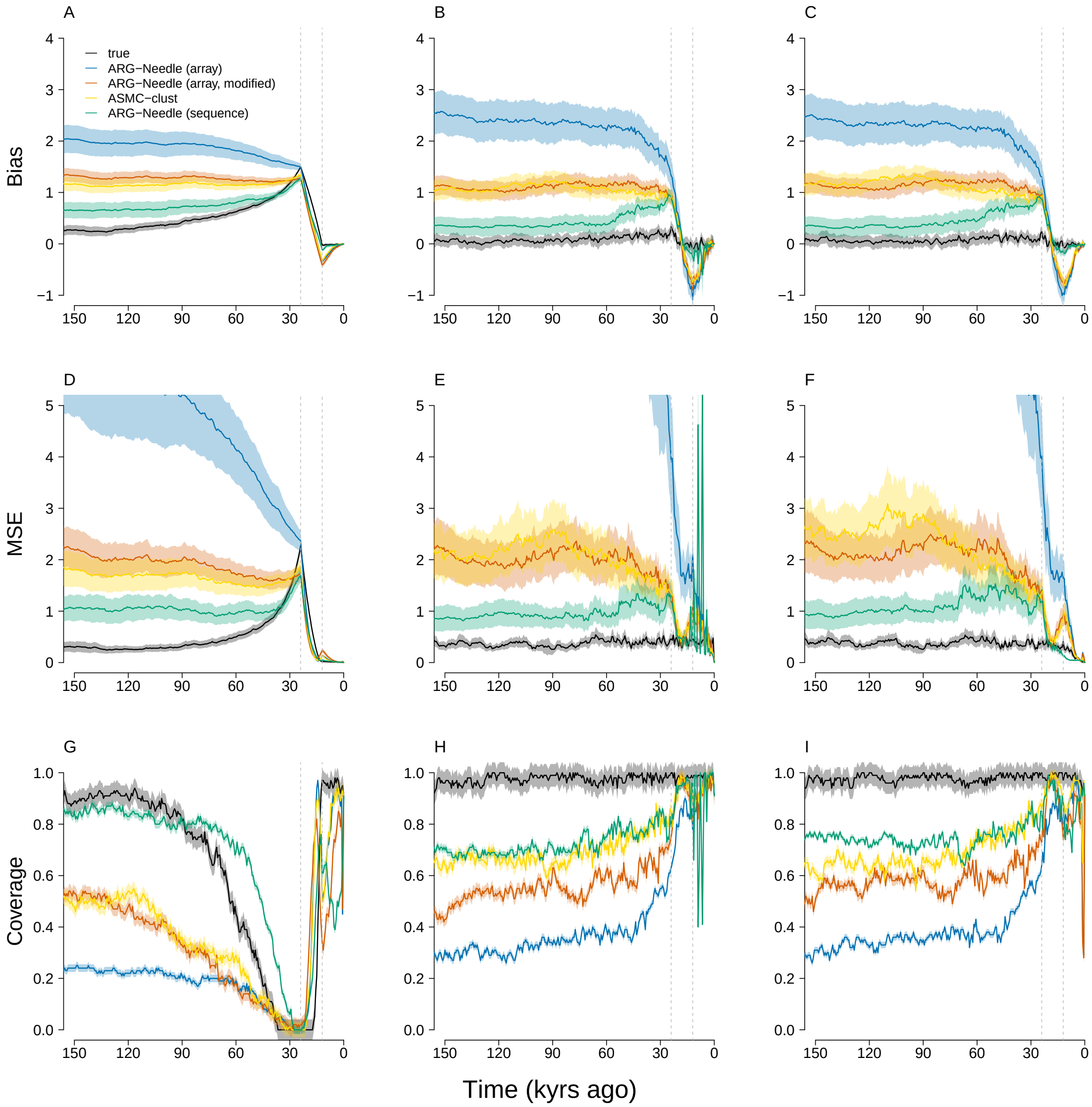


Figure S19: Performance of the estimators with a modified version of `ARG-Needle` under selection. Here, we compare bias (A-C), MSE (D-F), and confidence-interval coverage (G-I) of the proportion-of-lineages (left column), waiting-time (middle column), and lineages-remaining estimators (right column), with the estimates from original and modified `ARG-Needle` trees. In each simulation, the PGS is formed from 100 loci and evolved under directional selection. 100 simulations were performed with 2,000 chromosomes. As described, `ARG-Needle` was modified in order to induce the derived subtree to coalesce concurrently with the most anciently coalescing subtree of entirely derived tips.

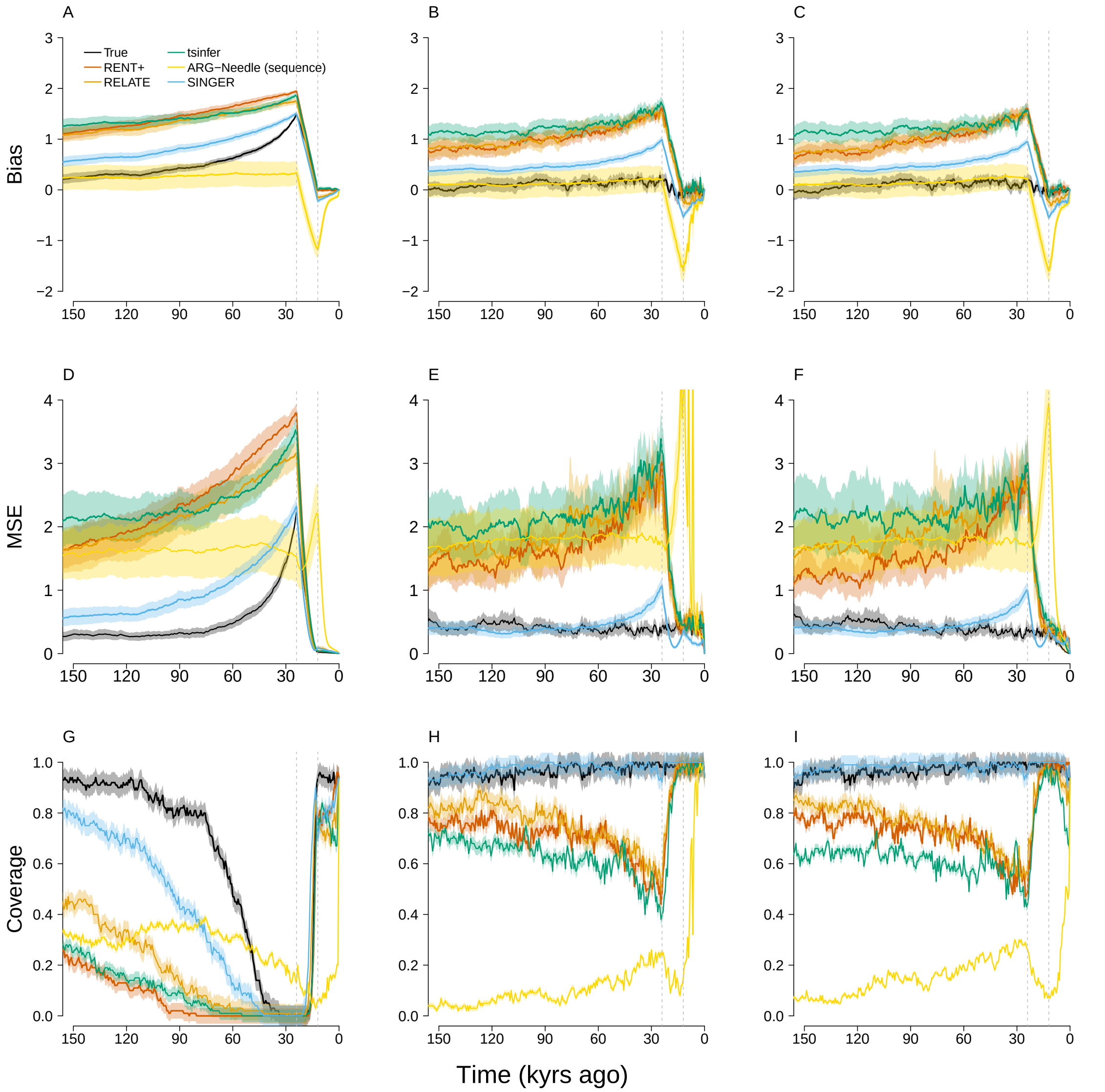


Figure S20: Simulation results with 500,000 bp haplotypes. The bias (A-C), MSE (D-F), and confidence-interval coverage (G-I) of the proportion-of-lineages (left column), waiting-time (middle column), and lineages-remaining estimators (right column), with the true trees and estimated trees from each ARG-estimation method as input. For the estimates computed from true coalescent trees (black lines), 100 simulations were performed with a sample of 2,000 chromosomes. In each simulation, the PGS was formed from 100 loci and evolved under directional selection between 0.02 and 0.04 coalescent units ago. The length of region surrounding each causal locus for ARG estimation is 500,000 base pairs. `Relate`, `tsinfer`, and `ASMC-clust` were used to reconstruct trees with 2,000 chromosomes. `SINGER` and `RENT+` were run with 200 chromosomes. Lines from estimated trees are based on 100 simulations.

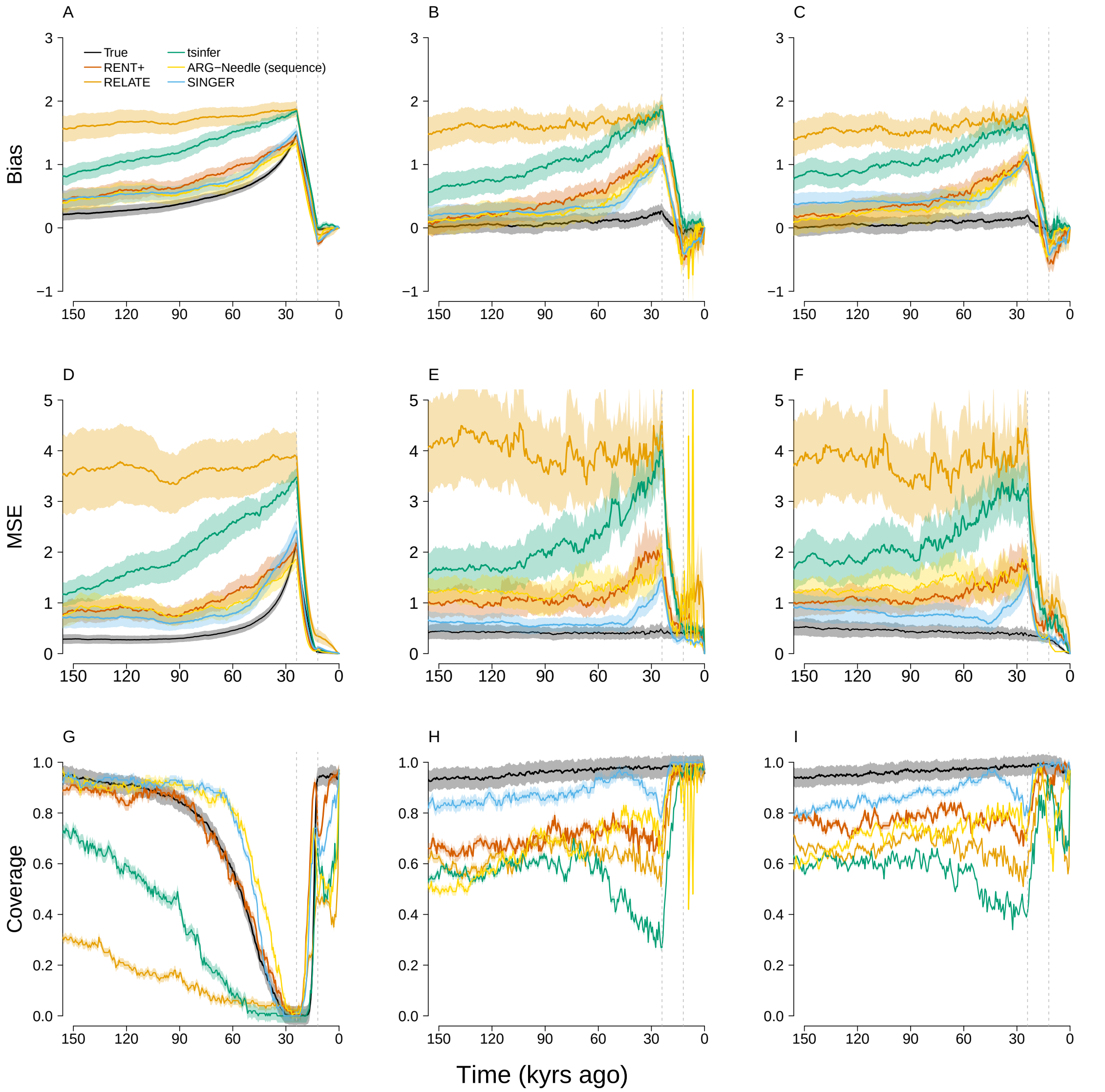


Figure S21: Simulation results including a CEU demography under directional selection. The bias (A-C), MSE (D-F), and confidence-interval coverage (G-I) of the proportion-of-lineages (left column), waiting-time (middle column), and lineages-remaining estimators (right column), with the true trees and estimated trees from each ARG-estimation method as input. For the estimates computed from true coalescent trees (black lines), 100 simulations were performed with a sample of 2,000 chromosomes and CRU demographic data. In each simulation, the PGS was formed from 100 loci and evolved under directional selection between 0.02 and 0.04 coalescent units ago. `Relate`, `tsinfer`, and `ASMC-clust` were used to reconstruct trees with 2,000 chromosomes. `SINGER` and `RENT+` were run with 200 chromosomes. Lines from estimated trees are based on 100 simulations.

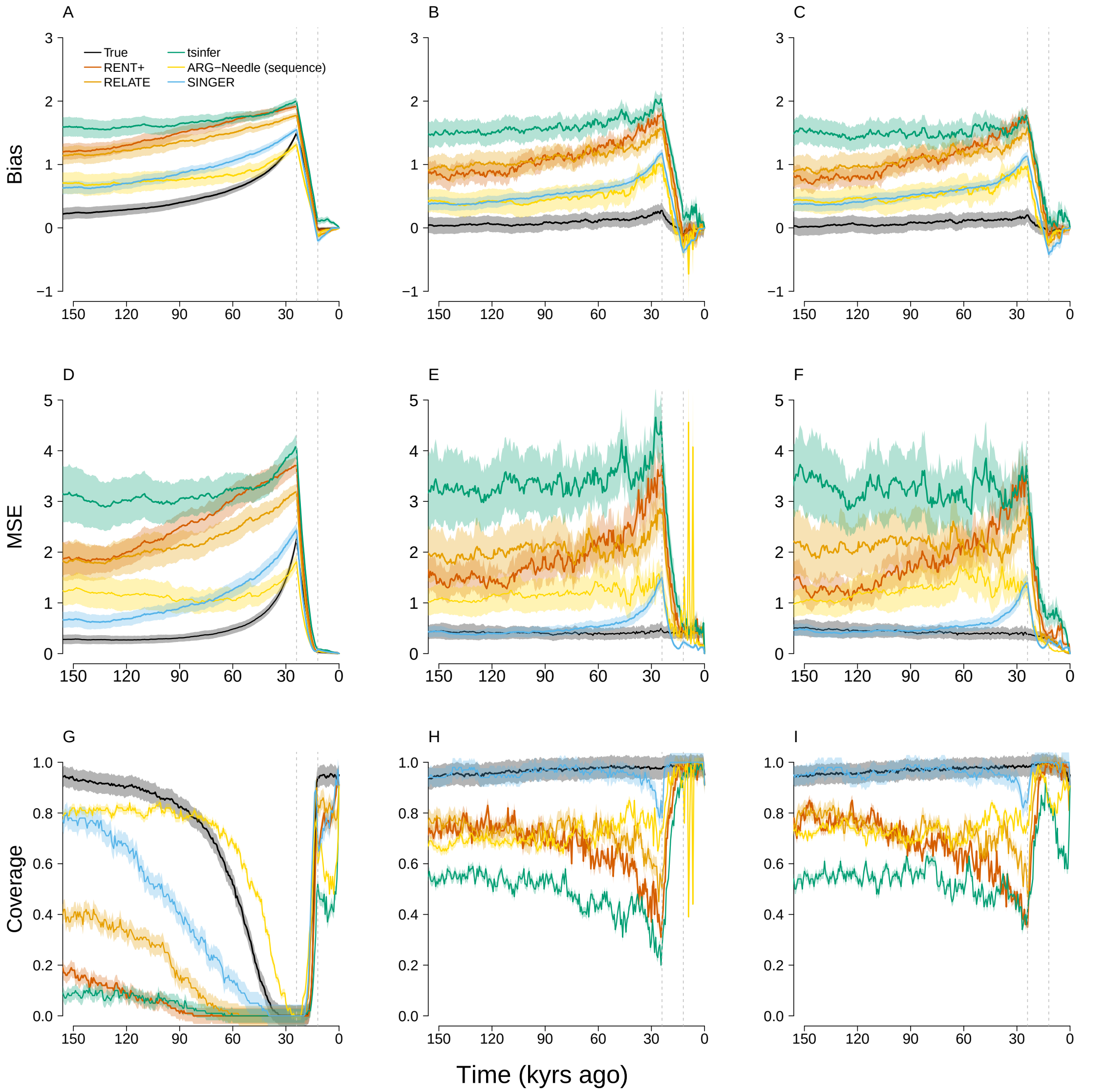


Figure S22: Simulation results including genotyping error with 0.1% rate under directional selection. The bias (A-C), MSE (D-F), and confidence-interval coverage (G-I) of the proportion-of-lineages (left column), waiting-time (middle column), and lineages-remaining estimators (right column), with the true trees and estimated trees from each ARG-estimation method as input. For the estimates computed from true coalescent trees (black lines), 100 simulations were performed with a sample of 2,000 chromosomes. In each simulation, the PGS was formed from 100 loci and evolved under directional selection between 0.02 and 0.04 coalescent units ago. The haplotypes input to ARG estimated tools were simulated with genotyping error. `Relate`, `tsinfer`, and `ASMC-clust` were used to reconstruct trees with 2,000 chromosomes. `SINGER` and `RENT+` were run with 200 chromosomes. Lines from estimated trees are based on 100 simulations.

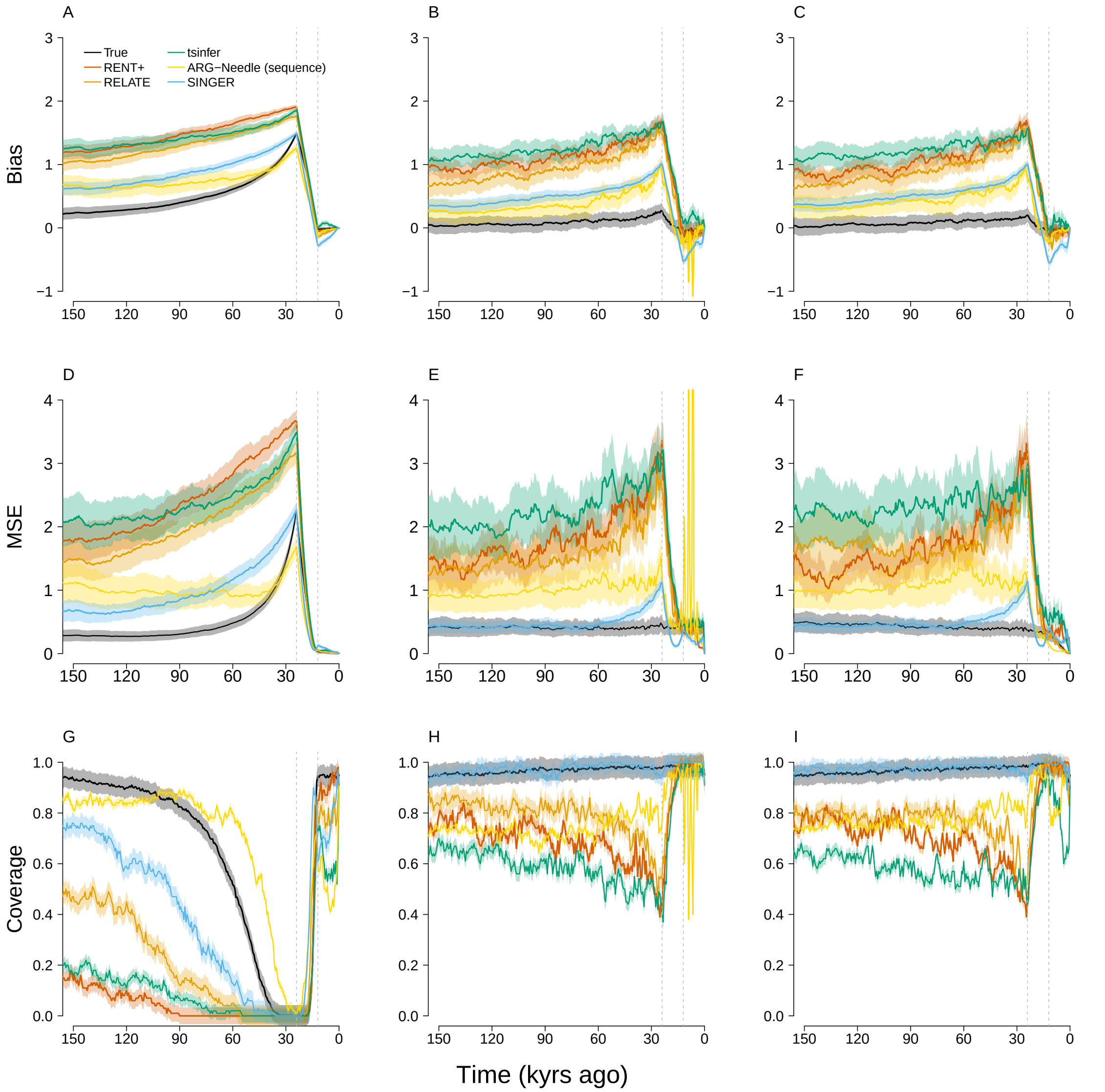


Figure S23: Simulation results including phasing error under directional selection. The bias (A-C), MSE (D-F), and confidence-interval coverage (G-I) of the proportion-of-lineages (left column), waiting-time (middle column), and lineages-remaining estimators (right column), with the true trees and estimated trees from each ARG-estimation method as input. For the estimates computed from true coalescent trees (black lines), 100 simulations were performed with a sample of 2,000 chromosomes. In each simulation, the PGS was formed from 100 loci and evolved under directional selection between 0.02 and 0.04 coalescent units ago. The haplotypes input to ARG estimated tools were simulated with phasing error. `Relate`, `tsinfer`, and `ASMC-clust` were used to reconstruct trees with 2,000 chromosomes. `SINGER` and `RENT+` were run with 200 chromosomes. Lines from estimated trees are based on 100 simulations.

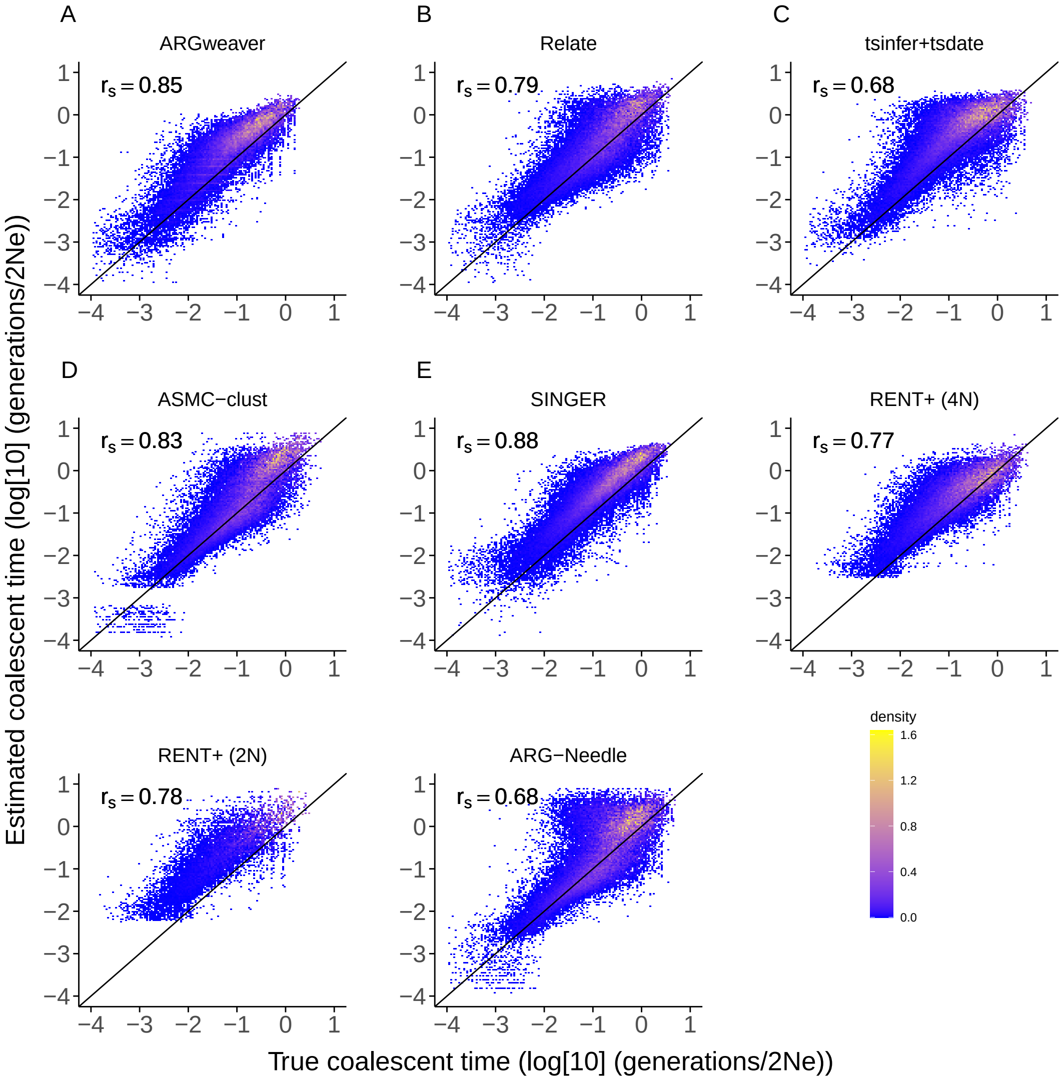


Figure S24: Analogous to Figure 5 in the main text, we show the comparison of pairwise coalescence times from true trees and estimated trees under neutrality. We also include `RENT+` (in units of $2N$ and $4N$ generations) and `ARG-Needle`. Both of them show overestimation of coalescence times—in the case of `ARG-Needle`, particularly overestimation of intermediate-to-long coalescence times. The method for sampling pairwise coalescence times is the same as under selection.

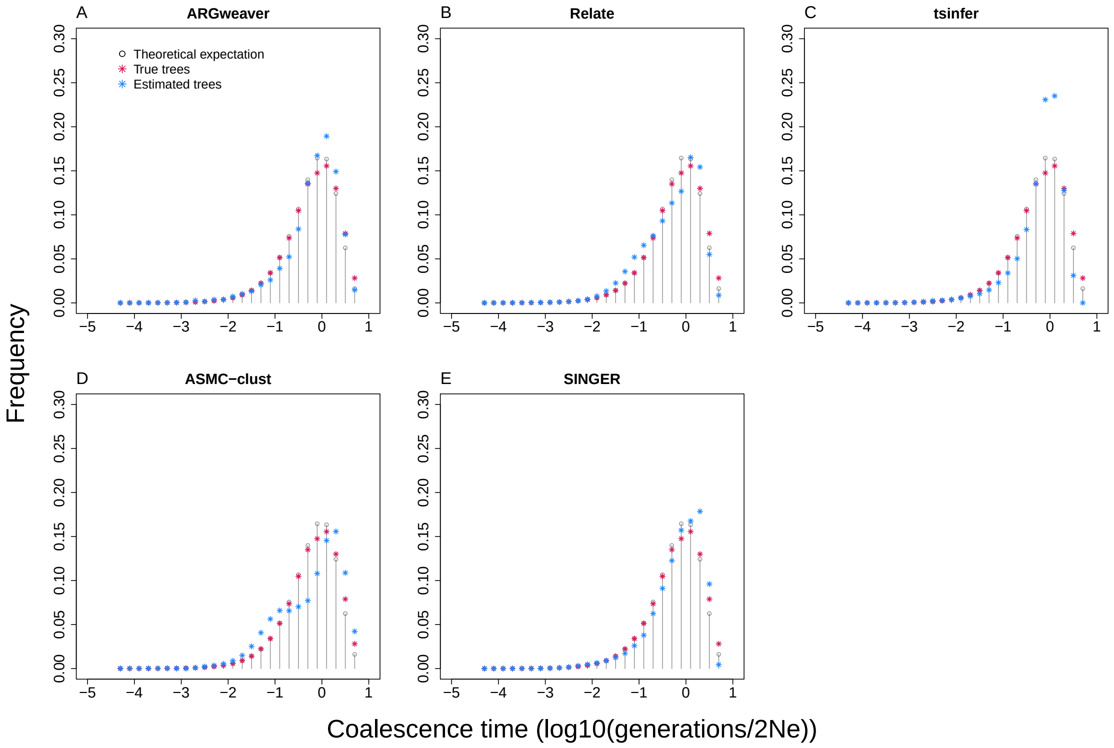


Figure S25: Distribution of the log-scaled pairwise coalescence times from the estimated trees under neutral evolution. The true trees' pairwise times differ somewhat from the theoretical expectation because we condition on a pre-defined allele frequency trajectory for an allele that is common in the present at every locus.

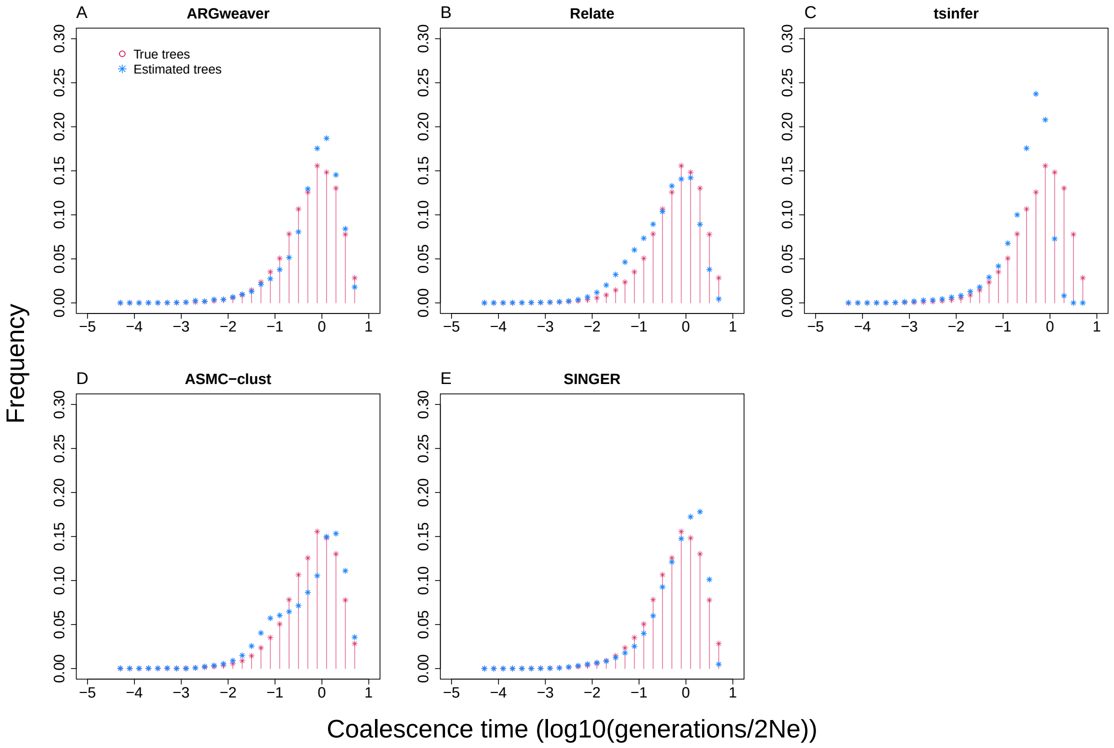


Figure S26: Distribution of the log-scaled pairwise coalescence times from the estimated trees under directional selection. Because selection changes the marginal trees, the distribution is not expected to follow the theoretical exponential distribution, which is no longer shown.

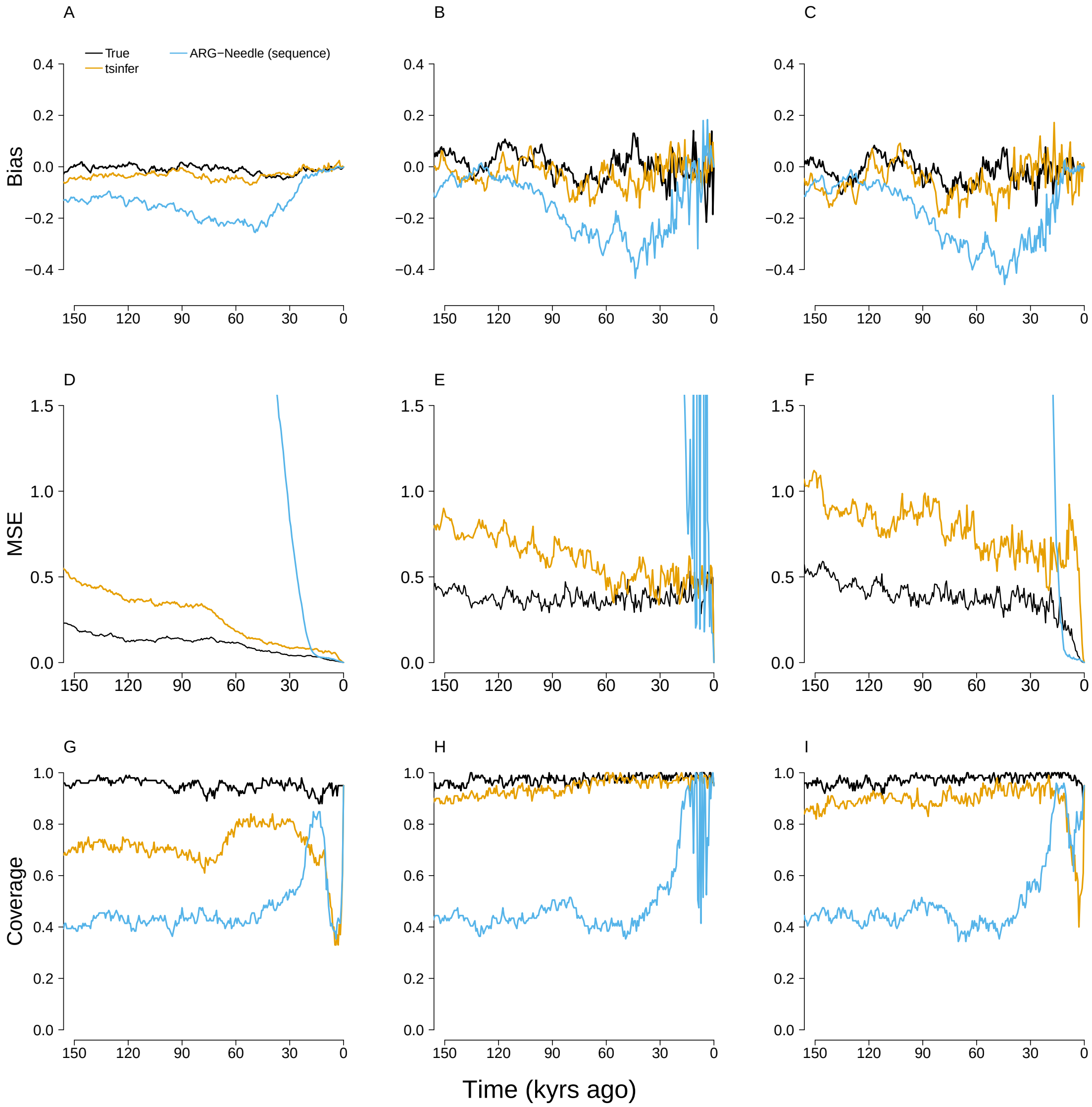


Figure S27: Performance of the methods under neutrality with samples of 5,000 chromosomes. Analogous to Figure 3 in the main text, we show the bias (A-C), MSE (D-F), and confidence-interval coverage (G-I) of the proportion-of-lineages (left column), waiting-time (middle column), and lineages-remaining estimators (right column), with the true trees and estimated trees from each ARG-estimation method as input under neutral evolution. Here, all methods are run with samples of 5,000 chromosomes. In each simulation, the PGS was formed from 100 loci and evolved neutrally. For each method, 100 simulations were performed.

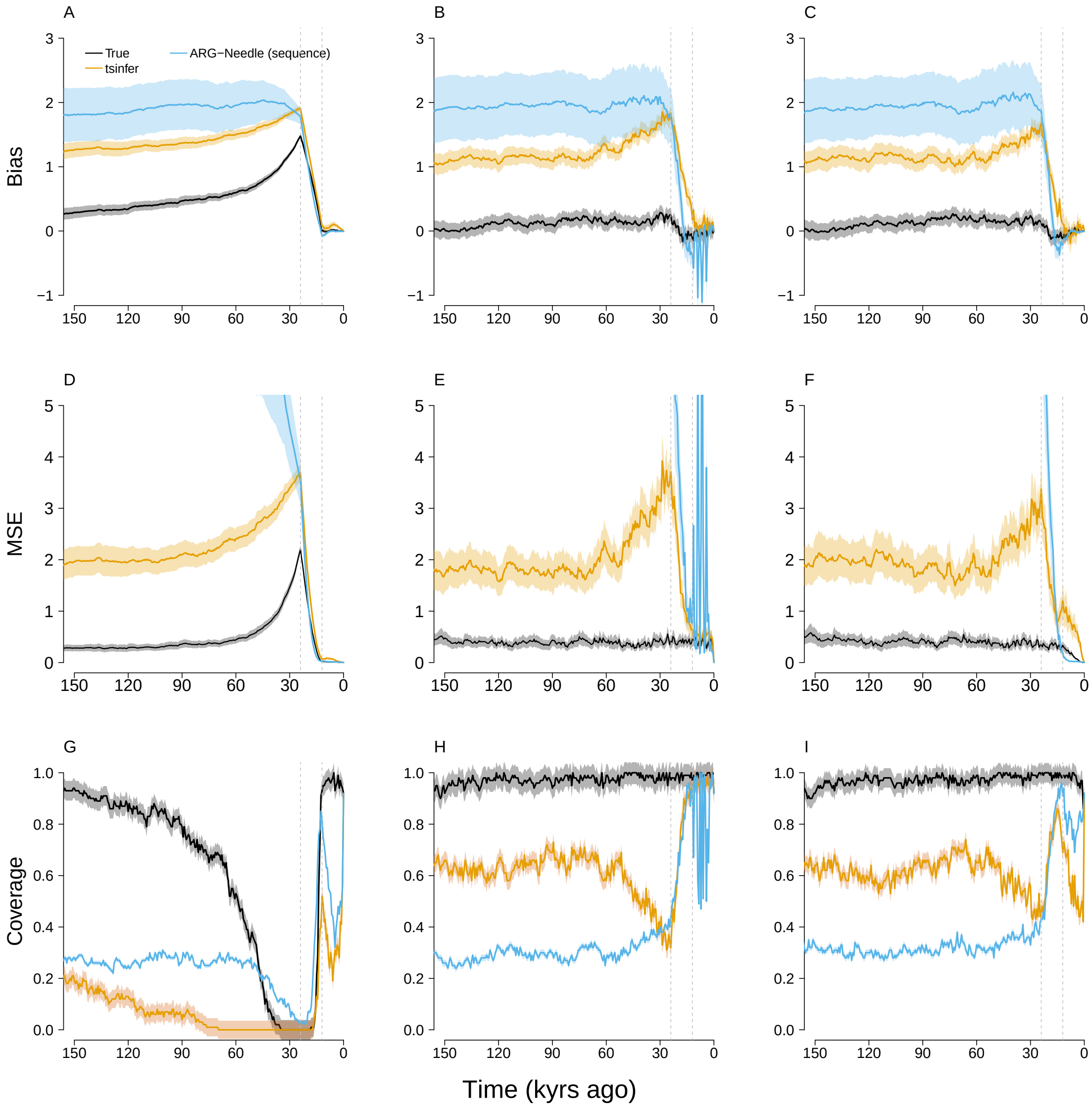


Figure S28: Performance of the methods under selection with samples of 5,000 chromosomes. Analogous to Figure 3 in the main text, we show the bias (A-C), MSE (D-F), and confidence-interval coverage (G-I) of the proportion-of-lineages (left column), waiting-time (middle column), and lineages-remaining estimators (right column), with the true trees and estimated trees from each ARG-estimation method as input under neutral evolution. Here, all methods are run with samples of 5,000 chromosomes. In each simulation, the PGS was formed from 100 loci and evolved under directional selection. For each method, 100 simulations were performed.

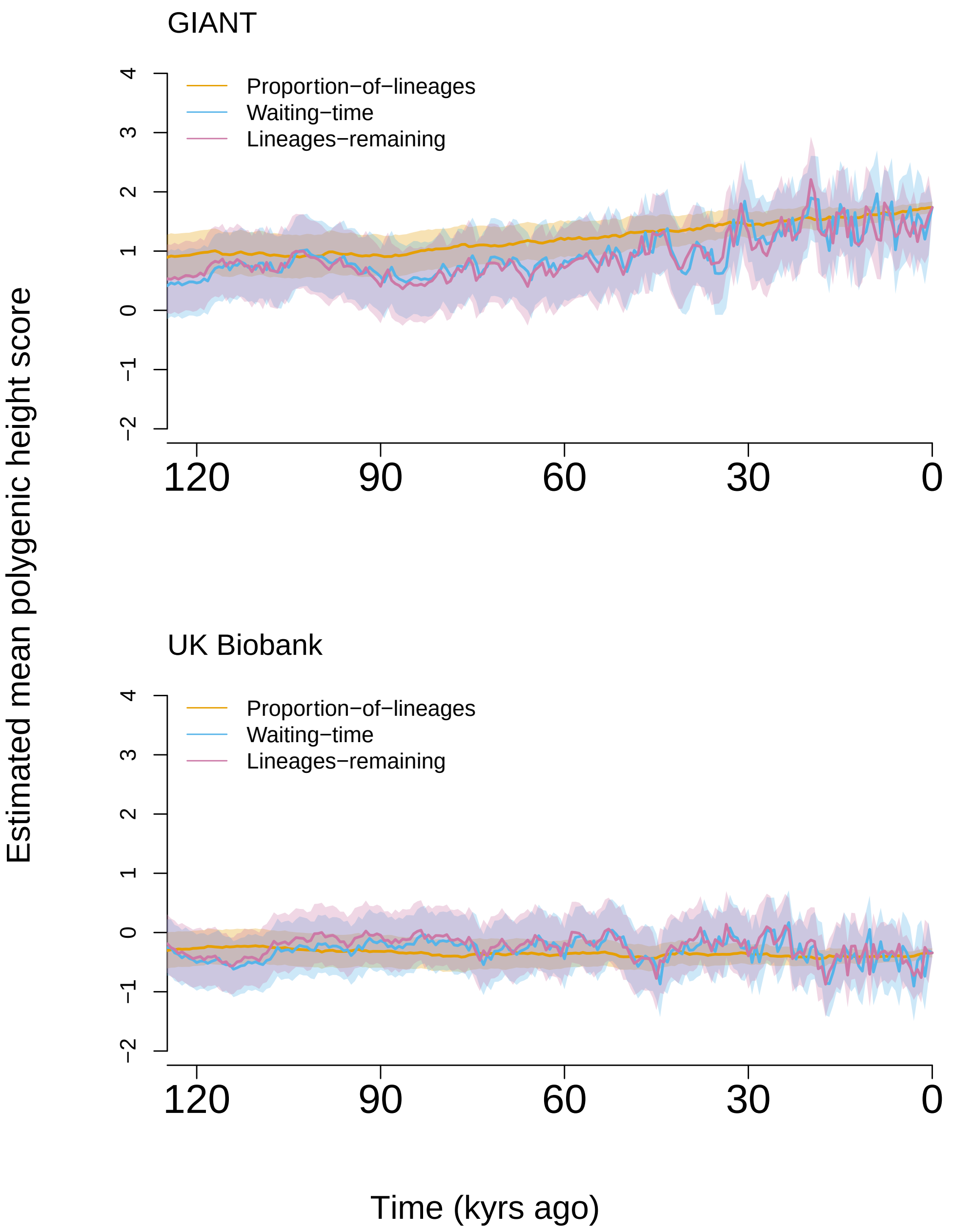


Figure S29: Estimated historical trajectories for population-mean height PGS in the ancestors of the GBR subset of the 1000 Genomes project, where local trees are estimated by `Relate`. See section S9 for details.

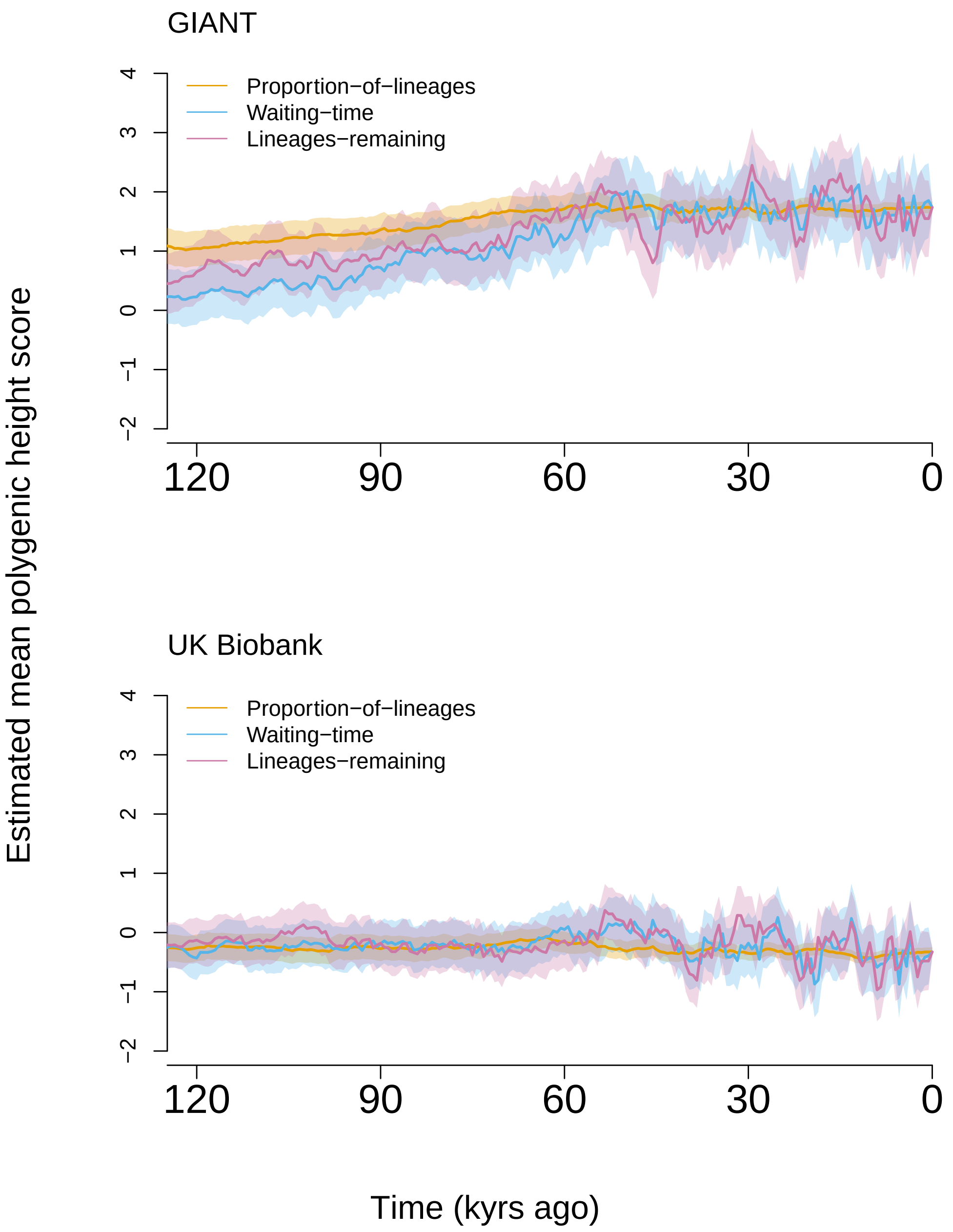


Figure S30: Estimated historical trajectories for population-mean height PGS in the ancestors of the GBR subset of the 1000 Genomes project, where local trees are estimated by `tsinfer+tsdate`. See section S9 for details.

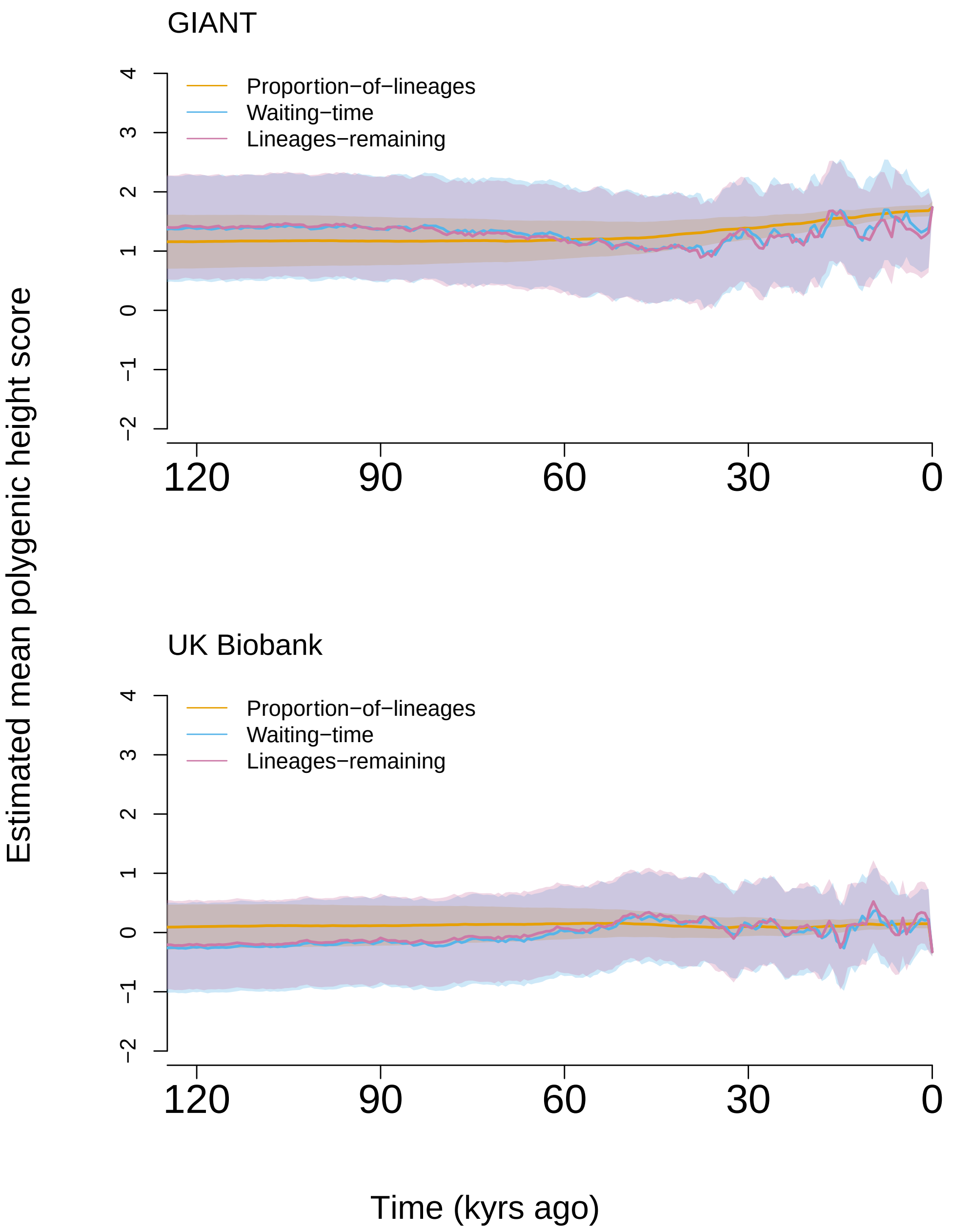


Figure S31: Estimated historical trajectories for population-mean height PGS in the ancestors of the GBR subset of the 1000 Genomes project, where local trees are estimated by `SINGER`. See section S9 for details.

Deng, Yun, Rasmus Nielsen, and Yun S Song (Mar. 2024). “Robust and accurate Bayesian inference of genome-wide genealogies for large samples”.

Edge, Michael D and Graham Coop (Jan. 2019). “Reconstructing the History of Polygenic Scores Using Coalescent Trees”. en. In: *Genetics* 211.1, pp. 235–262.

Falconer, Douglas Scott (1996). *Introduction to quantitative genetics*. Pearson Education India.

Ho, Simon Y W and Beth Shapiro (May 2011). “Skyline-plot methods for estimating demographic history from nucleotide sequences”. en. In: *Mol. Ecol. Resour.* 11.3, pp. 423–434.

Hudson, Richard R (Feb. 2002). “Generating samples under a Wright-Fisher neutral model of genetic variation”. en. In: *Bioinformatics* 18.2, pp. 337–338.

Hudson, Richard R and Norman L Kaplan (1988). “The coalescent process in models with selection and recombination.” In: *Genetics* 120.3, pp. 831–840.

Kelleher, Jerome, Alison M Etheridge, and Gilean McVean (May 2016). “Efficient Coalescent Simulation and Genealogical Analysis for Large Sample Sizes”. en. In: *PLoS Comput. Biol.* 12.5, e1004842.

Kelleher, Jerome, Yan Wong, et al. (Sept. 2019). “Inferring whole-genome histories in large population datasets”. en. In: *Nat. Genet.* 51.9, pp. 1330–1338.

Lee, Kristin M and Graham Coop (2017). “Distinguishing among modes of convergent adaptation using population genomic data”. In: *Genetics* 207.4, pp. 1591–1619.

— (2019). “Population genomics perspectives on convergent adaptation”. In: *Philosophical Transactions of the Royal Society B* 374.1777, p. 20180236.

Li, Heng (2011). “Tabix: fast retrieval of sequence features from generic TAB-delimited files”. In: *Bioinformatics* 27.5, pp. 718–719.

Li, Na and Matthew Stephens (Dec. 2003). “Modeling linkage disequilibrium and identifying recombination hotspots using single-nucleotide polymorphism data”. en. In: *Genetics* 165.4, pp. 2213–2233.

McVean, Gilean A T and Niall J Cardin (July 2005). “Approximating the coalescent with recombination”. en. In: *Philos. Trans. R. Soc. Lond. B Biol. Sci.* 360.1459, pp. 1387–1393.

Mirzaei, Sajad and Yufeng Wu (Apr. 2017). “RENT+: an improved method for inferring local genealogical trees from haplotypes with recombination”. en. In: *Bioinformatics* 33.7, pp. 1021–1030.

Neale Lab (2017). *Rapid GWAS of thousands of phenotypes for 337,000 samples in the UK Biobank*. `https://www.nealelab.is/blog/2017/7/19/rapid-gwas-of-thousands-of-phenotypes-for-337000-samples-in-the-uk-biobank`. Accessed: 2024-12-14.

Palamara, Pier Francesco et al. (Sept. 2018). “High-throughput inference of pairwise coalescence times identifies signals of selection and enriched disease heritability”. en. In: *Nat. Genet.* 50.9, pp. 1311–1317.

Pybus, O G, A Rambaut, and P H Harvey (July 2000). “An integrated framework for the inference of viral population history from reconstructed genealogies”. en. In: *Genetics* 155.3, pp. 1429–1437.

Racimo, Fernando, Jeremy J Berg, and Joseph K Pickrell (2018). “Detecting polygenic adaptation in admixture graphs”. In: *Genetics* 208.4, pp. 1565–1584.

Rasmussen, Matthew D et al. (May 2014). “Genome-wide inference of ancestral recombination graphs”. en. In: *PLoS Genet.* 10.5, e1004342.

Sokal, Robert R and Charles D Michener (1958). “A statistical method for evaluating systematic relationships”. In.

Speidel, Leo et al. (Sept. 2019). “A method for genome-wide genealogy estimation for thousands of samples”. en. In: *Nat. Genet.* 51.9, pp. 1321–1329.

Spence, Jeffrey P and Yun S Song (2019). “Inference and analysis of population-specific fine-scale recombination maps across 26 diverse human populations”. In: *Science Advances* 5.10, eaaw9206.

Stern, Aaron J et al. (Feb. 2021). “Disentangling selection on genetically correlated polygenic traits via whole-genome genealogies”. en. In: *Am. J. Hum. Genet.* 108.2, pp. 219–239.

Stoler, Nicholas and Anton Nekrutenko (2021). “Sequencing error profiles of Illumina sequencing instruments”. In: *NAR genomics and bioinformatics* 3.1, lqab019.

Terhorst, Jonathan, John A Kamm, and Yun S Song (2017). “Robust and scalable inference of population history from hundreds of unphased whole genomes”. In: *Nature genetics* 49.2, pp. 303–309.

Walsh, Bruce and Michael Lynch (2018). *Evolution and selection of quantitative traits*. Oxford University Press.

Wohns, Anthony Wilder et al. (Feb. 2022). “A unified genealogy of modern and ancient genomes”. en. In: *Science* 375.6583, eabi8264.

Wood, Andrew R et al. (2014). “Defining the role of common variation in the genomic and biological architecture of adult human height”. In: *Nature genetics* 46.11, pp. 1173–1186.

Wu, Yufeng (2011). “New methods for inference of local tree topologies with recombinant SNP sequences in populations”. en. In: *IEEE/ACM Trans. Comput. Biol. Bioinform.* 8.1, pp. 182–193.

Zhang, Brian C et al. (May 2023). “Biobank-scale inference of ancestral recombination graphs enables genealogical analysis of complex traits”. en. In: *Nat. Genet.* 55.5, pp. 768–776.
